# *Klebsiella pneumoniae* hijacks the Toll-IL-1R protein SARM1 in a type I IFN-dependent manner to antagonize host immunity

**DOI:** 10.1101/2021.09.29.462388

**Authors:** Claudia Feriotti, Joana Sa-Pessoa, Ricardo Calderón-González, Lili Gu, Brenda Morris, Ryoichi Sugisawa, Jose L. Insua, Michael Carty, Amy Dumigan, Rebecca J. Ingram, Adrien Kisenpfening, Andrew G. Bowie, José A. Bengoechea

## Abstract

Many bacterial pathogens antagonize host defence responses by translocating effector proteins into cells. It remains an open question how those pathogens not encoding effectors counteract anti-bacterial immunity. Here, we show that *Klebsiella pneumoniae* hijacks the evolutionary conserved innate immune protein SARM1 to control cell intrinsic immunity. *Klebsiella* exploits SARM1 to regulate negatively MyD88 and TRIF-governed inflammation, and the activation of the MAP kinases ERK and JNK. SARM1 is required for *Klebsiella* induction of IL10 by fine-tuning the p38-type I IFN axis. SARM1 inhibits the activation of *Klebsiella*-induced absent in melanoma 2 inflammasome to limit IL1β production, suppressing further inflammation. *Klebsiella* exploits type I IFNs to induce SARM1 in a capsule and LPS O-polysaccharide-dependent manner via TLR4-TRAM-TRIF-IRF3-IFNAR1 pathway. Absence of SARM1 reduces the intracellular survival of *K. pneumonaie* in macrophages whereas *sarm1* deficient mice control the infection. Altogether, our results illustrate a hitherto unknown anti-immunology strategy deployed by a human pathogen.

## INTRODUCTION

*Klebsiella pneumoniae* is one of the pathogens sweeping the World in the antimicrobial resistance pandemic. More than a third of the *K. pneumoniae* isolates reported to the European Centre for Disease Prevention and Control were resistant to at least one antimicrobial group, being the most common resistance phenotype the combined resistance to fluoroquinolones, third-generation cephalosporins and aminoglycosides (Penalva et al., 2019). In addition, *Klebsiella* species are a known reservoir for antibiotic resistant genes, which can spread to other Gram-negative bacteria. Infections caused by multidrug resistant *K. pneumoniae* are associated with high mortality rates and prolonged hospitalization (Giske et al., 2008). Alarmingly, recent studies have also recognised that *K. pneumoniae* strains have access to a mobile pool of virulence genes (Holt et al., 2015; Lam et al., 2018); enabling the emergence of a multidrug, hypervirulent *K. pneumoniae* clone capable of causing untreatable infections in healthy individuals. Worryingly, there are already reports describing the isolation of such strains (Gu et al., 2018; Yao et al., 2018; Zhang et al., 2015; Zhang et al., 2016). Unfortunately, at present, we cannot identify candidate compounds in late-stage development for treatment of multidrug *K. pneumoniae* infections. This pathogen is exemplary of the mismatch between unmet medical needs and the current antimicrobial research and development pipeline.

An attractive approach to develop new therapeutics against *K. pneumoniae* infections is to boost innate immune defence mechanisms. Indeed, more than two decades of research demonstrates the need of an adequate activation of the innate immune system for the clearance of *K. pneumoniae* (Bengoechea and Sa Pessoa, 2019). However, this pathway requires an in-depth understanding of which innate responses benefit the host versus the pathogen, as well as deconstructing the strategies used by *K. pneumoniae* to survive within the infected tissue. In this regard, the fact that *K. pneumoniae* does not encode type III or IV secretion systems known to deliver effectors into immune cells, or any of the toxins affecting cell biology, makes interesting to uncover how the pathogen controls the activation of immune cells.

Successful elimination of infections by the innate immune system is dependent on the activation of pattern recognition receptors (PRRs) detecting the so-called pathogen-associated molecular patterns (PAMPs). The PRRs Toll-like receptor (TLR) 4 and TLR2 play a significant role restricting *K. pneumoniae* infection (Wieland et al., 2011). TLR4 and TLR2 signal via the adaptors MyD88 and TRIF leading to the activation of NF-κB and IRF3, respectively. These transcription factors and MAP kinases control the activation of host defence antimicrobial responses (Jenner and Young, 2005). The fact that *IL1R^-/-^* mice are exquisitely susceptible to *K. pneumoniae* infection demonstrates the importance of IL1β-controlled responses for host survival and bacterial clearance (Cai et al., 2012). Production of the mature active form of IL1β requires the expression of the pro- IL1β, following PRR-mediated recognition of a pathogen, and its cleavage by caspase 1 to release the active form of the cytokine. The activation of caspase 1 also leads to pyroptosis through the proteolytic cleavage of gasdermin-D (GSDMD). The activation of caspase 1 requires the assembly of a multiprotein platform, known as an inflammasome. Evidence suggests that *K. pneumoniae* induces the secretion of IL1β in vivo and in vitro via inflammasome activation (Cai et al., 2012; Willingham et al., 2009b). Whether *K. pneumoniae* has evolved any strategy to limit early events of TLR signalling, and inflammasome activation remains an open question.

SARM1 (Sterile α and HEAT Armadillo motif-containing protein) is an evolutionary conserved innate immune protein across mammalian species with identities higher than 90% (Belinda et al., 2008). Moreover, analysis of human SARM1 has revealed no nonsense mutations and a worldwide selective sweep, indicating a strong selective pressure to preserve the integrity of the protein (Fornarino et al., 2011). SARM1 contains a Toll-IL-1R (TIR) domain (Bratkowski et al., 2020; O’Neill and Bowie, 2007). The presence of this domain indicates a role in IL1 and TLR signalling. Interestingly, bacterial proteins containing the TIR domain interfere with TLR signalling to inhibit innate immune responses (Askarian et al., 2014; Cirl et al., 2008; Coronas-Serna et al., 2020; Imbert et al., 2017; Xiong et al., 2019). It is intriguing that the SARM1 TIR domain is more closely related to bacteria TIR proteins than to the other mammalian TIR containing adaptors (Zhang et al., 2011). Therefore, it can be speculated that SARM1 may play a negative role regulating TLR signalling. Indeed, there is data suggesting that SARM1 inhibits lipopolysaccharide (LPS)-induced signalling via TLR4-TRIF and TLR4-MyD88 pathways (Carlsson et al., 2016; Carty et al., 2006). Furthermore, recent work has uncovered that SARM1 negatively regulates IL1β release by directly targeting the NLRP3 inflammasome (Carty et al., 2019). Collectively, this evidence led us to speculate whether *K. pneumoniae* may hijack SARM1, an endogenous TIR-containing protein regulating TLR and inflammasome activation, to control immune responses. The role of SARM1 in infections has been only conclusively established to restrict West Nile virus infection in the central nervous system (Szretter et al., 2009; Uccellini et al., 2020). *Sarm1^-/-^* mice do not control West Nile virus infection, and this is associated with enhanced mortality (Szretter et al., 2009; Uccellini et al., 2020). To the best of our knowledge, there is no evidence supporting any role of SARM1 in bacterial infections.

Here, we reveal that hypervirulent *K. pneumoniae* leverages the immunomodulatory roles of SARM1 to control cell intrinsic immunity. We show that *K. pneumoniae* negatively regulates TLR-governed inflammatory responses via SARM1. We demonstrate that SARM1 is required for *K. pneumoniae*-induction of the anti-inflammatory cytokine IL10. We identify absent in melanoma 2 (AIM2) as the inflammasome activated by *K. pneumoniae* that is inhibited directly by SARM1 to limit IL1β production. We establish that *K. pneumoniae* exploits the immune effector type I IFNs to induce SARM1 in a capsule and LPS O-polysaccharide-dependent manner. In vitro, absence of SARM1 reduces the intracellular survival of *K. pneumoniae* in macrophages due to the recruitment of lysosomes to the *Klebsiella* containing vacuole (KCV), whereas, in vivo, *Sarm1^-/-^* mice clear the infection. Collectively, our findings illustrate the crucial role of SARM1 in *K. pneumoniae* immune evasion strategies, revealing one of the Achilles heel of our immune system exploited by the pathogen to overcome host protective responses.

## RESULTS

### SARM1 negatively regulates *K. pneumoniae*-induced inflammation

To examine the effect of SARM1 on *K. pneumoniae*-induced responses, we infected immortalized bone marrow derived macrophages (iBMDMs) from wild-type and *sarm1^-/-^* mice with the hypervirulent strain of *K. pneumoniae* CIP52.145 (hereafter Kp52145). This strain belongs to the *K. pneumoniae* KpI group and it encodes all virulence functions associated with invasive community-acquired disease in humans (Holt et al., 2015; Lery et al., 2014). In the supernatants of cells lacking SARM1, we observed a significant increase in the levels of the MyD88-dependent cytokines TNFα, and IL1β, and of the TRIF-dependent cytokines CXCL10 and type I IFNs following Kp52145 infection. (Fig 1A). The levels of the TRIF-dependent proteins ISG15 and Viperin were also higher in the lysates of Kp52145-infected *sarm1^-/-^* macrophages than in those of wild-type cells (Fig 1B). Kp52145 also increased the levels of the MyD88-dependent cytokines TNFα, and IL1β, and of the TRIF-dependent cytokine CXCL10 in BMDMs from *sarm1* mice (Fig S1A), ruling out that the heightened responses observed in inmortalized *sarm1^-/-^* cells were due to the process of immortalization of the cells. To confirm that the phenotype of *sarm1^-/-^* cells was due to the absence of the SARM protein, rescue experiments were performed by retroviral expression of FLAG-SARM1 in *sarm1^-/-^* iBMDMs. Following infection with Kp52145, we observed a reduction in the levels of TNFα, IL1β, and CXCL10 in FLAG SARM1 cells compare to those found in infected *sarm1^-/-^* macrophages (Fig 1C). Collectively, these data demonstrate that SARM1 negatively regulates *K. pneumoniae*-induced inflammation.

**Figure 1.**
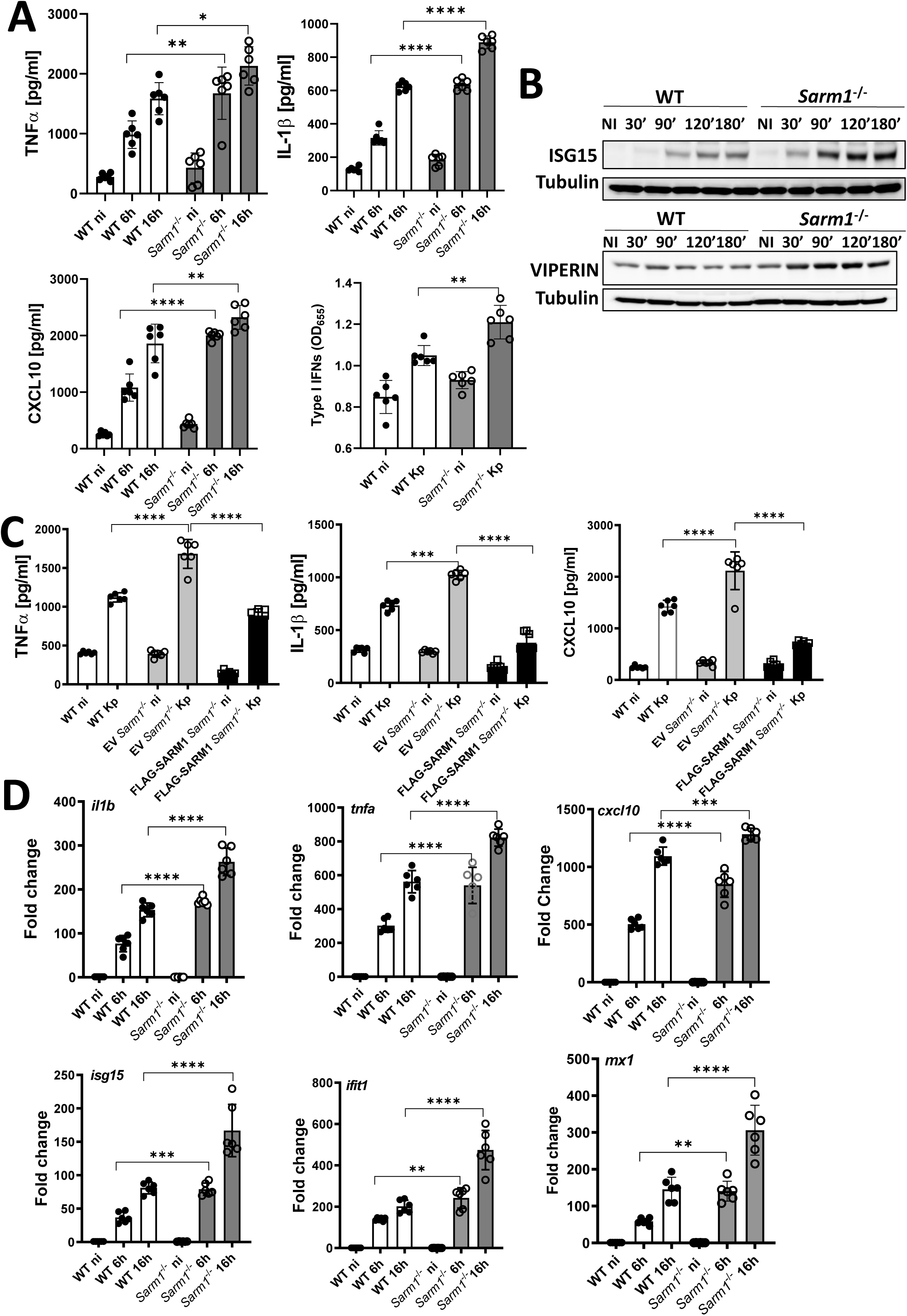
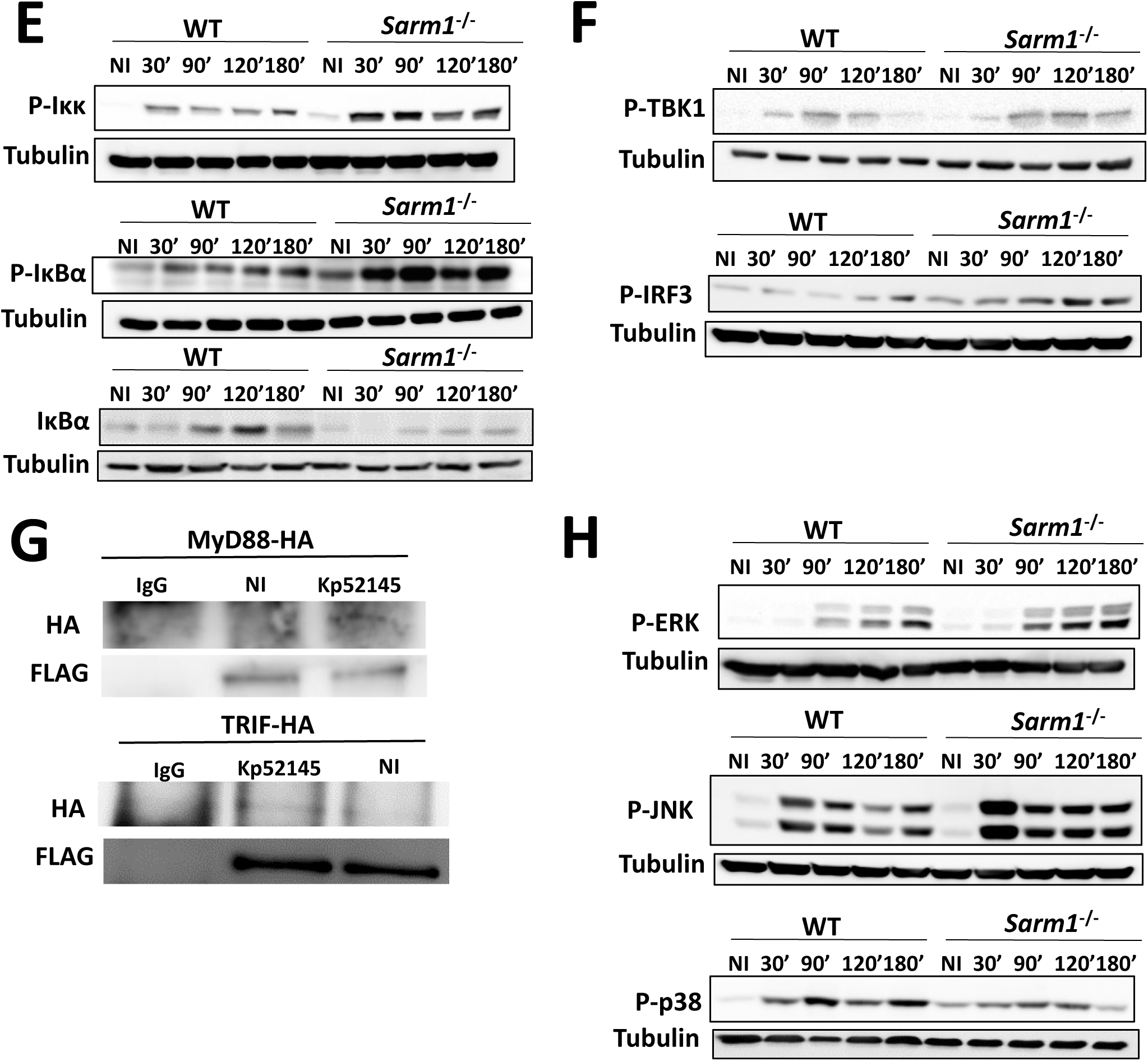
SARM1 negatively regulates *K. pneumoniae*-induced inflammation. A. ELISA of TNFα, IL1β, CXCL10 secreted by wild-type (WT) and *sarm1^-/-^* macrophages non-infected (ni) or infected with Kp52145 for 6 and 16 h. Type I IFN levels determined in the supernatants of macrophages 16 h post infection. The reporter cell line B16-Blue IFN-α/β was used for the quantification of levels of SEAP produced upon stimulation of the supernatants with the detection medium QUANTI-Blue and presented as OD_655_. After 1 h contact, the medium was replaced with medium containing gentamicin (100 µg/ml) to kill extracellular bacteria. B. Immunoblot analysis of ISG15, Viperin and tubulin levels in lysates of wild-type (WT) and *sarm1^-/-^* macrophages non-infected (NI) or infected with Kp52145 for the indicated time points. After 1 h contact, the medium was replaced with medium containing gentamicin (100 µg/ml) to kill extracellular bacteria. C. ELISA of TNFα , IL1β, CXCL10 secreted by wild-type (WT) macrophages, and retrovirally transfected *sarm1^-/-^* cells with FLAG-SARM1 or control vector (EV) non-infected (ni) or infected with Kp52145 (Kp) for 16 h. After 1 h contact, the medium was replaced with medium containing gentamicin (100 µg/ml) to kill extracellular bacteria. D. *il1b, tnfa, cxcl10, isg15, ifit1*, and *mx1* mRNA levels were assessed by qPCR, in wild-type (WT) and *sarm1^-/-^* macrophages non-infected (ni) or infected with Kp52145 for 6 and 16 h. After 1 h contact, the medium was replaced with medium containing gentamicin (100 µg/ml) to kill extracellular bacteria. E. Immunoblot analysis of phosphorylated Iκκα/β (P-Iκκ), phosphorylated IκBα (P-IκBα), total IκBα (IκBα) and tubulin levels in lysates of wild-type (WT) and *sarm1-/-* macrophages non-infected (NI) or infected with Kp52145 for the indicated time points. After 1 h contact, the medium was replaced with medium containing gentamicin (100 μg/ml) to kill extracellular bacteria. F. Immunoblot analysis of phosphorylated TBK1 (P-TBK1), phosphorylated Irf3 (P-IRF3) and tubulin levels in lysates of wild-type (WT) and *sarm1^-/-^* macrophages non-infected (NI) or infected with Kp52145 for the indicated time points. After 1 h contact, the medium was replaced with medium containing gentamicin (100 µg/ml) to kill extracellular bacteria. G. *Sarm1^FLAG^* macrophages were transfected with a MyD88-HA or TRIF-HA plasmids, and the following day infected with Kp52145. Cells were lysed in RIPA buffer, and lysates immunoprecipitated using anti-FLAG antibody. Preimmune mouse IgG served as negative control. H. Immunoblot analysis of phosphorylated ERK (P-ERK), phosphorylated JNK (P-JNK phosphorylated p38 (P-p38) and tubulin levels in lysates of wild-type (WT) and *sarm1^-/-^* macrophages non-infected (NI) or infected with Kp52145 for the indicated time points. In panels B, E, F, G, and H images are representative of three independent experiments. In panels A, C and D values are presented as the mean ± SD of three independent experiments measured in duplicate. ****P ≤ 0.0001; ***P ≤ 0.001; ** P ≤ 0.01; *P ≤ 0.05 for the indicated comparisons determined using one way-ANOVA with Bonferroni contrast for multiple comparisons test.

Recently, it has been reported that commercially available *sarm1^-/-^* mice carry a passenger mutation which may affect cytokine responses due to the background of the knockout strains and not due to the absence of SARM1 protein (Uccellini et al., 2020). Although the cytokines affected are not those assessed in our study, we decided to examine the role of SARM1 in *K. pneumoniae* infection by reducing its levels via siRNA. Control experiments showed the reduction of the *Sarm1* transcript in transfected iBMDMs with SARM siRNA (Fig S1B), confirming knockdown. We again found higher levels of IL1β, TNFα, and CXCL10 in the supernatants of infected *sarm1* knockdown macrophages that in macrophages transfected with a non-targeting (All Stars) siRNA control (Fig S1C). To provide additional support to our observations demonstrating a role of SARM1 as negative regulator of *K. pneumoniae*-induced inflammation, we challenged iBMDMs from a recently described new knockout SARM1 strain generated using CRISPR/Cas9-mediated genome engineering, *Sarm1^em1.1Tft^* (Doran et al., 2021). Kp52145 induced a heightened inflammatory response in *Sarm1^em1.1Tft^* macrophages compared to wild-type ones from littermates (Fig S1D). Altogether, these data provide further evidence supporting the role of SARM1 to attenuate *K. pneumoniae*-induced inflammation.

To determine whether the observed changes in cytokine production in the absence of SARM1 involved changes in the transcription of genes, we assessed the transcription of MyD88 and TRIF-dependent cytokines by real time quantitative PCR (RT-qPCR). Figure 1D shows that Kp52145 increased the transcription of the MyD88-governed cytokines *tnfa, il1b,* and of the TRIF- controlled cytokine *ifnb* in *sarm1^-/-^* macrophages compare to wild-type ones. The transcription of the interferon-stimulated genes (ISG) *isg15*, *mx1* and *ifit1* was also upregulated in infected *sarm1^-/-^* cells (Fig 1D).

The fact that the transcription factors NF-κB and IRF3 governs the MyD88 and TRIF-dependent responses, respectively led us to determine whether SARM1 regulates these pathways in *K. pneumoniae*-infected cells. In the NF-κB signalling cascade, the IKKα/β kinase controls the phosphorylation of IκBα that leads to the subsequent degradation of the protein by the ubiquitin proteasome, allowing the nuclear translocation of NF- κB (Taniguchi and Karin, 2018). Immunoblotting analysis showed an increase in the phosphorylation of IKKα/β in *sarm1* macrophages following infection (Fig 1E). As expected, we observed an increased phosphorylation of IκBα with a concomitant reduction in the levels of total I κBα in Kp52145-infected *sarm1^-/-^* macrophages compare to wild-type ones (Fig 1E). Altogether, these results show an enhance activation of NF-κB in infected *sarm1^-/-^* macrophages. To investigate the activation of the IRF3 signalling cascade, we assessed the phosphorylation of TBK1 and IRF3. TBK1 is the kinase mediating the phosphorylation of IRF3 which it is an essential event for IRF3 nuclear translocation (Fitzgerald et al., 2003). Immunoblotting experiments revealed an increased phosphorylation of TBK1 and IRF3 in Kp52145-infected *sarm1^-/-^* macrophages (Fig 1F), hence confirming an increased activation of IRF3 in the absence of SARM1.

Reconstitution experiments in HEK293 cells by transfecting SARM1, and either MyD88 or TRIF, and reporter systems to assess activation of NF- κB and IRF3 have demonstrated that SARM1 interacts with MyD88 and TRIF to block the activation of these signalling pathways (Carlsson et al., 2016; Carty et al., 2006). Therefore, we sought to determine whether *K. pneumoniae* infection would induce the interaction between SARM1, and MyD88, and TRIF. There is no commercially available antibody to assess mouse SARM1 protein levels reliably. Therefore, to facilitate these experiments, we took advantage of a recently described mouse expressing an epitope-tagged SARM1 endogenously with a triple FLAG tag and double strep tag on the C-terminal end, *Sarm1^FLAG^* (Doran et al., 2021). Control experiments confirmed that the tagged protein retain functionality (Doran et al., 2021). Figure 1G shows that in *Sarm1^FLAG^* iBMDMs SARM1-FLAG co-immunoprecipitates MyD88-HA and TRIF-HA only in Kp52145-infected cells, indicating that *K. pneumoniae*-induced interaction of SARM1 with MyD88 and TRIF explains the reduced activation of NF-κB and IRF3.

We next assessed the activation of MAPKs due to their role in governing the expression of inflammatory genes (Dong et al., 2002). There is indirect evidence suggesting that SARM1 inhibits MAPK activation (Peng et al., 2010). The activation of the three MAPKs p38, JNK and ERK occurs through phosphorylation of serine and threonine residues. Western blot analysis showed an increase in the levels of phosphorylated ERK and JNK in infected *sarm1^-/-^* macrophages compared to infected wild-type cells (Fig 1H). In contrast, there was a reduction in the phosphorylation of p38 in infected *sarm1^-/-^* macrophages (Fig 1H). These results indicate that SARM1 exerts a negative effect on the activation of ERK and JNK whereas SARM1 is needed for the activation of p38 following *K. pneumoniae* infection.

### SARM1 is required for *K. pneumoniae* induction of IL10 via p38

We next sought to determine the effect of the reduced activation of p38 in the absence of SARM1 following *K. pneumoniae* infection. Because the activation of p38 is linked to the production of IL10 (Saraiva and O’Garra, 2010), we asked whether SARM1 would affect *K. pneumoniae* induction of IL10. Control experiments confirmed that the p38 inhibitor SB203580 abrogated Kp52145-induced production of IL10 in wild-type cells (Fig S2A), connecting p38 activation and IL10 production in *K. pneumoniae*-infected macrophages. Consistent with the reduced activation of p38 in *sarm1^-/-^* macrophages, RT-qPCR analysis showed a reduction in *il10* transcription in the absence of SARM1 (Fig 2A). As expected, the levels of IL10 were significantly lower in the supernatants of Kp52145-infected *sarm1^-/-^* macrophages than in those from infected wild-type cells (Fig 2B). The reduced levels of IL10 found in infected *sarm1^-/-^* macrophages were consistent with the reduced phosphorylation of the IL10-governed transcriptional factor STAT3 in Kp52145-infected *sarm1^-/-^* macrophages (Fig 2C). The addition of recombinant IL10 to Kp52145-infected *sarm1^-/-^* macrophages decreased the levels of IL1β, TNFα, and CXCL10 (Fig 2D), suggesting that the reduced levels of IL10 in the absence of SARM1 contributes to the upregulation of inflammation in infected *sarm1^-/-^* macrophages. Interestingly, Kp52145 did increase the levels of *il1b*, *tnfa*, and *cxcl10* in *il10^-/-^ sarm1* knockdown macrophages beyond the levels found in *il10^-/-^* infected cells (Fig 2E), suggesting that the regulatory effect of SARM1 on inflammation is the sum of the IL10-dependent attenuation, and the direct negative effect of SARM1 on MyD88 and TRIF previously shown. The efficiency of *sarm1* knockdown in the *il10*^-/-^ background is shown in Figure S2B.

**Figure 2.**
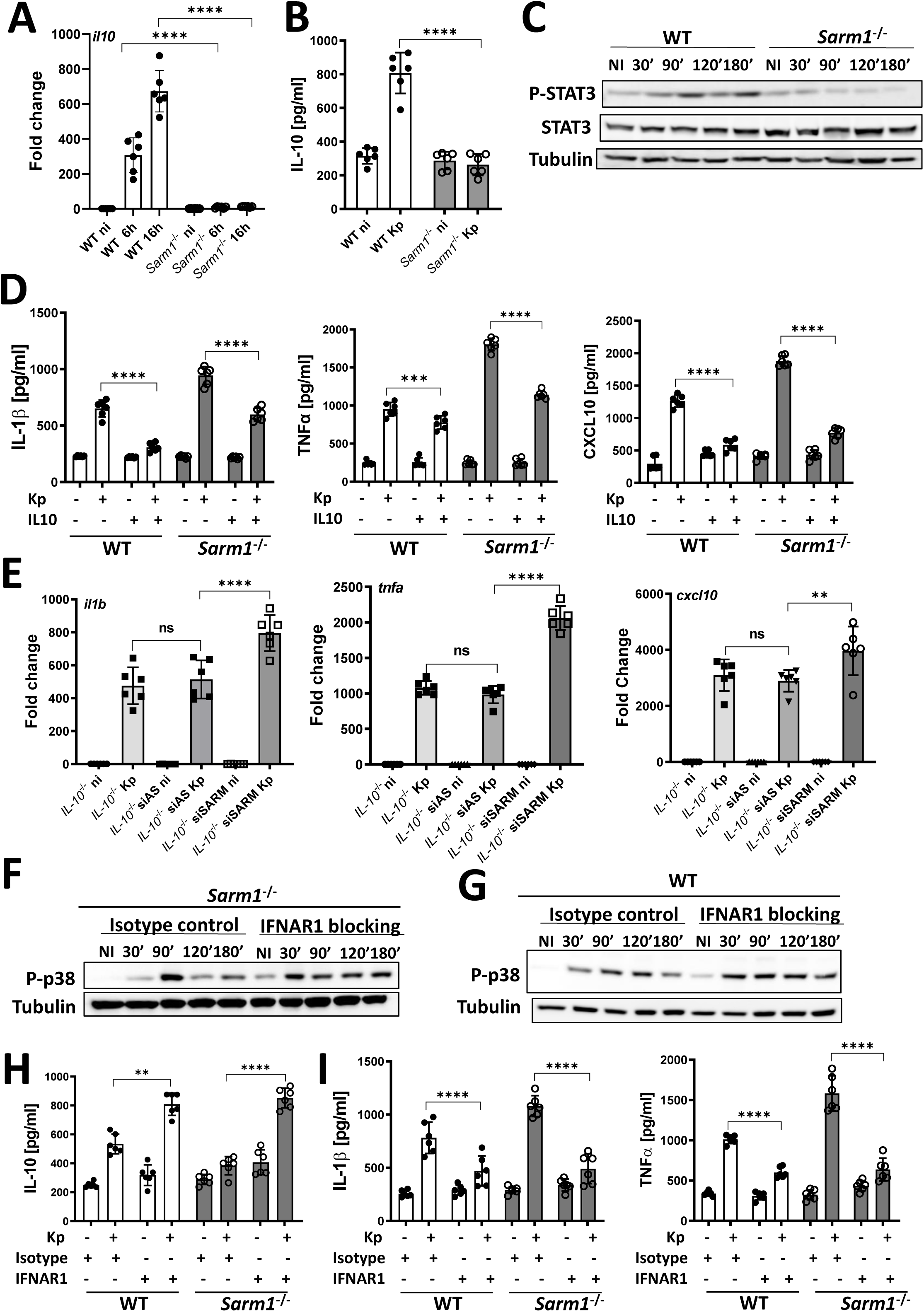
SARM1 is required for *K. pneumoniae* induction of IL10 via p38. A. *il10* mRNA levels were assessed by qPCR, in wild-type (WT) and *sarm1^-/-^* macrophages non- infected (ni) or infected with Kp52145 for 6 and 16 h. After 1 h contact, the medium was replaced with medium containing gentamicin (100 µg/ml) to kill extracellular bacteria. B. ELISA of IL10 secreted by wild-type (WT) and *sarm1^-/-^* macrophages non-infected (ni) or infected with Kp52145 for 16 h. After 1 h contact, the medium was replaced with medium containing gentamicin (100 µg/ml) to kill extracellular bacteria. C. Immunoblot analysis of phosphorylated STAT3 (P-STAT3), total STAT3 (STAT3) and tubulin levels in lysates of wild-type (WT) and *sarm1^-/-^* macrophages non-infected (NI) or infected with Kp52145 for the indicated time points. After 1 h contact, the medium was replaced with medium containing gentamicin (100 µg/ml) to kill extracellular bacteria. D. ELISA of TNFα, IL1β, CXCL10 secreted by wild-type (WT) and *sarm1^-/-^* macrophages non- infected or infected with Kp52145 for 16 h. Where indicated, cells were treated with recombinant IL10 (1 ng/ml) overnight before infection. After 1 h contact, the medium was replaced with medium containing gentamicin (100 µg/ml) to kill extracellular bacteria. E. *il1b, tnfa,* and *cxcl10* mRNA levels were assessed by qPCR in *il10^-/-^* macrophages, and *il10^-/-^* cells transfected with All Stars siRNA control (AS), or SARM1 siRNA (siSARM) non-infected (ni) or infected with Kp52145 (Kp) for 16 h. F. Immunoblot analysis of phosphorylated p38 (P-p38), and tubulin levels in lysates of *sarm1^-/-^* macrophages treated with isotype control antibody, or IFNAR1 blocking non-infected (NI) or infected with Kp52145 for the indicated time points. After 1 h contact, the medium was replaced with medium containing gentamicin (100 µg/ml) to kill extracellular bacteria. G. Immunoblot analysis of phosphorylated p38 (P-p38), and tubulin levels in lysates of wild-type macrophages (WT) treated with isotype control antibody, or IFNAR1 blocking non-infected (NI) or infected with Kp52145 for the indicated time points. After 1 h contact, the medium was replaced with medium containing gentamicin (100 µg/ml) to kill extracellular bacteria. H. ELISA of IL10, secreted by wild-type (WT) and *sarm1^-/-^* macrophages non-infected or infected with Kp52145 for 16 h. Where indicated, cells were treated with isotype control antibody, or IFNAR1 blocking overnight before infection. After 1 h contact, the medium was replaced with medium containing gentamicin (100 µg/ml) to kill extracellular bacteria. I. ELISA of IL1β, and TNFα secreted by wild-type (WT) and *sarm1^-/-^* macrophages non-infected or infected with Kp52145 for 16 h. Where indicated, cells were treated with isotype control antibody, or IFNAR1 blocking overnight before infection. After 1 h contact, the medium was replaced with medium containing gentamicin (100 µg/ml) to kill extracellular bacteria. In panels C, F, and G images are representative of three independent experiments. In panels A, B, D, E, H and I values are presented as the mean ± SD of three independent experiments measured in duplicate. ****P ≤ 0.0001; ***P ≤ 0.001; ** P ≤ 0.01; ns, P > 0.05 for the indicated comparisons determined using one way-ANOVA with Bonferroni contrast for multiple comparisons test.

To explain the reduced activation of p38 in the absence of SARM1, we reasoned that the heightened inflammation upon infection of *sarm1^-/-^* macrophages might have a negative effect on p38 activation. Because there are reports demonstrating a connection between type I IFN signalling and p38 (Ivashkiv and Donlin, 2014), we speculated that the elevated levels of type I IFNs found in Kp52145-infected *sarm1^-/-^* macrophages might underline the reduced activation of p38. To explore this possibility, we asked whether abrogating type I IFN signalling in the absence of SARM1 could rescue p38 activation following infection. Indeed, when the infection of *sarm1^-/-^* macrophages was done in the presence of blocking antibodies against the type I IFN receptor (IFNAR1) we observed an increase in the levels of phosphorylated p38 (Fig 2F). Likewise, we observed an increase in the levels of phosphorylated p38 in infected wild-type cells treated with the IFNAR1 receptor blocking antibody (Fig 2G), reinforcing the connection between type I IFN levels and *K. pneumoniae-* induced activation of p38. As anticipated, the levels of IL10 were higher in cells treated with the IFNAR1 blocking antibody than in those treated with the isotype control antibody (Fig 2H). In turn,, we found a reduction in the levels of IL1β, and TNFα in the supernatants of cells treated with the blocking antibody (Fig 2I). The connection between type I IFN and p38 activation in *K. pneumoniae*-infected macrophages was further corroborated by the fact that Kp52145-induced p38 phosphorylation was higher in *ifnar1^-/-^* cells than in wild-type ones (Fig S2C). Likewise, we found an increase phosphorylation of p38 in infected *tlr4^-/-^* (Fig S2D), and *tram^-/-^trif^-/-^* macrophages (Fig S2E), which is consistent with the fact that TLR4-TRAM-TRIF signalling mediates the production of *K. pneumoniae*-induced type I IFN (Ivin et al., 2017). As we anticipated, the levels of *il10* were higher in infected *tlr4^-/-^*, *tram^-/-^trif^-/-^* and *ifnar1^-/-^* macrophages compare to infected wild-type cells (Fig S2F).

Altogether, these data demonstrate that absence of SARM1 impairs *K. pneumoniae*- mediated activation of p38 due to the negative regulation exerted by type I IFN. The reduced activation of p38 limits the levels of IL10 induced by *K. pneumoniae* with a concomitant increase in inflammation.

### SARM1 negatively regulates *K. pneumoniae*-induced AIM2 inflammasome activation

The increased production of IL1β by *K. pneumoniae*-infected *sarm1^-/-^* macrophages led us to characterize the effect of SARM1 on *K. pneumoniae*-triggered inflammasome activation. Immunoblotting experiments showed elevated levels of cleavage of pro-IL1β (Fig 3A), and an increased activation of caspase 1 in infected *sarm1^-/-^* macrophages compared to infected wild-type cells (Fig 3B). Absence of SARM1 resulted in enhanced levels of processed GSDMD following infection (Fig 3C). The use of the caspase 1 inhibitor YVAD (Motani et al., 2011) confirmed that the release of IL1β by Kp52145-infected wild-type and *sarm1* macrophages was caspase 1- dependent (Fig 3D). Additional experiments supported that the adaptor protein ASC and GSDMD are required for IL1β release after inflammasome activation in *Klebsiella*-infected cells because we found a significant decrease levels of IL1β in the supernatants of infected *asc* and *gsdmd* macrophages compared to infected wild-type cells (Fig S3A). Moreover, immunoblotting experiments showed a decrease in the levels of processed pro-IL1β in the supernatants of infected *asc^-/-^* and *gsdmd^-/-^* macrophages (Fig S3B). Together, these results are consistent with enhanced inflammasome activation in *K. pneumoniae*-infected *sarm1^-/-^* macrophages. To sustain this notion further, we examined whether absence of SARM1 affects ASC speck formation. After inflammasome activation, ASC oligomerizes in large protein aggregates enabling the subsequent of clustering of caspase 1 (Cai et al., 2014; Lu et al., 2014). Therefore, detection of ASC specks is a distinguish feature of inflammasome activation. Single cell analysis by flow cytometry revealed that a greater percentage of cells displayed ASC-speck formation after Kp52145 infection of *sarm1^-/-^* macrophages (Fig 3E). Collectively, these results show that SARM1 negatively regulates inflammasome activation following *K. pneumoniae* infection.

**Figure 3.**
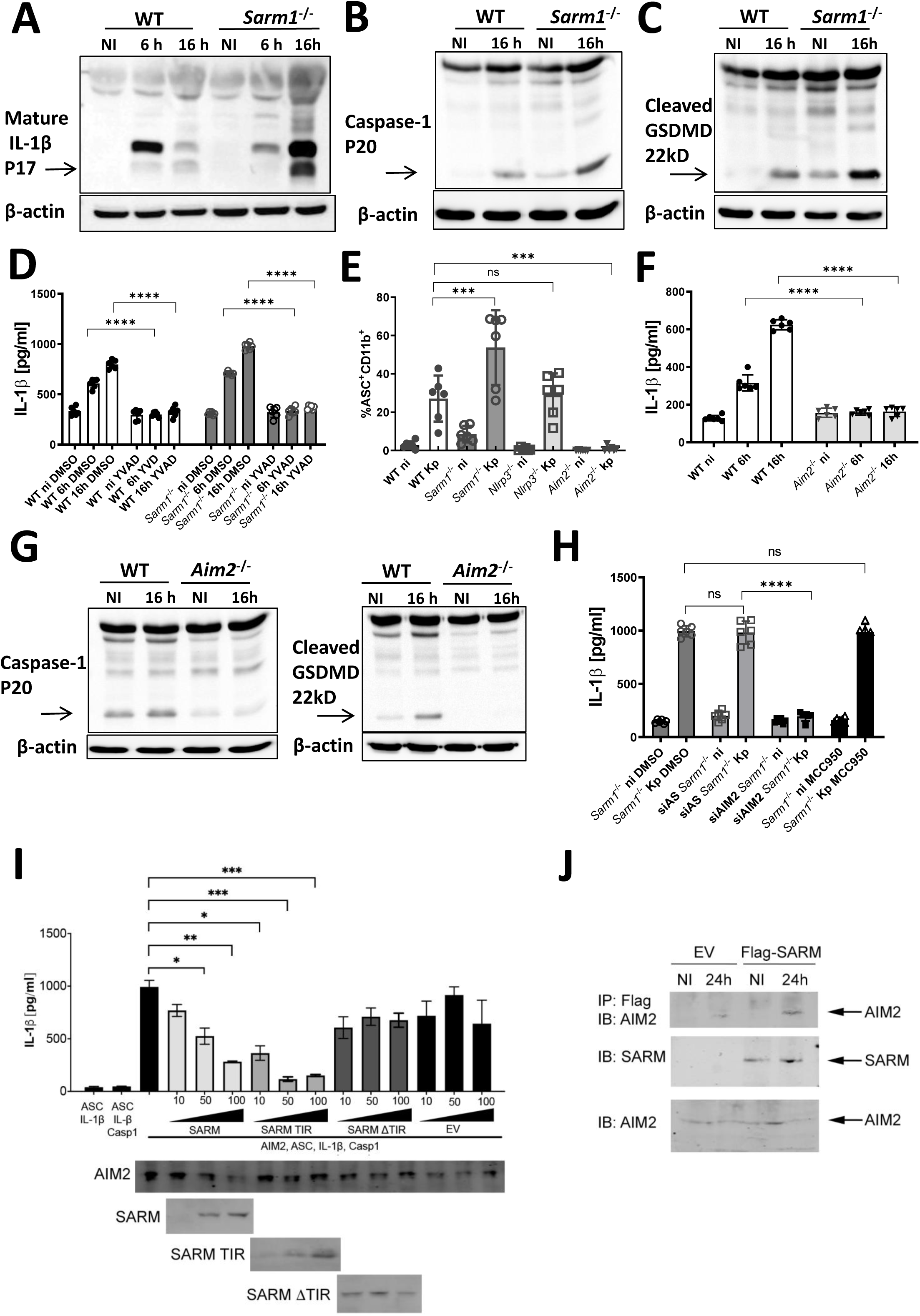
SARM1 negatively regulates *K. pneumoniae*-induced AIM2 inflammasome activation. A. Immunoblot analysis of processed pro-IL1β, and β-actin levels in lysates of wild-type macrophages (WT) and *sarm1^-/-^* macrophages non-infected (NI) or infected with Kp52145 for the indicated time points. After 1 h contact, the medium was replaced with medium containing gentamicin (100 µg/ml) to kill extracellular bacteria. B. Immunoblot analysis of processed caspase 1, and β-actin levels in lysates of wild-type macrophages (WT) and *sarm1^-/-^* macrophages non-infected (NI) or infected with Kp52145 for the indicated time points. After 1 h contact, the medium was replaced with medium containing gentamicin (100 µg/ml) to kill extracellular bacteria. C. Immunoblot analysis of cleaved gasdermin D (GSDMD), and β-actin levels in lysates of wild- type macrophages (WT) and *sarm1^-/-^* macrophages non-infected (NI) or infected with Kp52145 for the indicated time points. After 1 h contact, the medium was replaced with medium containing gentamicin (100 µg/ml) to kill extracellular bacteria. D. ELISA of IL1β secreted by wild-type (WT) and *sarm1* macrophages non-infected (ni) or infected with Kp52145 for 6 and 16h. Cells were treated with the caspse-1 inhibitor YVAD or the DMSO vehicle solution. After 1 h contact, the medium was replaced with medium containing gentamicin (100 µg/ml) to kill extracellular bacteria. E. Wild-type (WT), *sarm1^-/-^*, *nlrp3^-/-^*, and *aim2^-/-^* macrophages were non-infected (ni) or infected with Kp52145 (Kp) for 16 h, and ASC specks were detected by flow cytometry. After 1 h contact, the medium was replaced with medium containing gentamicin (100 µg/ml) to kill extracellular bacteria. F. ELISA of IL1β secreted by wild-type (WT) and *aim2* macrophages non-infected (ni) or infected with Kp52145 for 6 and16 h. After 1 h contact, the medium was replaced with medium containing gentamicin (100 µg/ml) to kill extracellular bacteria. G. Immunoblot analysis of processed caspase 1, cleaved gasdermin D (GSDMD) and β-actin levels in lysates of wild-type macrophages (WT) and *aim2^-/-^* macrophages non-infected (NI) or infected with Kp52145 for 16h. After 1 h contact, the medium was replaced with medium containing gentamicin (100 µg/ml) to kill extracellular bacteria. H. ELISA of IL1β secreted by *sarm1* macrophages treated with the NLRP3 inhibitor MCC950 or DMSO vehicle control, and *sarm1^-/-^* cells transfected with All Stars siRNA control (AS), or Aim2 siRNA (siAim2). Cells were non-infected (ni) or infected with Kp52145 (Kp) for 16 h. After 1 h contact, the medium was replaced with medium containing gentamicin (100 µg/ml) to kill extracellular bacteria. I. Reconstitution of AIM2 inflammasome activation in HEK293T cells by co-transfection of plasmids expressing HA-AIM2, ASC, procaspase-1, and pro-IL-1β. Plasmids expressing FLAG SARM1, FLAG SARM1 TIR, FLAG SARM1 ΔTIR (10, 50, 100ng), or empty vector (EV) were co-transfected. Secreted IL-1β in the culture supernatants was detected by ELISA. HA-AIM2 and FLAG SARM1 (or truncations) were detected by immunoblotting with anti-HA and anti-FLAG antibodies respectively. J. *Sarm1^-/-^* iBMDMs expressing empty vector (EV) or FLAG-SARM1 were non-infected (NI) or infected with Kp52145 for 24 h. Cells were lysed by RIPA buffer and immunoprecipitation was performed using anti-FLAG (M2) beads. The immune complexes were detected by immunoblotting with anti-SARM1, anti-AIM2 antibodies. In panels A, B, C, G, and J images are representative of three independent experiments. In panels D, E, F, H, and I values are presented as the mean ± SD of three independent experiments measured in duplicate. ****P ≤ 0.0001; ***P ≤ 0.001; **P 0.01; *P≤ 0.05; ns, P > 0.05 for the indicated comparisons determined using one way-ANOVA with Bonferroni contrast for multiple comparisons test.

We next sought to identify the inflammasome regulated by SARM1 in Kp52145-infected cells. Because SARM1 has been shown to inhibit NLRP3 (Carty et al., 2019), we asked whether NLRP3 mediates the secretion of IL1β following *K. pneumoniae* infection. However, Figure S3C shows that the NLRP3 inhibitor MCC950 (Coll et al., 2015) did not reduce Kp52145-induced secretion of IL1β. Furthermore, we found no reduction in IL1β levels in the supernatants of infected *nlrp3^-/-^* macrophages compared to wild-type cells (Fig S3D), and no decrease in the levels of cleavage pro-IL1β (Fig S3E). Control experiments showed that Kp52145 infection even increased the levels of NLRP3 (Fig S3F). Altogether, these data demonstrates that NLRP3 is not required for *K. pneumoniae* induction of IL1β. Although NLRC4 mediates IL1β secretion following infection with other Gram-negative pathogens, we consider unlikely that *K. pneumoniae* activates NLRC4 because *Klebsiella* does not express any of the bacterial proteins known to activate this inflammasome. We next considered whether AIM2, which it is also activated by Gram-negative pathogens (Ge et al., 2012; Rathinam et al., 2010; Tsuchiya et al., 2010), might mediate *K. pneumoniae-*induced release of IL1β. Indeed, IL1β release was abrogated in Kp52145-infected *aim2^-/-^* macrophages (Fig 3F). Further corroborating that AIM2 is the inflammasome activated by *K. pneumoniae*, neither caspase 1 nor GSDMD were processed in infected *aim2^-/-^* macrophages (Fig 3G). Moreover, ASC-speck formation was not detected in infected *aim2^-/-^* cells in contrast to infected wild type and *nlrp3^-/-^* cells (Fig 3E). The fact that the percentage of cells with ASC-specks was not significantly different between wild type and *nlrp3^-/-^* macrophages corroborates further that *K. pneumoniae* does not activate the NLRP3 inflammasome. Collectively, this evidence demonstrates that AIM2 is the inflammasome mediating IL1β release following *K. pneumoniae* infection. However, the possibility exists that other inflammasome(s) might be activated in the absence of SARM1. To confirm that indeed AIM2 mediates IL1β secretion in Kp52145-infected *sarm1^-/-^* macrophages, we reduced *aim2* levels by siRNA in *sarm1^-/-^* macrophages. Control experiments confirmed the knockdown efficiency (Fig S3F). As we expected, we found a reduction in IL1β levels in the supernatants of *aim2* knockdown cells compared to All stars siRNA transfected control cells (Fig 3H). Treatment of infected *sarm1^-/-^* macrophages with the NLRP3 inhibitor MCC950 did not result in any decrease in IL1β levels (Fig 3H), indicating that *K. pneumoniae* does not activate NLRP3 even in the absence of SARM1.

To examine whether SARM1 had a direct effect on AIM2, we reconstituted the AIM2 inflammasome in HEK293 cells by transfecting plasmids expressing pro-IL-1β, pro-caspase-1, ASC, and AIM2. Under these conditions, the inflammasome is active to induce the secretion of IL1β without external stimulus (Shi et al., 2016), and this is AIM2-dependent since no detectable mature IL-1β was produced from cells transfected with all the inflammasome components except AIM2 (Fig 3I). AIM2-dependent secretion of IL1β was inhibited by the expression of SARM1 (Fig 3I). We next determined which domains of SARM1 were required for AIM2 inhibition by expressing different truncations of SARM1. This experiment showed that the TIR domain alone was sufficient to inhibit IL1β release (Fig 3I). These data led us to determine whether *K. pneumoniae* induces the interaction between SARM1 and AIM2 to inhibit inflammasome activation. We carried out co-immunoprecipitation experiments infecting retrovirally transfected FLAG-SARM1 in *sarm1^-/-^* macrophages. Figure 3J shows that SARM1 immunoprecipitated AIM2 only in Kp52145-infected cells.

Altogether, we propose that *K. pneumoniae* exploits SARM1 to inhibit AIM2 inflammasome activation by a direct interaction between SARM1 and AIM2.

### *K. pneumoniae* induces AIM2 in a type I IFN-dependent manner

We next sought to investigate whether *K. pneumoniae* infection affects the expression levels of AIM2. RT-qPCR analysis revealed that Kp52145 induced the expression of *aim2* in vitro (Fig 4A), and in the lungs of infected mice (Fig 4B). Western blot experiments demonstrated that Kp52145 increased the expression of AIM2 in wild-type macrophages (Fig 4C). We next investigated the signalling pathways governing *K. pneumoniae* induction of *aim2*. *Aim2* has been identified as an ISG (Fernandes-Alnemri et al., 2009), and the interferome prediction tool (Rusinova et al., 2013) indicates that type I IFN activates the expression of *aim2* in human and mouse cells. Consistent with this prediction, *aim2* and AIM2 levels were reduced in *Klebsiella* infected *ifnar1^-/-^* cells (Fig 4D). We then tested whether *K. pneumoniae* would induce *aim2* and AIM2 in cells deficient for the TLR4-TRAM-TRIF-IRF3 pathway mediating type I IFN production by *K. pneumoniae* (Ivin et al., 2017). Indeed, Kp52145 did not increase *aim2* levels in *tlr4^-/-^*, *tram^-/-^trif^-/-^* and *irf3^-/-^* macrophages (Fig S4A). No significant differences were found between infected wild-type and *myd88^-/-^* macrophages (Fig S4A). Kp52145 did not increase AIM2 levels in *tlr4^-/-^*, and *tram^-/-^trif^-/-^* macrophages (Fig S4B). Together, these results demonstrate that *K. pneumoniae* infection induces AIM2 in a type I IFN-dependent manner following activation of TLR4-TRAM-TRIF-IRF3 pathway. These results led us to investigate whether the capsule polysaccharide (CPS), and LPS O-polysaccharide, mediating the production of type I IFN following *K. pneumoniae* infection (Ivin et al., 2017), are involved in *aim2* induction. Cells were infected with single mutants lacking each of the polysaccharides, and a double mutant lacking both (Ivin et al., 2017; Sa-Pessoa et al., 2020). Indeed, the three mutants induced less *aim2* than the wild-type strain (Fig 4E). Furthermore, immunoblotting analysis showed that the three mutants did not increase AIM2 levels (Fig 4F). As anticipated, the CPS and LPS O-polysaccharide mutants induced less IL1β than the wild-type strain, being the double mutant the strain inducing the lowest IL1β levels (Fig 4G).

**Figure 4.**
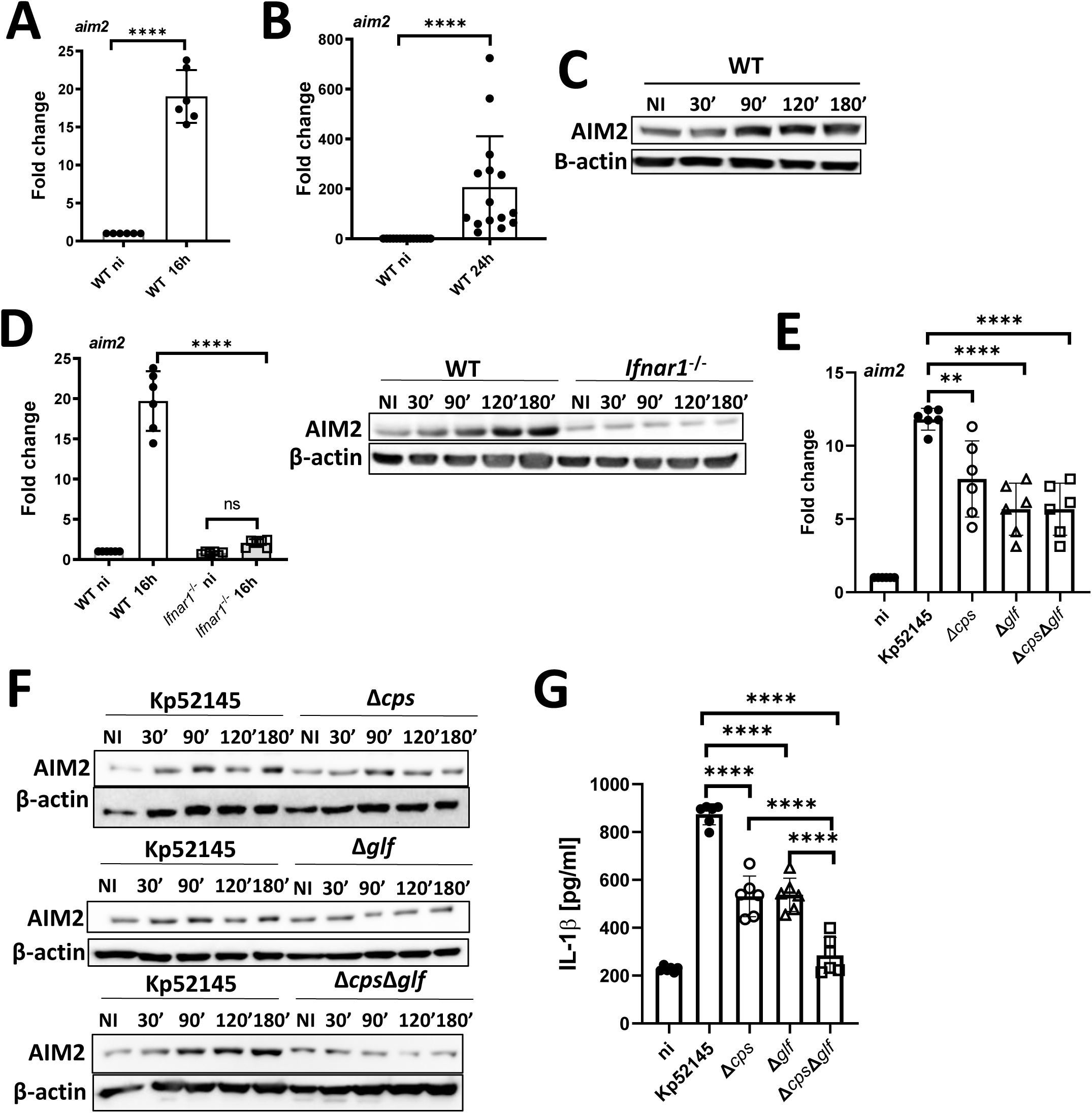
*K. pneumoniae* induces AIM2 in a type I IFN-dependent manner. A. *aim2* mRNA levels were assessed by qPCR in wild-type macrophages (WT) non-infected (ni) or infected with Kp52145 for 16 h. After 1 h contact, the medium was replaced with medium containing gentamicin (100 µg/ml) to kill extracellular bacteria. B. *aim2* mRNA levels were assessed by qPCR in the lungs of infected wild-type mice (WT) for 24 h. C. Immunoblot analysis of AIM2 and β-actin levels in lysates of wild-type macrophages (WT) macrophages non-infected (NI) or infected with Kp52145 for the indicated time points. After 1 h contact, the medium was replaced with medium containing gentamicin (100 µg/ml) to kill extracellular bacteria. D. *aim2* mRNA levels were assessed by qPCR in wild-type (WT), and *ifnar1^-/-^* macrophages non-infected (ni) or infected with Kp52145 for 16 h. Immunoblot analysis of Aim2 and β-actin levels in lysates of wild-type macrophages (WT) and *ifnar1^-/-^* macrophages non-infected (NI) or infected with Kp52145 for the indicated time points. After 1 h contact, the medium was replaced with medium containing gentamicin (100 µg/ml) to kill extracellular bacteria. E. *aim2* mRNA levels were assessed by qPCR in wild-type macrophages (WT) non-infected (ni) or infected with Kp52145, the capsule mutant 52145-Δ*manC* (Δ*cps*), the mutant lacking the LPS O-polysaccharide, 52145-Δ*glf* (Δ*glf),* and the double mutant lacking the CPS and the LPS O-polysaccharide, 52145- Δ*wca* -Δ*glf (*Δ*cps*Δ*glf)* for 16 h. After 1 h contact, the medium was replaced with medium containing gentamicin (100 µg/ml) to kill extracellular bacteria. F. Immunoblot analysis of Aim2 and β-actin levels in lysates of wild-type macrophages (WT) macrophages non-infected (NI) or infected with Kp52145, the capsule mutant 52145-Δ*manC (*Δ*cps*) the mutant lacking the LPS O-polysaccharide, 52145-Δ*glf* (Δ*glf),* and the double mutant lacking the CPS and the LPS O-polysaccharide, 52145- *wca_k2_*- Δ*glf (*Δ*cps* Δ*glf)* for the indicated time points. After 1 h contact, the medium was replaced with medium containing gentamicin (100µg/ml) to kill extracellular bacteria. G. ELISA of IL1β secreted by wild-type macrophages non-infected (ni) or infected with Kp52145, the capsule mutant 52145-Δ*manC* (Δ*cps*), the mutant lacking the LPS O-polysaccharide, 52145-Δ*glf* (Δ*glf),* and the double mutant lacking the CPS and the LPS O-polysaccharide, 52145-Δ*wcak2*-Δ*glf (*Δ*cps* Δ*glf)* for 16 h. After 1 h contact, the medium was replaced with medium containing gentamicin (100 μg/ml) to kill extracellular bacteria In panels C, D, and F images are representative of three independent experiments. In panels A, B, D, E, and G values are presented as the mean ± SD of three independent experiments measured in duplicate. ****P ≤ 0.0001; ** P ≤ 0.01; ns, P > 0.05 for the indicated comparisons determined using one way-ANOVA with Bonferroni contrast for multiple comparisons test.

To further link type I IFN signalling to *K. pneumoniae*-induced AIM2 inflammasome activation, we determined IL1β production in cells deficient for the signalling pathway mediating type I IFN production following *K. pneumoniae* infection. As expected, Kp52145 did not induce the release of IL1β in *tlr4* , *tram trif* and *ifnar1* macrophages (Fig S4C). Control experiments showed that pro-ILβ production was not significantly reduced in infected *tlr4^-/-^* cells, ruling out that the lack of IL1β production in the absence of TLR4 was due to reduced levels of pro-IL1β (Fig S4D).

Collectively, these results demonstrate that signalling via IFNAR1 is required for activation of AIM2 inflammasome by *K. pneumoniae* upon recognition of the CPS and the LPS O-polysaccharide by TLR4.

### *K. pneumoniae* induces SARM1 in a type I IFN-dependent manner

It is common for pathogens to upregulate or activate the expression of the host proteins they do target for their own benefit. It might be then expected that *K. pneumoniae* upregulates the expression of SARM1. Indeed, Kp52145 induced the expression of *sarm1* in vitro (Fig 5A), and in the lungs of infected mice (Fig 5B). Infection of *Sarm1^FLAG^* cells confirmed that Kp52145 increased the expression of SARM1 (Fig 5C). We next sought to identify the signalling pathways governing *K. pneumoniae* induction of *sarm1*; however, the regulation of SARM1 is poorly understood.

**Figure 5.**
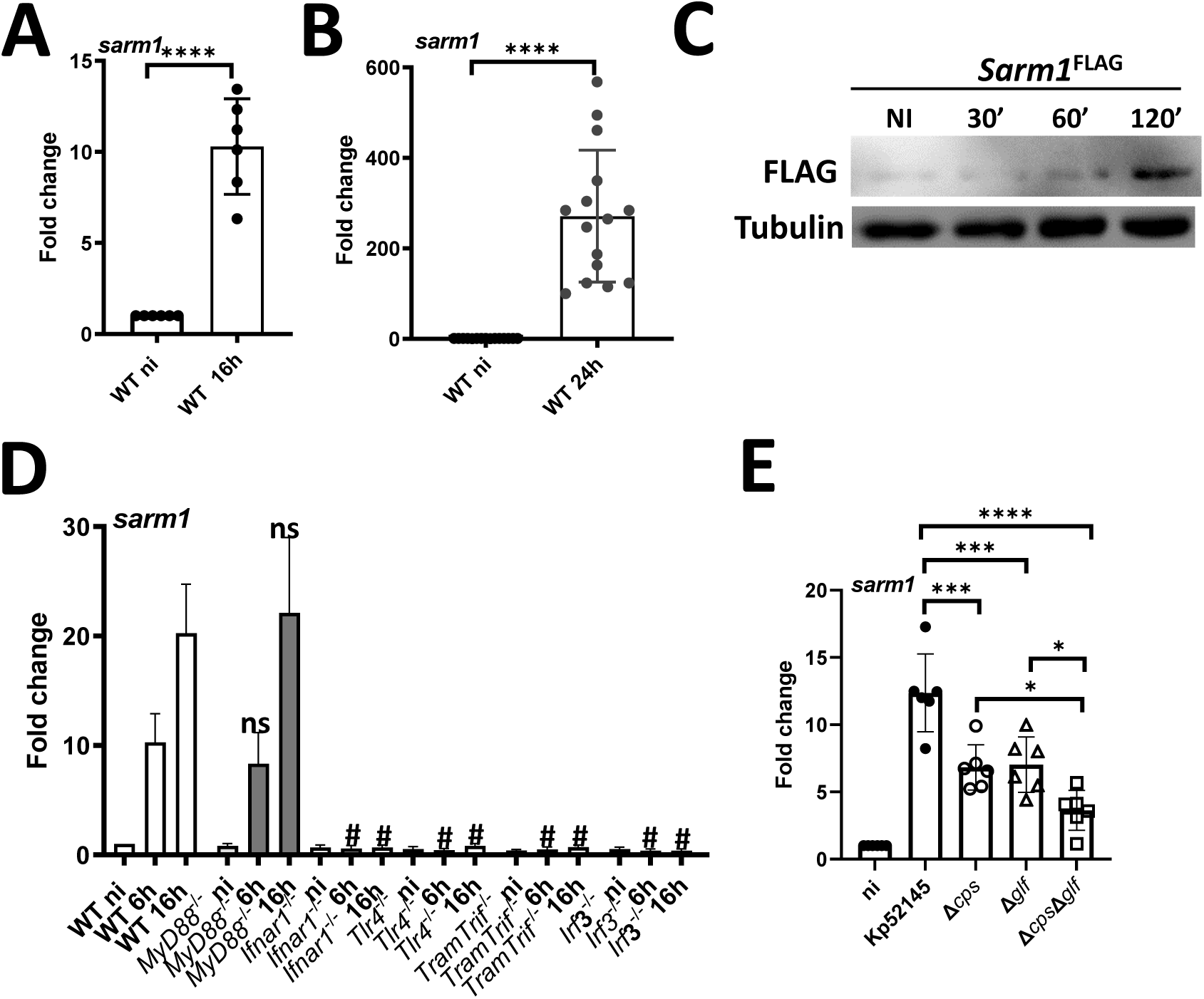
*K. pneumoniae* induces SARM1 in a type I IFN-dependent manner. A. *sarm1* mRNA levels were assessed by qPCR in wild-type macrophages (WT) non-infected (ni) or infected with Kp52145 for 16 h. After 1 h contact, the medium was replaced with medium containing gentamicin (100 µg/ml) to kill extracellular bacteria. B. *sarm1* mRNA levels were assessed by qPCR in the lungs of infected wild-type mice (WT) for 24 h. C. Immunoblot analysis of SARM1-FLAG and tubulin levels in lysates of *Sarm1^FLAG^* macrophages non-infected (NI) or infected with Kp52145 for the indicated time points. After 1 h contact, the medium was replaced with medium containing gentamicin (100 µg/ml) to kill extracellular bacteria. D. *sarm1* mRNA levels were assessed by qPCR in wild-type macrophages (WT), *myd88^-/-^*, *ifnar1^-/-^*, *tlr4^-/-^*, *tram^-/-^trif^-/-^*, and *irf3^-/-^* non-infected (ni) or infected with Kp52145 for 6 and 16 h. After 1 h contact, the medium was replaced with medium containing gentamicin (100 µg/ml) to kill extracellular bacteria. E. *sarm1* mRNA levels were assessed by qPCR in wild-type macrophages (WT) non-infected (ni) or infected with Kp52145, the capsule mutant 52145-Δ*manC* (Δ*cps*), the mutant lacking the LPS O polysaccharide, 52145-Δ*glf* (Δ*glf),* and the double mutant lacking the CPS and the LPS O polysaccharide, 52145-Δ*wca_k2_*- Δ*glf (*Δ*cps* Δ*glf)* for 16 h. After 1 h contact, the medium was replaced with medium containing gentamicin (100 μg/ml) to kill extracellular bacteria. In panel C, image is representative of three independent experiments. In panels A, B, D, and E values are presented as the mean ± SD of three independent experiments measured in duplicate. In panels A, B and D ****P ≤ 0.0001; ***P ≤ 0.001; *P ≤ 0.05 for the indicated comparisons; in panel C # P ≤ 0.0001; ns, P > 0.05 for the comparisons between the knock-out and wild-type cells at the same time point post infection. Significance was established using one way-ANOVA with Bonferroni contrast for multiple comparisons test.

Analysis of the promoter region of SARM1 interrogating the interferome database (Rusinova et al., 2013) identified SARM1 as an ISG. Therefore, we speculated that *K. pneumoniae* may regulate SARM1 in a type I IFN-dependent manner. Providing initial support to this notion, RT-qPCR analysis showed that Kp52145 did not induce *sarm1* in *ifnar1^-/-^* macrophages (Fig 5C). Furthermore, *sarm1* levels were reduced in infected *tlr4^-/-^*, *tram^-/-^trif^-/-^*, and *irf3^-/-^* macrophages (Fig 5C). As anticipated, Kp52145 induced *sarm1* in *myd88^-/-^* macrophages (Fig 5C). These results led us to investigate whether the CPS, and the LPS O-polysaccharide are involved in *sarm1* induction. Indeed, the three mutants induced less *sarm1* than the wild-type strain, although the double mutant lacking CPS and the LPS O-polysaccharide induced less *sarm1* than each of the single mutants did (Fig 5D).

Altogether, these results confirm experimentally that *K. pneumoniae* leverages type I IFN signalling to induce SARM1 following activation of TLR4-TRAM-TRIF-IRF3 pathway. The CPS and the LPS O-polysaccharide are the *K. pneumoniae* factors responsible for the upregulation of the expression of SARM1.

### SARM1 promotes *K. pneumoniae* virulence

Having established that *K. pneumoniae* exploits the immunomodulatory roles of SARM1 to control MyD88 and TRIF-governed cytokine production, and the activation of AIM2 inflammasome, we next sought to investigate whether SARM1 contributes to *K. pneumoniae* subversion of cell-autonomous immunity. We have demonstrated that *K. pneumoniae* manipulates the phagosome traffic following phagocytosis to create a unique niche that does not fuse with lysosomes, the KCV, allowing the intracellular survival of *Klebsiella* (Cano et al., 2015). Therefore, we asked whether the absence of SARM1 impairs *K. pneumoniae* intracellular survival. Control experiments revealed that the attachment of Kp52145 was not affected in *sarm1^-/-^* cells (Fig S5A) whereas there was a slight reduction in the number of engulfed bacteria (Fig S5B). Time-course experiments revealed that the intracellular survival of Kp52145 was significantly reduced in *sarm1^-/-^* macrophages (Fig 6A). We then sought to determine whether the reduced intracellular survival was due to an increase in the colocalization of lysosomes with the KCV. Lysosomes were labelled with the membrane-permeant fluorophore cresyl violet (Ostrowski et al., 2016), and cells were infected with GFP-labelled Kp52145 to assess the KCV at the single cell level by immunofluorescence. Confocal microscopy experiments revealed that the majority of the KCVs from wild-type macrophages did not colocalize with cresyl violet (Fig 6B and Fig 6C), corroborating our previous work (Cano et al., 2015). In contrast, there was an increase in the colocalization of KCVs from *sarm1^-/-^* macrophages with cresyl violet (Fig 6B and Fig 6C), demonstrating that the absence of SARM1 results in the fusion of the KCV with lysosomes with a concomitant reduction in the numbers of intracellular bacteria.

**Figure 6.**
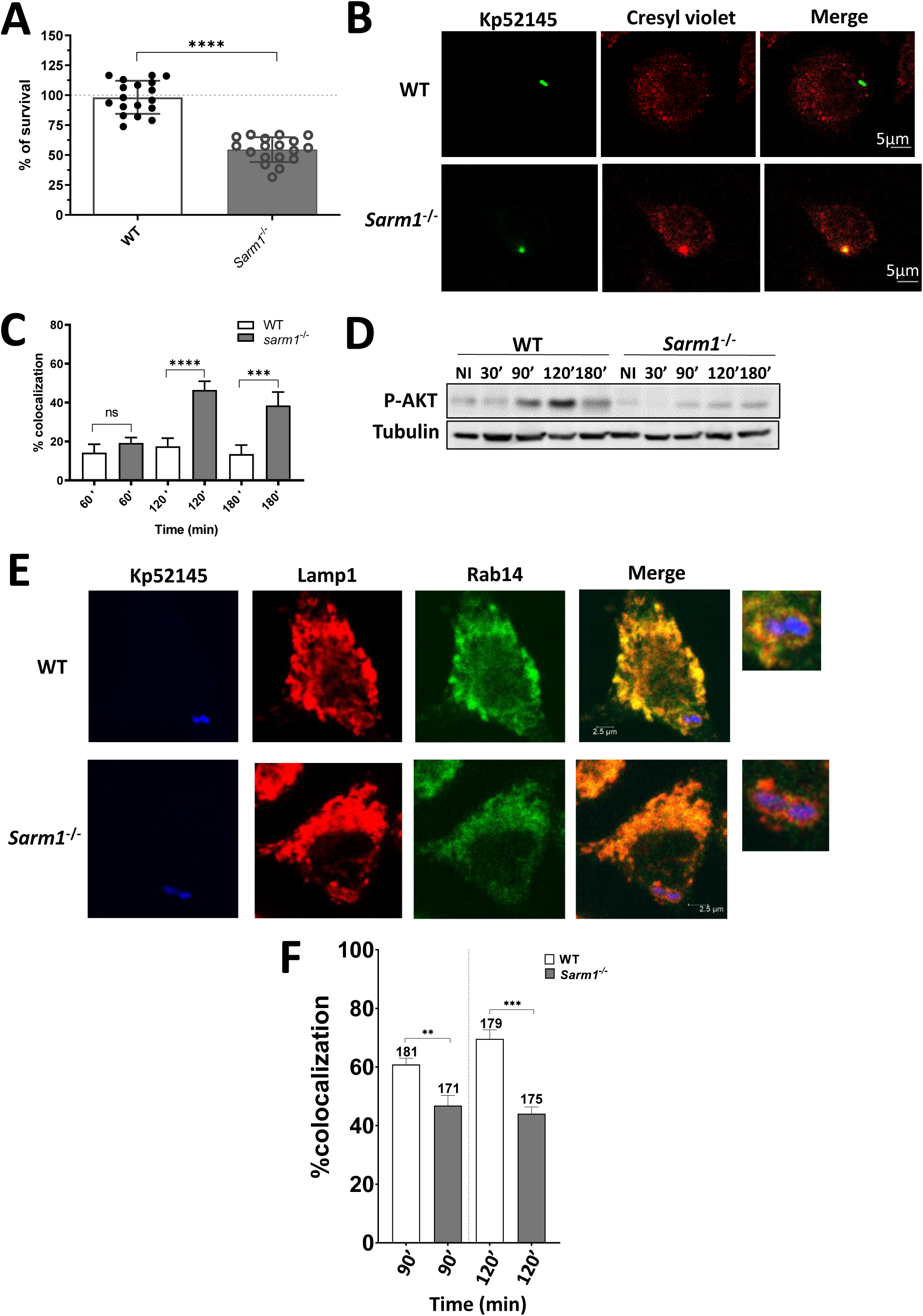
SARM1 is required for *K. pneumoniae* intracellular survival. A. Kp52145 intracellular survival in wild-type (WT) and *sarm1^-/-^* 4 h after addition of gentamycin (30 min of contact). Results are expressed as % of survival (CFUs at 4 h versus 1 h in *sarm1*^-/-^ cells normalized to the results obtained in wild-type macrophages set to 100%). Values are presented as the mean ± SD of six independent experiments measured in triplicate. B. Immunofluorescence confocal microscopy of the colocalziation of Kp52145 harbouring pFPV25.1Cm and cresyl violet in wild-type (WT) and *sarm1^-/-^* macrophages. The images were taken 90 min post infection. Images are representative of duplicate coverslips in three independent experiments. C. Percentage of Kp52145 harbouring pFPV25.1Cm co-localization with cresyl violet over a time course. Wild-type (WT) and *sarm1^-/-^* macrophages were infected; coverslips were fixed and stained at the indicated times. Values are given as mean percentage of Kp52145 co-localizing with the marker ± SD At least 200 infected cells belonging to three independent experiments were counted per time point. D. Immunoblot analysis of phosphorylated Akt (P-AKT), and tubulin levels in lysates of wild-type (WT) and *sarm1^-/-^* macrophages non-infected (NI) or infected with Kp52145 for the indicated time points. After 1 h contact, the medium was replaced with medium containing gentamicin (100 µg/ml) to kill extracellular bacteria. Images are representative of three independent experiments. E. Immunofluorescence confocal microscopy of the colocalization of Kp52145 harbouring pFPV25.1Cm, Lamp1, and Rab14 in wild-type (WT) and *sarm1^-/-^* macrophages. The images were taken 90 min post infection. Images are representative of duplicate coverslips in three independent experiments. F. Percentage of Kp52145 harbouring pFPV25.1Cm co-localization with Lamp1 and Rab14 over a time course. Wild-type (WT) and *sarm1^-/-^* macrophages were infected; coverslips were fixed and stained at the indicated times. Values are given as mean percentage of Kp52145 co-localizing with the marker ± SD. The number of infected cells counted per time in three independent experiments are indicated in the figure. In panels, A, C and F, values are presented as the mean ± SD of three independent experiments measured in duplicate. ****P ≤ 0.0001; ***P ≤ 0.001; ** P ≤ 0.01; ns, P > 0.05 for the indicated comparisons determined using unpaired t test.

Previously, we showed that *K. pneumoniae* targets the PI3K-AKT axis to survive intracellularly (Cano et al., 2015). Therefore, we asked whether the absence of SARM1 would affect *K. pneumoniae*-induced AKT phosphorylation. Immunoblotting experiments confirmed that Kp52145-induced phosphorylation of AKT was reduced in *sarm1^-/-^* macrophages compare to wild-type cells (Fig 6D). In *K. pneumoniae*-infected cells, AKT activation is linked to the recruitment of Rab14 to the KCV to block the fusion with lysosomes (Cano et al., 2015). We then investigated the recruitment of Rab14 to the KCV in *sarm1^-/-^* macrophages. Figure 6E illustrates that Rab14 does not colocalize with the KCV in *sarm1^-/-^* macrophages in contrast to wild-type macrophages. Altogether, this evidence demonstrates that SARM1 is crucial for *K. pneumoniae*-induced activation of the PI3K–AKT–Rab14 axis to control the phagosome maturation to survive inside macrophages.

To obtain a global view of the role of SARM1 in *K. pneumoniae* infection biology, we examined the contribution of SARM1 to modulate the inflammatory responses induced by *K. pneumoniae* in vivo. We analysed several inflammatory-associated cytokines and chemokines in the lungs of *K. pneumoniae*-infected animals. Kp52145 induced the expression of *il1b*, *tnfa*, *il12*, *cxcl10 ifnb,* and *isg15* in vivo (Fig 7A), although the levels of *il1b*, *tnfa*, and *il12* were significantly higher in *sarm1^-/-^* mice than in wild-type ones. Furthermore, we observed a significant decrease in the levels of *il10* in *sarm1^-/-^* mice compare to wild-type ones (Fig 7B). Similar results were obtained infecting *Sarm1^em1.1Tft^* mice, indicating that the results are neither dependent on the mouse strain nor on the way the *sarm1* knock-out mice were generated (Fig 7A and Fig 7B). Together, these results demonstrate that the absence of SARM1 in vivo results in heightened inflammation following *K. pneumoniae* infection.

**Figure 7.**
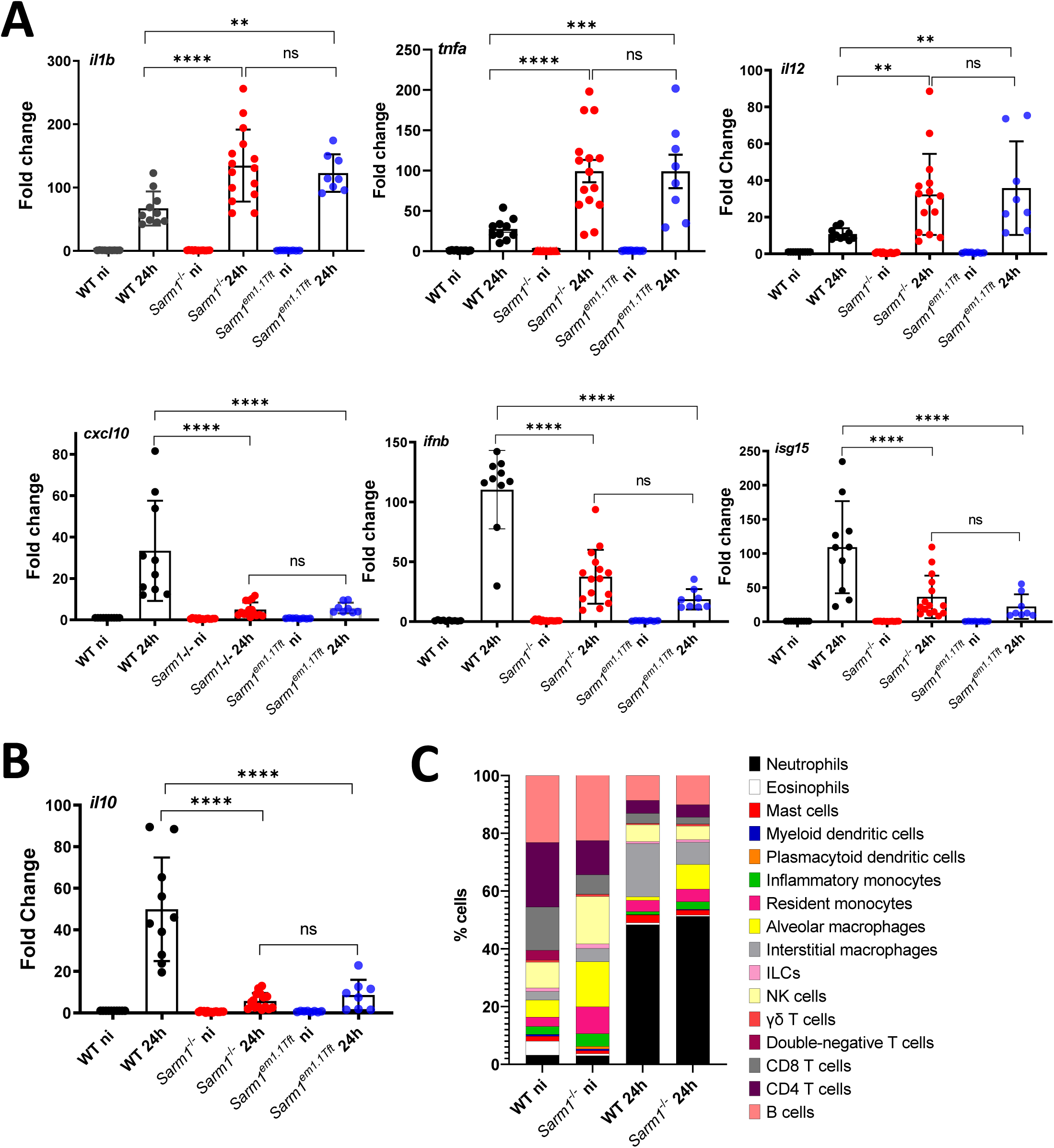
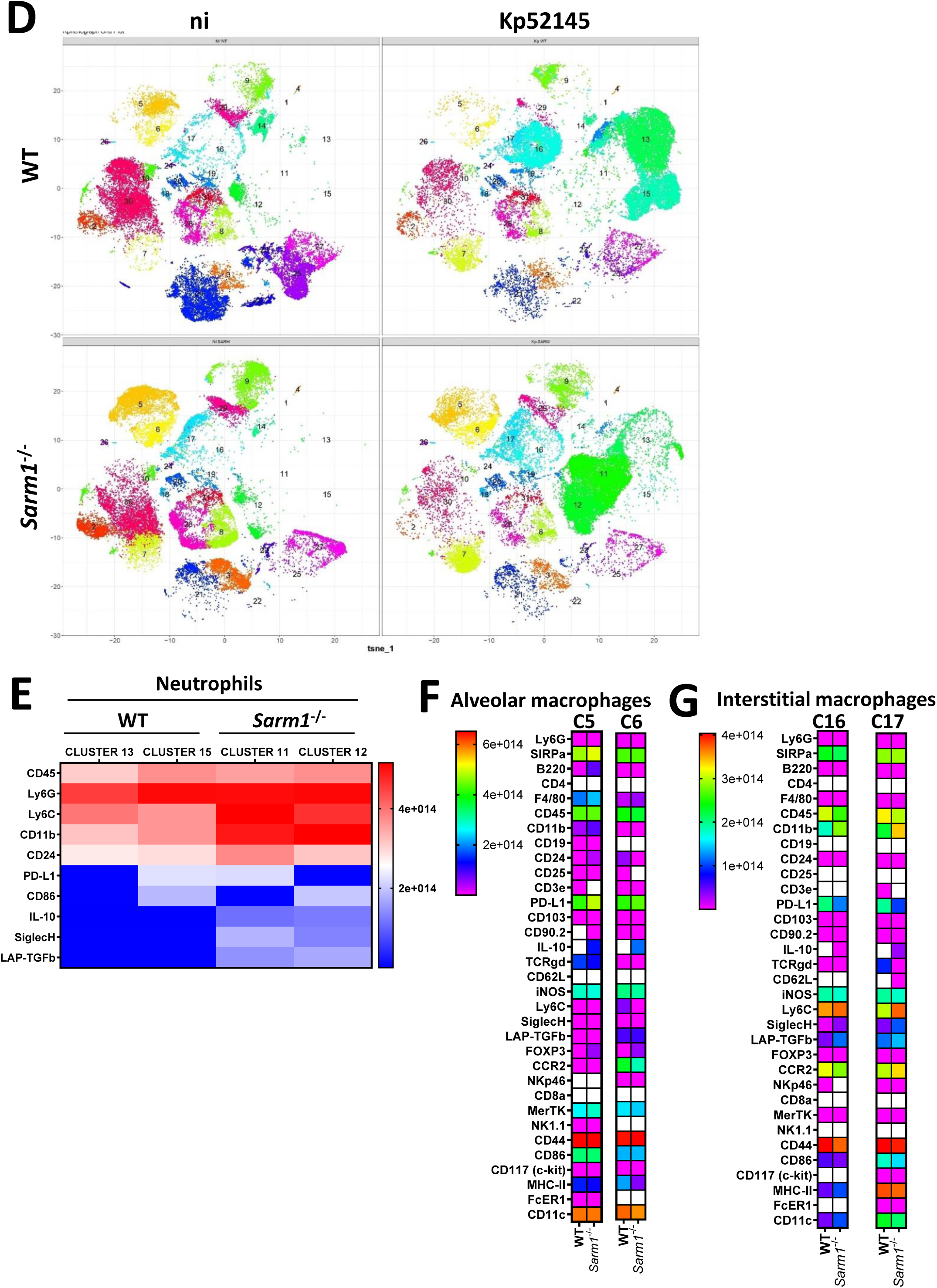
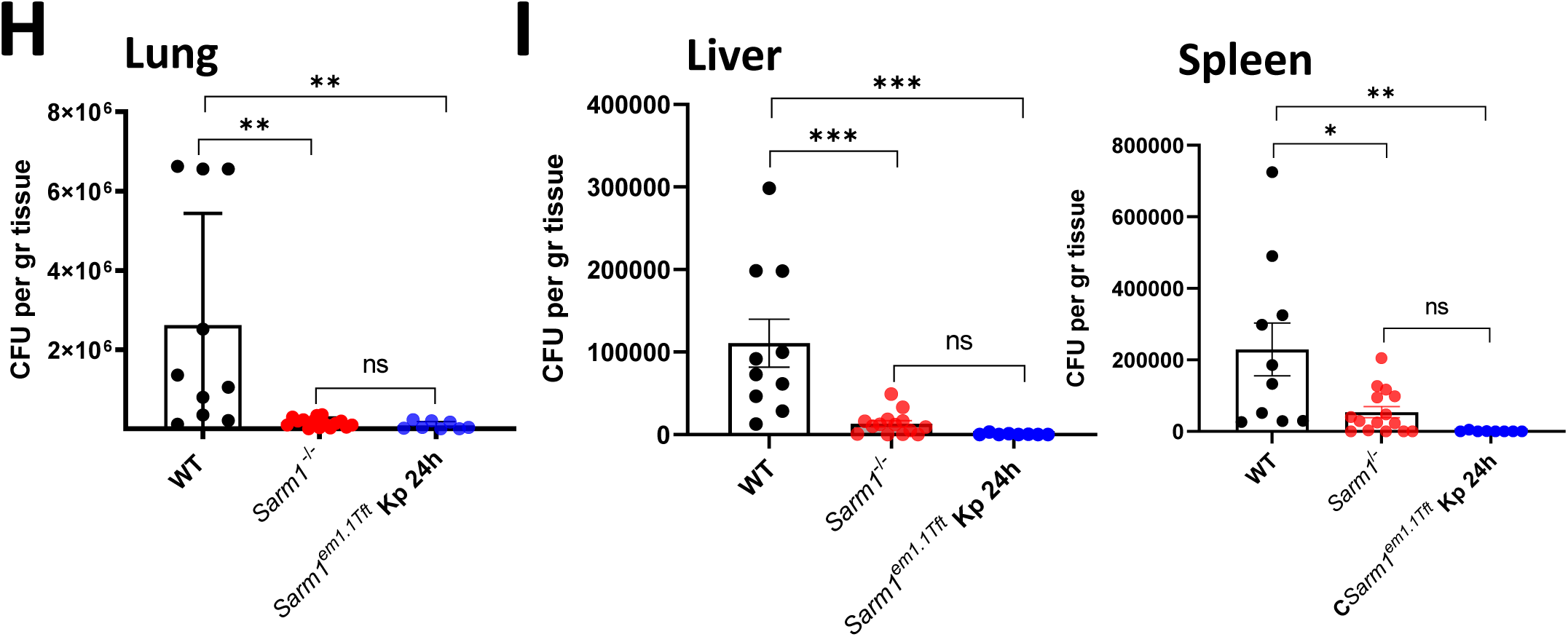
SARM1 promotes *K. pneumoniae* virulence. A. *il1b, tnfa, il12, cxcl10, ifnb, and isg15* mRNA levels were assessed by qPCR in the lungs of infected wild-type mice (WT), *sarm1^-/-^*, and *Sarm1^em1.1Tft^* for 24. Each dot represents a different mouse. B. *il10* mRNA levels were assessed by qPCR in the lungs of infected wild-type (WT), *sarm1^-/-^*, and *Sarm1^em1.1Tft^* mice for 24. C. Percentage of immune cells in the lungs of wild-type (WT), and *sarm1^-/-^* mice non-infected (ni) or infected intranasally with Kp52145 for 24. Results are based on data from three mice per group. D. PhenoGraph cluster analysis of immune populations in the lungs wild-type (WT), and *sarm1^-/-^* mice non-infected (ni) or infected intranasally with Kp52145 for 24. Results are based on data from three mice per group. E. Heat map showing relative signal intensities of the indicated markers on neutrophils of clusters 13, 15 found in the lungs of infected wild-type mice, and clusters 11 and 13 detected in the lungs of *sarm1^-/-^* mice. The heat map is coloured based on signal intensity of the indicated markers. Results are based on data from three mice per group. F. Heat map showing relative signal intensities of the indicated markers on alveolar macrophages of clusters 5 and 6 found in the lungs of infected wild-type and *sarm1^-/-^* mice. The heat map is coloured based on signal intensity of the indicated markers. Results are based on data from three mice per group. G. Heat map showing relative signal intensities of the indicated markers on interstitial macrophages of clusters 16 and 17 found in the lungs of infected wild-type and *sarm1^-/-^* mice. The heat map is coloured based on signal intensity of the indicated markers. Results are based on data from three mice per group. H. Bacterial load in the lungs of infected wild-type mice (WT), *sarm1^-/-^*, and *Sarm1^em1.1Tft^* for 24. Each dot represents a different mouse. I. Bacterial load in the livers and spleens of infected wild-type mice (WT), *sarm1^-/-^*, and *Sarm1^em1.1Tft^* for 24. Each dot represents a different mouse. In panels A, B, H and I values are presented as the mean ± SD of three independent experiments measured in duplicate. ****P ≤ 0.0001; ***P ≤ 0.001; **P≤ 0.01; *P ≤ 0.05; ns, P > 0.05 for the indicated comparisons using one way-ANOVA with Bonferroni contrast for multiple comparisons test.

To find out whether the absence of SARM1 has any effect on immune cells, we used mass cytometry to profile the cells of infected and non-infected mice. We tested a panel of 33 surface and intracellular markers that would enable resolution of 100 lymphoid and myeloid cell types (Table S1). To define cell communities, we employed the clustering algorithm PhenoGraph (Levine et al., 2015). In *sarm1^-/-^* non-infected mice, we found a significant increase in the numbers of resident monocytes (MHC-II-Ly6G-Ly6C^+^CD11b^+^CD11c-CCR2^high^) (p < 0.05) compare to non-infected wild-type mice, whereas there were no significant differences in the numbers of any other immune cell (Fig 7C). Following infection, we observed a significant increase in the numbers of neutrophils (MHC-II^-^Ly6G^+^Ly6C^+^F4/80^-/low^) in both genotypes, although the numbers were not significantly different between them (Fig 7C). In infected wild-type mice there was an increase in the numbers of interstitial macrophages (MHC-II^+^Ly6G^-^Ly6C^+^CD11b^+^) compared to infected *sarm1^-/-^* mice (p< 0.01) (Fig 7C). In contrast, there was a significant increase in the number of alveolar macrophages (MHC-II^+^Ly6G^-^Ly6C^-^CD11b^low^CD11c^+^) (p < 0.01) in infected *sarm1^-/-^* mice compared to infected wild-type ones (Fig 7C). The numbers of alveolar macrophages were not significantly different between infected and non-infected *sarm1^-/-^* mice. There were no significant differences in the numbers of other immune cells between infected genotypes (Fig 7C).

PhenoGraph analysis identified 30 clusters with similar marker expression (Fig S6A and Table S2). The heat map of the markers expressed by each of the clusters is shown in Figure S6B. Differences were found between samples (Fig 7D, and Fig S6C). Clusters 11 and 12 were only present in *sarm1^-/-^* infected mice whereas clusters 13 and 15 were only present in wild-type-infected mice. These four clusters represent different subsets of neutrophils (Fig S6A). Heat map analysis of these clusters revealed that in both genetic backgrounds each of the clusters can be differentiated based on the expression levels of PD-L1 and CD86 (Fig 7E). Clusters 11 and 12 were characterized by the expression levels of the markers Ly6C, CD11b, CD24, IL10, Siglec H and LAP-TGFβ (Fig 7B), revealing an increase activation of neutrophils in *sarm1^-/-^* mice following infection. Clusters 5 and 6, corresponding to alveolar macrophages (Fig S6A), were predominant in *sarm1*^-/-^ mice, and they can be differentiated by the expression of CCR2. The expression of CCR2 is higher in cluster 6 than in cluster 5. No major differences were noted between genetic backgrounds with infection, except that *sarm1^-/-^* cells showed an increase in the levels of IL10 (p <0.0001) (Fig 7H). Differences were found between the subsets of interstitial macrophages. Cluster 16 was the predominant in wild-type infected mice, whereas cluster 17 was the predominant one in *sarm1^-/-^* infected mice (Fig 7D). The levels of CD11c differentiates both clusters, higher in cluster 17 than in cluster 16 (Fig 7G). High levels of CD11c are associated with the activation of immune cells (Arnold et al., 2016; Lewis et al., 2015). Cells belonging to cluster 17, found in *sarm1*^-/-^ mice, were characterized by high levels of CD11b, iNOS, Ly6C, CCR2, Siglec H, SIRPα, LAP-TGFβ, CD44, CD86. MHC-II, and Cd11c (Fig 7G), all markers of activation of immune cells. Altogether, mass cytometry analysis demonstrate an increase in the numbers of neutrophils and alveolar macrophages in *sarm1^-/-^* infected mice. These cells are crucial in host defence against *K. pneumoniae* (Broug-Holub et al., 1997; Xiong et al., 2015; Xiong et al., 2016; Ye et al., 2001). Furthermore, PhenoGraph cluster analysis revealed the presence of different subsets of neutrophils and interstitial macrophages in *sarm1^-/-^* infected mice characterized by elevated levels of markers associated with the activation of immune cells.

Finally, we determined the ability of *sarm1^-/-^* mice to control bacterial growth following intranasal infection. At 24 h post infection, there was a 94% reduction in bacterial load in the lungs of infected *sarm1^-/-^* mice compared to wild-type infected ones (Fig 7H). Moreover, we found a significant lower dissemination of Kp52145 to liver and spleen in *sarm1^-/-^* mice than in wild-type ones (Fig 7I). The infection of *Sarm1^em1.1Tft^* mice yielded similar results; the knockout mice controlled the lung infection more efficiently than the wild-type ones and there was less dissemination to deeper tissues (Fig 7H and Fig 7I). Altogether, this evidence establishes the crucial role of SARM1 for *K. pneumoniae* survival in vivo.

## DISCUSSION

The human pathogen *K. pneumoniae* exemplifies the global threat of antibiotic resistant bacteria. Hundreds of mobile antimicrobial resistant genes are found in *K. pneumoniae*, and these can be disseminated to other bacteria. *K. pneumoniae* is the species within which several new antimicrobial resistance genes were first discovered (e.g. carbapenem-resistance genes KPC, OXA- 48 and NDM-1). Less obvious, but central to pathogenesis, are *K. pneumoniae* adaptations to the human immune system allowing the pathogen to flourish in the tissues. However, our knowledge of the strategies deployed by *K. pneumoniae* to counteract the innate immune system is still elementary, as it is our understanding of which of such responses benefit the host versus *K. pneumoniae*. In this study, we show that *K. pneumoniae* exploits SARM1 to control MyD88 and TRIF-governed inflammation, to limit the activation of the MAP kinases ERK and JNK, and to induce the anti-inflammatory cytokine IL10 by fine-tuning the p38-type I IFN axis. SARM1 also inhibits the activation of *K. pneumoniae*-induced AIM2 inflammasome with a concomitant reduction in IL1β (Fig 8) to further supress inflammatory responses. We have established that SARM1 is necessary for *K. pneumoniae* intracellular survival whereas, in vivo, absence of SARM1 facilitates the clearance of the pathogen. Altogether, these results demonstrate that SARM1 plays an integral role in *K. pneumoniae* infection biology. Manipulation of the Toll-IL-1R protein SARM1 is a hitherto unknown anti-immunology strategy deployed by a human pathogen.

**Figure 8.**
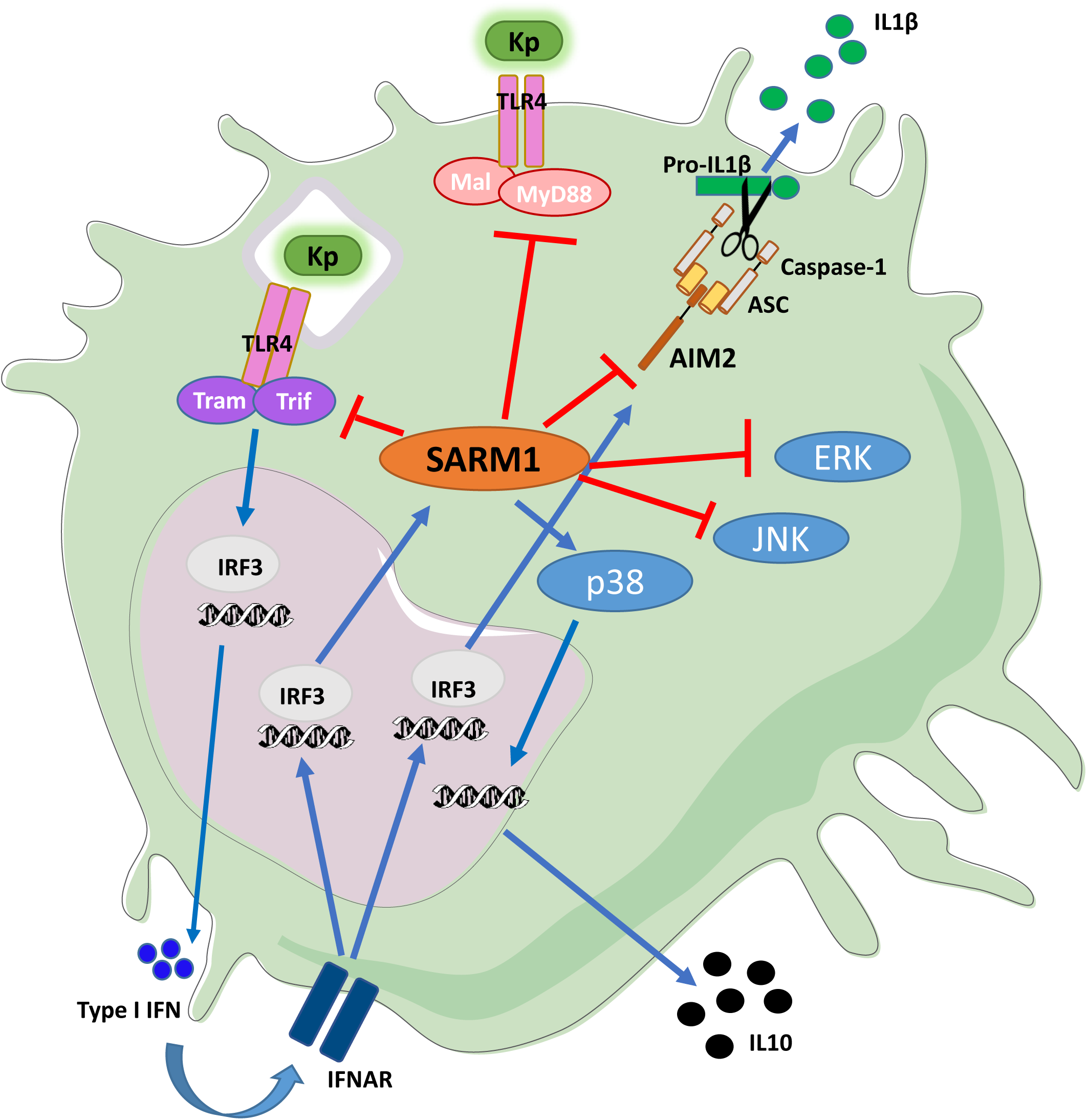
*K. pneumoniae* exploits the immunomodulatory properties of SARM1 to antagonize cell intrinsic immunity. Kp52145 activates the signalling pathway TLR4-TRAM-TRIF-IRF3 to induce the production of type I IFN, which signals through the IFNAR1 receptor (**). Type I IFN stimulates the transcription of SARM1, and AIM2 via IRF3. SARM1 negatively regulates MyD88 and TRIF- governed inflammatory responses, the activation of the MAP kinases ERK and JNK, and the AIM2 inflammasome. In contrast, SARM1 is required for the activation of the MAP kinase p38, which controls the production of IL10. Kp52145 exploits IL10 to control inflammation. Absence of SARM1 impairs the intracellular survival of Kp52145, and *sarm1^-/-^* mice do control Kp52145 infection. Collectively, our findings illustrate the crucial role of SARM1 in *K. pneumoniae* immune evasion strategies.

The evidence of this study suggests that *K. pneumoniae* leverages the TIR-TIR interactions between SARM1, and MyD88 and TRIF to attenuate MyD88 and TRIF-dependent inflammatory responses. There are few examples of pathogens exploiting TIR-TIR interactions to blunt the activation of TLR-controlled signalling pathways (Askarian et al., 2014; Cirl et al., 2008; Coronas-Serna et al., 2020; Imbert et al., 2017; Xiong et al., 2019). However, and without exception, these pathogens deploy a prokaryotic protein containing the TIR domain into immune cells, whereas *K. pneumoniae* is the first pathogen hijacking an endogenous mammalian TIR-containing protein. SARM1TIR domain is more closely related to bacteria TIR proteins than to the other mammalian TIR containing adaptors (Zhang et al., 2011). This data highlights an evolutionary convergence between *K. pneumoniae* and the pathogens encoding TIR containing proteins to exploit TIR-TIR interactions to attenuate inflammation.

Another novel finding of our work is that the absence of SARM1 impairs *K. pneumoniae* induction of IL10. IL10 production complements the reduction in inflammation achieved by limiting the activation of TLR signalling due to TIR-TIR interactions following the recognition of *K. pneumoniae* by PRRs. The fact that neutralization of the cytokine enhances the clearance of the pathogen (Greenberger et al., 1995) illustrates the crucial role of IL10 in *K. pneumoniae* infection biology. How *K. pneumoniae* induces IL10 was unknown. Our data implicates p38 whose activation is fine-tuned by type I IFN elicited by *K. pneumoniae*. The absence of SARM1 perturbs the p38-type I IFN axis by increasing the levels of type I IFN, resulting in a reduction of *K. pneumoniae*-induced p38 activation with a concomitant reduction in the production of IL10. The regulatory connection between type I IFNs and IL10 has been described; however, and in contrast to our results, the data indicates that type I IFN signalling is needed to sustain IL10 production in macrophages following challenge with LPS or *Mycobacterium spp*. (Chang et al., 2007; McNab et al., 2014; Pattison et al., 2012). These results reflect the importance of type I IFNs levels in the host-pathogen interface although the consequences are context dependent.

Previous work has demonstrated the importance of IL1β-governed responses in host defence against *K. pneumoniae* (Cai et al., 2012). Not surprisingly, *K. pneumoniae* has evolved to blunt IL1β-mediated inflammation (Frank et al., 2013; Regueiro et al., 2011). However, there was no evidence indicating whether *K. pneumoniae* is able to counteract inflammasome activation to limit the production of IL1β. Here we demonstrate that SARM1 inhibits inflammasome activation by *K. pneumoniae*. The fact that SARM1 has recently been shown to inhibit NLRP3 activation (Carty et al., 2019), and that there are observations indication that *K. pneumoniae* may activate NLRP3 inflammasome (Hua et al., 2015; Willingham et al., 2009a), made plausible that SARM1 would inhibit NLRP3 activation in *K. pneunomiae* infected cells. However, this was not the case. Our data demonstrate that AIM2 is the inflammasome activated by *K. pneumoniae* that it is inhibited by SARM1. Components of the type I IFN signalling pathway were essential for the activation of AIM2 inflammasome by *K. pneumoniae*. This is similar to *Listeria monocytogenes* and *Francisella spp*, two other pathogens activating AIM2 (Fernandes-Alnemri et al., 2010; Henry et al., 2007; Jones et al., 2010; Man et al., 2015; Rathinam et al., 2010; Tsuchiya et al., 2010), reinforcing the link between type I IFN signalling and AIM2 inflammasome. However, cGAS-STING-IRF3- IFNAR1 signalling is necessary in the case of *Listeria* and *Francisella*-mediated activation of AIM2 (Fernandes-Alnemri et al., 2010; Hansen et al., 2014; Man et al., 2015; Rathinam et al., 2010), whereas TLR4-TRAM-TRIF-IRF3-IFNAR1 mediates *K. pneumoniae* induction of AIM2. This evidence uncovers the crucial role of IRF3-IFNAR1 signalling in the host-bacteria interface. Recently, we have demonstrated the importance of this hub to control *K. pneumoniae* infections (Ivin et al., 2017).

Mechanistically, *K. pneumoniae* triggered an association between SARM1 and AIM2, and the SARM1 TIR domain was sufficient to inhibit AIM2 activation. Altogether, our data is consistent with a model in which SARM1 directly targets AIM2 to supress the recruitment of ASC and ASC-speck formation, restraining the activation of caspase 1. To the best of our knowledge, *K. pneumoniae* is the first pathogen deploying a strategy to target directly AIM2 activation because the other known examples are based on reducing the activating signal (Ge et al., 2012; Ulland et al., 2010). On the other hand, the strategy deployed by *K. pneumoniae* is reminiscent of how cells avoid an excessive activation of AIM2 by leveraging two small proteins, p202 in mouse, and IFI16β in human cells, that impede AIM2-ASC complex formation (Wang et al., 2018; Yin et al., 2013).

It is intriguing that *K. pneumoniae* did not activate NLRP3 even in the background of *sarm1^-/-^* and *aim2^-/-^* cells. This is even more puzzling considering that *K. pneumoniae* increased the expression of NLRP3. Considering that the stimuli reported to activate NLRP3, such as ROS, are most likely also present in *K. pneumoniae*-infected cells, it is then tempting to speculate that *K*. *pneumoniae* has evolved mechanisms to blunt the activation of NLRP3. Future studies are warranted to uncover how *K. pneumoniae* inhibits NLRP3 activation.

Except in neurons, the levels of SARM1 are low in most cells types, including monocytes and macrophages (Doran et al., 2021; Uhlen et al., 2010), suggesting that SARM1 levels are under tight control. We provide evidence demonstrating that *K. pneumoniae* induced SARM1 in a type I IFN dependent manner via a TLR4-TRAM-TRIF-IRF3-IFNAR1 signalling pathway, hence placing SARM1 as an ISG. Likewise SARM1, type I IFNs are also conserved during evolution and appear in the first vertebrates (Secombes and Zou, 2017), suggesting that *K. pneumoniae* manipulates an ancient SARM1-type I IFNs axis to counteract the activation of host defences. It is interesting to note the complex interface between *K. pneumoniae* and type I IFN. On the one hand, TRIF-mediated type I IFN is essential for host defence against *K. pneumoniae* (Cai et al., 2009; Ivin et al., 2017) including the expression of IL1β as a result of AIM2 activation (this work), and to limit the production of IL10 (this work). On the other hand, *K. pneumoniae* exploits type I IFN to induce SARM1 to attenuate TRIF and AIM2 activation. This evidence supports the notion that there is a threshold of type I IFN levels that needs to be reached in order to exert a protective role whereas below this threshold type I IFNs promote *K. pneumoniae* infection. In this scenario, SARM1 is one of the breaks that *K. pneumoniae* uses to control type I IFNs. Future studies should investigate whether *K. pneumoniae* uses other means to control the levels of type I IFNs.

We were keen to identify the bacterial factor(s) mediating the expression of SARM1. *K. pneumoniae* does not encode any type III or IV secretion system or any of the toxins implicated in counteracting innate immunity, making then interesting to uncover how *K. pneumoniae* manipulates any host protein. Our results establish that the CPS and the LPS O-polysaccharide induced the expression of SARM1. This is in perfect agreement with the evidence demonstrating that both polysaccharides trigger the production of type I IFNs (Ivin et al., 2017). Importantly, these polysaccharides are required for *K. pneumoniae* survival in mice (pneumonia model) (Cortes et al., 2002; Lawlor et al., 2005; Tomas et al., 2015), underlining the importance of SARM1 induction as a *K. pneumoniae* virulence trait since this process is abrogated in these mutant strains. We recently demonstrated that both polysaccharides are crucial to reduce the SUMOylation of proteins to limit host defence responses involving type I IFN-regulated miRNAs of the *let-7* family (Sa-Pessoa et al., 2020). Altogether, this evidence underscores the role of *K. pneumoniae* CPS and LPS to hijack regulators of the host immune system, hence expanding their well-established role in *K. pneumoniae* stealth behaviour (Bengoechea and Sa Pessoa, 2019).

Previous work established that *K. pneumoniae* survives intracellularly in macrophages residing in the KCV (Cano et al., 2015). Here, we demonstrate that SARM1 is essential for the survival of *K. pneumoniae*. Mechanistically, absence of SARM1 impaired *K. pneumoniae*-induced activation of AKT which in turn limited the recruitment of Rab14 to the KCV resulting in the fusion of the KCV with lysosomes ((Cano et al., 2015) and this work). The reduction in AKT activation found in *sarm1^-/-^* cells also explains the reduction of phagocytosis of *K. pneumoniae* because previous studies have demonstrated conclusively the connection between PI3-K-AKT activation and phagocytosis of bacteria, including *K. pneumoniae*, and large particles (Cano et al., 2015; Schlam et al., 2015). It is intriguing to note that two other pathogens, *Salmonella typhimurium* and *M. tuberculosis,* also manipulate the PI3K-AKT-Rab14 pathway to arrest phagosome maturation (Kuijl et al., 2007; Kyei et al., 2006). It is then tempting to postulate that SARM1 may also play an important role in the intracellular survival of these two pathogens. If this is the case, the axis SARM1-PI3K-AKT-Rab14 will become one of the central nodes targeted by pathogens to take control over cellular functions. Current efforts of the laboratory are devoted to investigate this hypothesis.

The fact that *sarm1* deficient mice were more efficient at controlling *K. pneumoniae* infection than wild-type mice provides strong support to the notion that *K. pneumoniae* leverages SARM1 to counteract host defences. Somewhat unexpectedly considering our in vitro results, we found a reduction in the levels of type I IFN and ISGs in infected *sarm1* deficient mice. This might be due to the fact that type I IFNs and ISGs are produced early during *K. pneumoniae* infection (Ivin et al., 2017). Nonetheless, it is important to realize that type I IFN signalling is essential to control *K. pneumoniae* infections (Ivin et al., 2017). On the other hand, the in vivo data support that *K. pneumoniae* exploits SARM1 to limit inflammatory cytokines and chemokines, and to produce IL10, mirroring the in vitro results. Interestingly, a wealth of evidence supports that this type of lung inflammatory environment is essential to clear *K. pneumoniae* infections (Bengoechea and Sa Pessoa, 2019). Therefore, it can be concluded that *K. pneumoniae* exploits SARM1 to modify the lung microenvironment to flourish. Mass cytometry analysis uncovered the presence of high numbers of alveolar macrophages, and neutrophils in *sarm1^-/-^* deficient mice. This is in good agreement with previous studies showing the importance of these cell types for the clearance of *K. pneumoniae* infections (Broug-Holub et al., 1997; Xiong et al., 2015; Xiong et al., 2016). Our profile analysis revealed subsets of neutrophils and interstitial macrophages only present in *sarm1^-/-^* infected mice. These cells expressed high levels of markers associated with immune activation further reinforcing the notion that the microenvironment in the absence of SARM1 supports *K. pneumoniae* clearance.

*K. pneumoniae* nosocomial infections are associated with high morbidity and mortality (Giske et al., 2008), and, worryingly, there is an increase in the number of community acquired infections (Lipworth et al., 2021; Magiorakos et al., 2013). Not surprisingly, the World Health Organization has singled out *K. pneumoniae* as a global threat to human health, and includes the pathogen among those for which new therapeutics are urgently needed. Our findings, including in vivo experiments probing a pre-clinical pneumonia mouse model, provide compelling evidence demonstrating that SARM1 is a target to boost human defence mechanisms against *K. pneumoniae*. Host-directed therapeutics aiming to interfere with host factors required by pathogens to counter the immune system are emerging as untapped opportunities that are urgently needed in the face of the global pandemic of antibiotic resistant infections. SARM1 is a druggable protein, and the crystal structure of the TIR domain of SARM1 is solved at 1.8 Å (Horsefield et al., 2019). This high-resolution structural information should facilitate the development of small-molecule inhibitors. Indeed, major efforts are underway to develop pharmacological approaches to inhibit SARM1 in the context of diseases with pathophysiological neuronal cell death (DiAntonio, 2019; Hughes et al., 2021). Based on the results of this study, we propose that these drugs shall show a beneficial effect to treat *K. pneumoniae* infections alone or as a synergistic add-on to antibiotic treatment. Future studies shall confirm whether this is the case.

## MATERIALS and METHODS

### Ethics statement

The experiments involving mice were approved by the Queen’s University Belfast’s Ethics Committee and conducted in accordance with the UK Home Office regulations (project licences PPL2778 and PPL2910) issued by the UK Home Office. Animals were randomized for interventions but researches processing the samples and analysing the data were aware which intervention group corresponded to which cohort of animals.

### Bacterial strains and growth conditions

Kp52145 is a clinical isolate (serotype O1:K2) previously described (Lery et al., 2014; Nassif et al., 1989). The *cps* mutant strain, 52145-Δ*manC*, the mutant lacking the LPS O-polysaccharide, 52145-Δ*glf*, and the double mutant lacking the CPS and the LPS O-polysaccharide, 52145- *wca_k2_*- Δ*glf,* are isogenic strains of Kp52145 and they have been described previously (Kidd et al., 2017; Sa-Pessoa et al., 2020). Strain 52145-Δ*glf* expresses similar levels of CPS than the wild-type strain (Sa-Pessoa et al., 2020). Bacteria were grown in 5 ml Luria-Bertani (LB) medium at 37 °C on an orbital shaker (180 rpm), and where appropriate, antibiotics were added to the growth medium at the following concentration: carbenicllin, 50 μg/ml; chloramphenicol, 25 μg/ml.

### Mammalian cells and cell culture

iBMDMs cells from wild-type (WT), *tlr4*^−/−^, *myd88*^−/−^, and *tram^-/-^trif^-/-^* mice on a C57BL/6 background were obtained from BEI Resources (NIAID, NIH) (repository numbers NR-9456, NR- 9458, NR-15633, and NR-9568, respectively). *Il-10*^-/-^ and *irf3*^-/-^ iBMDMs were described previously (Bartholomew et al., 2019; Ivin et al., 2017). Additional iBMDMs were generated as previously described (Sa-Pessoa et al., 2020). Briefly, tibias and femurs from C57BL/6, *ifnar1* , *sarm1*^-/-^, *Sarm1^em1.1Tf^*, *Sarm1^FLAG^, aim2*^-/-^, *nlrp3*^-/-^, *casp-1*^-/-^, *asc*^-/-^, and *gsdmd*^-/-^ were removed using sterile techniques, and the bone marrow was flushed with fresh medium. To obtain macrophages, cells were plated in Dulbecco’s modified Eagle’s medium (DMEM) supplemented with 20% filtered L929 cell supernatant (a source of macrophage colony-stimulating factor) and maintained at 37°C in a humidified atmosphere of 5% CO2. Medium was replaced with fresh supplemental medium after 1 day. Immortalization of BMDMs was performed after 5 days by exposing them for 24 h to the J2 CRE virus (carrying v-myc and v-Raf/v-Mil oncogenes, kindly donated by Avinash R. Shenoy, Imperial College London). This step was repeated 2cdays later (day 7), followed by continuous culture in DMEM supplemented with 20% (vol/vol) filtered L929 cell supernatant for 4 to 6cweeks. The presence of a homogeneous population of macrophages was accessed by flow cytometry using antibodies for CD11b (clone M1/70; catalog number 17-0112-82; eBioscience) and CD11c (clone N418; catalog number 48-0114-82; eBioscience). Retroviral transduction of SARM1 in *sarm1*^-/-^ cells was done as previously described (Carty et al., 2006; Carty et al., 2019). iBMDMs and BMDMs were grown in DMEM (catalog number 41965; Gibco) supplemented with heat-inactivated fetal calf serum, 100 U/ml penicillin, and 0.1 mg/ml streptomycin (Gibco) at 37°C in a humidified 5% CO2 incubator. Cells were routinely tested for *Mycoplasma* contamination. Cells were seeded a density of 2 ×10^4^ cells/well in 24-well plates, 5 ×10^5^ cells/well in 12-well plates, and 2 ×10^6^ cells/well in 6-well plates.

### Infection conditions

Overnight bacterial cultures were refreshed 1/10 into a new tube containing 4.5 mL of fresh LB. After 2.5 h at 37°C, bacteria were pelleted (2500× g, 20 min, 22°C), resuspended in PBS and adjusted to an optical density of 1.0 at 600 nm (5 x 10^8^ CFU/ml). Infections were performed using a multiplicity of infection (MOI) of 100 bacteria per cell in a 1 ml volume. Synchronization of the infection was performed by centrifugation (200 x g for 5 min). For incubation times longer than 30 min, cells were washed and 1 ml of fresh medium containing gentamycin (100 μg/ml) was added to the wells to kill extracellular bacteria. Medium containing gentamycin was kept until the end of the experiment. Infections were performed one day after seeding the cells in the same medium used to maintain the cell line without antibiotics. Infected cells were incubated at 37°C in a humidified 5% CO2 incubator.

### siRNA experiments

For transfection of siRNAs, 2×10^4^ iBMDMs (6-well plates) were transfected in suspension with 20 nM siRNA using Lipofectamine RNAiMAX (Invitrogen) in 200 μ Opti-MEM I (ThermoFisher).AllStars negative-control siRNA (Qiagen) or ON-TARGET plus SMART pool siRNA targeting AIM2 (no. L-044968-01-0020; Dharmacon) and SARM1 (no. L-041633-01-0005; Dharmacon) were used to transfect cells. The macrophages were infected 16 h post transfection. Efficiency of transfection was confirmed by RT-qPCR analysis of duplicate samples from three independent transfections by normalizing to the hypoxanthine phosphoribosyltransferase 1 (*hprt*) gene and comparing gene expression in the knockdown sample with the AllStars negative control. Primers are listed in Table S3.

### Inhibitors, recombinant cytokines, and blocking antibodies

The NLRP3 inhibitor MCC950 ([vehicle solution DMSO], 10 μM CAS 256373-96-3 – Calbiochem, Sigma), and the caspase 1 inhibitor YVAD ([vehicle solution DMSO], 10 μM CAS 256373-96 Sigma) were added 2 h before infection to the cells. Recombinant mouse IL-10 ([vehicle solution water] 1ng/ml, Biolegend) was added overnight before infection. The p38 inhibitor SB203580 ([vehicle solution DMSO], 10 μM, Sigma) was added 2 h before infection. The mouse anti-IFNAR1 receptor antibody (clone MAR1-5A3 [vehicle solution water] 5 ng/ml, BioXcell) was added overnight before infection. All these reagents were kept for the duration of the experiment.

### RNA isolation and RT-qPCR

Infections were performed in 6-well plates. Cells were washed three times with pre-warmed sterile PBS, and total RNA was extracted from the cells in 1 ml of TRIzol reagent (Ambion) according to the manufacturer’s instructions. Extracted RNA was treated with DNase I (Roche) and precipitated with sodium acetate (Ambion) and ethanol. RNA was quantified using a Nanovue Plus spectrophotometer (GE Healthcare Life Sciences). cDNA was generated by retrotranscription of 1g of total RNA using M-MLV reverse transcriptase (Invitrogen) and random primers (Invitrogen). Two duplicates were generated from each sample. Ten nanograms of cDNA were used as a template in a 5- l reaction mixture from a KAPA SYBR FAST qPCR kit (Kapa Biosystems). Primers used are listed in table S3. RT-qPCR was performed using a Rotor-Gene Q (Qiagen) with the following thermocycling conditions: 95°C for 3 min for hot-start polymerase activation, followed by 40 cycles of 95°C for 5 s and 60°C for 20 s. Fluorescence of SYBR green dye was measured at 510 nm. Relative quantities of mRNAs were obtained using the ΔΔC_T_ method by using hypoxanthine phosphoribosyltransferase 1 (*hprt*) gene normalization.

### Immunoblots

Macrophages were seeded in 6-well plates for 24 h before infection. Cell lysates were prepared in lysis buffer (1x SDS Sample Buffer, 62.5 mM Tris-HCl pH 6.8, 2% w/v SDS, 10% glycerol, 50 mM DTT, 0.01% w/v bromophenol blue). Proteins were resolved on 8, 10 or 12% SDS-PAGE gels and electroblotted onto nitrocellulose membranes. Membranes were blocked with 3% (wt/vol) bovine serum albumin in TBS-Tween (TBST), and specific antibodies were used to detect protein using chemiluminescence reagents and a G:BOX Chemi XRQ chemiluminescence imager (Syngene).

The following antibodies were used: anti-IL-1β (anti-goat, 1:1000; # AF-401-NA, R&D Systems), anti-caspase-1 (anti-rabbit, 1:1000; #24232, Cell Signaling), anti-AIM2 (anti-rabbit, 1:1000; sc- 515895, Santa Cruz), anti-NLRP3 (anti-mouse, 1:1000; #15101, Cell Signaling), anti-Gasdermin-D (anti-rabbit, 1:1000; #93709, Cell Signaling), anti-Viperin (anti-rabbit, 1:1000 # NBP2-03971, Novus Biologicals), anti-ISG15 (1:1000; #9636, Cell Signaling), anti-phospho-STAT3 (anti-rabbit, 1:1,000; #9145, Cell Signaling), anti- κBα (anti-rabbit, 1:1,000; #4814, Cell Signaling), anti- phospho-Iκ Bα (Ser32) (anti-goat, 1:1,000; #9246, Cell Signaling), anti-phospho-AKT1/2/3 (Ser 473) (anti-rabbit, 1:1000; sc-33437, Santa Cruz), anti-phospho-IKKα/β (Ser176/180)(16A6) (anti- rabbit, 1:1000; #2697,Cell Signaling), anti-phospho-IRF3 (Ser 396) (anti-rabbit, 1:1000; #4947, Cell Signaling), anti-phospho-p-TBK-1/NAK (Ser172) (D52C2) (anti-rabbit, 1:1000; #5483, Cell Signaling), anti-phospho-JNK (anti-rabbit, 1:1000; #9251S, Cell Signaling), anti-phospho-ERK (anti-rabbit, 1:1000; #9101, Cell Signaling), anti-phospho-p38 MAPK (Thr180/Tyr182) (D3F9) (anti-rabbit, 1:1000; #4511, Cell Signaling), anti-SARM1 (anti-chicken, 1:70; generated by Icosagen by immunizing chicken with the TIR domain of human SARM1), anti-Flag M2 (1 μ Sigma F3165), anti-HA (1:1000, Santa Cruz Biotechnology sc-805). Immunoreactive bands were visualized by incubation with HRP-conjugated IgG Secondary antibody (anti-goat, 1:5000; # HAF017, R&D Systems, goat anti-rabbit, 1:5000; #170-6515, Bio Rad, goat anti-mouse, 1:5000; #6516, Bio-Rad). To ensure that equal amounts of proteins were loaded, blots were re-probed with α-tubulin (1:3000; #T9026, Sigma- Aldrich) or β-actin (anti-mouse, 1:1000; sc-130065, Santa Cruz). To detect multiple proteins, membranes were re-probed after stripping of previously used antibodies using a pH 2.2 glycine-HCl/SDS buffer.

### Processing cell free supernatants for inflammasome studies

iBMDMs were seeded in 6 wells plates and were infected 24 h later . At the indicated time points, the plates were centrifuged at 200xg for 5 min. at room temperature, and the supernatants were transferred to microcentrifuge tubes and placed on ice. The cells were lysed in 80 μl of Laemmeli buffer with β-mercaptoethanol (1 in 19 ratio), collected in a microcentrifuge tube and stored at -20°C. The supernatants were processed by adding 9 μl of StrataClean Resin, hydroxylated silica particles (Cat. 400714) per 1 ml of supernatant. The samples were homogenized in vortex for 1 min, and were centrifuged at 9000 x g for 2 min. The supernatant was discarded, and the pellets were suspended in 40 μl of Laemmli buffer and transferred to filtered columns within collection tubes. The columns were centrifuged at 8,000 x g at RT for 1 min, and the eluate collected. The samples were boiled for 5 min in heat block at 95°C and loaded for western blot analysis.

### Enzyme-linked immunosorbent assay (ELISA), and cytokine measurement

Infections were performed in 12-well plates. Supernatants from infected cells were collected at the indicated time points in the figure legends, and spun down at 12,000 x g for 5 min to remove any debris. TNF-α (#900-K54), IL-1β (#900-K47), IL-10 (#900-K53) and IP-10 (CXCL10) (#250-16) in the supernatants were quantified using ABTS ELISA Development Kit (PeproTech) according to the manufacturer’s instructions. All experiments were performed in duplicate, and three independent experiments were conducted.

For quantification of type I IFN (INF-α/β) in the supernatants of iBMDMs, cells were infected for 16 h, and supernatants were collected. Murine type I IFNs were detected using B16-Blue IFN-α/β reporter cells (Invivogen) which carry an SEAP reporter gene under the control of the IFN-α/β- of inducible ISG54 promoter and that have an inactivation of the IFN-γ receptor. Supernatants from iBMDM cells were incubated with the reporter cell line, and levels of SEAP in the supernatants were determined using the detection medium QUANTI-Blue (Invivogen) after 24 h as per the manufacturer’s instructions using recombinant mouse IFN-β (PBL Assay Science, catalogue number 12401-1) as a standard. Experiments were run in duplicates and repeated at least three times. Results are expressed as OD at 655 nm.

### Detection of ASC specks formation by flow cytometry

To detect ASC speck formation by flow cytometry, we adapted the protocol described by Sester and colleagues (Sester et al., 2015). Cells were harvested from 6-wells plates with ice-cold PBS, centrifuged at 1,000 x g for 5 min, and resuspended in 1 ml ice-cold PBS. Samples were then fixed by the drop wise addition of 4 ml ice-cold molecular grade ethanol while vortexing. After 15 min, cells were pelleted by centrifugation at 600 x g for 10 min, supernatants gently removed and pellets suspended in 250 μl ASC speck buffer (ASB, PBS/0.1% sodium azide, 0.1% BSA, 1.5% FCS) containing 1 μl Fc block anti-CD16/CD32 (2.4G2, BD Biosciences) for 20 min. To stain ASC specks, 0.2 μl anti-ASC (Cat# sc-22514R, Santa Cruz) in 50 μl ASB buffer were added to the samples, and incubated for 90 min at room temperature. The cells were washed with 1 ml ASB, and the re suspended in 50 μl ASB containing 0.1 μl Alexa 488 goat anti-rabbit IgG (H+L) (Molecular Probes). After 45 min, cells were washed with 1 ml ASB, and re suspended in 500 μl ASB. Samples were processed on a BD FACS Canto and analyzed using FlowJo X (Tree Star) software and graphical representation.

### AIM2 reconstitution in HEK cells

HEK293T cells were seeded at 2×10^5^ cells/well in 24-well plates and incubated overnight. The cells were transfected using Lipofectamine 2000 with plasmids expressing pro-IL-b-FLAG (50 ng), pro-Caspase-1-FLAG (10 ng), ASC-FLAG (1 ng), HA-AIM2 (50 ng) and 10, 50 or 100 ng of pdlNotInPkMCSR FLAG SARM1, FLAG SARM1 TIR, FLAG SARM1 ΔTIR or pdlNotInPkMCSR empty vector control. Medium was replaced 24 h after transfection and supernatants were collected 16 h after media change. Quantification of secreted murine IL-1β was performed using ELISA (R&D). Cells were lysed with RIPA buffer and subjected to immunoblotting by using anti-HA or anti-FLAG antibodies for the detection of AIM2 and SARM/SARM TIR/ SARM ΔTIR expression.

### Coimmunoprecipiration analysis

iBMDMs were seeded onto 6-wells plates (8×10^5^ cells/well). Cells were transfected the following day with 1 μg of MyD88-HA or TRIF-HA plasmids (Carty et al, 2006) diluted in 200 μl of opti-MEM (Gibco) using 6 μ of Lipofectamine 2000 (Invitrogen). Transfected cells were infected 20 h post transfection at a MOI of 100. After 1 h of contact, media was replaced by media containing gentamicin (100 μg/ml), and cells were collected at 3 h and lysed in RIPA buffer containing: 50 mM Tris-HCl, pH 7.2, 0.15 M NaCl, 0.1% SDS, 1% Sodium Deoxycholate, 1% Triton X-100 and proteinase inhibitors: 1 mM PMSF and halt protease inhibitor cocktail (ThermoFisher Scientific, catalogue number 78430). The whole cell lysates were centrifuged at 10,000 ×g for 20 minutes at 4°C. The supernatants were transferred to a new tube and the pellets were kept to probe the input. Whole cell lysates were incubated with 1 μg FLAG (Sigma, F3165) or normal mouse IgG (Santa Cruz, c-2025) antibodies for 2 hours at 4 °C in a rotary wheel mixer. Protein A/G Plus agarose suspension (Santa Cruz # sc-2003) was added to the whole cells lysates suspension and incubated at 4 °C on a rotary mixer overnight. The suspension was centrifuged at 1,000 ×g for 4 min at 4 °C and the supernatant was aspirated and discarded. Pellets were washed 2 times with RIPA buffer, suspended in 40 μl of 2 x electrophoresis sample buffer (Laemmli buffer) and boiled for 5 min at 95°C.

### Adhesion, phagocytosis and intracellular survival

iBMDMs were seeded in 12-well plates approximately 16 h before infection. Infections were performed as previously described. To enumerate the number of bacteria adhered to macrophages, after 30 min of contact cells were washed twice with PBS, and they were lysed in 300 μl of 0.1% (wt/vol) saponin in PBS for 5 min at 37°C. Serial dilutions were plated in LB and the following day bacterial CFUs were counted. Results are expressed as CFU per ml. To determine the number of bacteria phagocytosed by the cells, after 30 min of contact, cells were washed once with PBS and fresh medium containing gentamycin (100 μg/ml) was added to the wells. After 30 min, cells were washed three times with PBS, and lysed with saponin. Samples were serially diluted in PBS and plated in PBS. After 24 h incubation at 37°C, CFUs were counted and results expressed as CFUs per ml. To assess intracellular survival, 4 h after the addition of gentamycin, cells were washed three times with PBS and lysed with saponin. Serial dilutions were plated on LB to quantify the number of intracellular bacteria. Results are expressed as % of survival (CFUs at 4 h versus 1 h in *sarm1*^-/-^ cells normalized to the results obtained in wild-type macrophages set to 100%). All experiments were carried out with triplicate samples on at least five independent occasions.

### Assessment of the colocalization of the KCV with cellular markers

The protocol was adapted from (Cano et al., 2015). Briefly, wild-type and *sarm1*^-/-^ iBMDMs (2×10^4^ per well) were grown on 13 mm circular coverslips in 24-well plates and were infected with Kp52145 harbouring pFPV25.1Cm (March et al., 2013). After 30 min of contact the coverslips were washed with PBS and gentamycin (100μg/ml in DMEM medium) was added to kill extracellular bacteria

#### (i) Staining of lysosomes

Cresyl violet acetate salt (Sigma) was used to label lysosomes (Ostrowski et al., 2016). Cresyl violet in fresh medium (5µM) was added to the cells 12 min before fixing the cells. The residual fluid marker was removed by washing the cells three times with PBS, followed by fixation (4% paraformaldehyde in PBS pH 7.4 for 20 min at room temperature). Coverslips were mounted with ProLong™ Gold antifade mountant (Invitrogen). Coverslips were visualised on the Leica SP8 Confocal microscope within 24 h after fixing. To determine the percentage of bacteria that co localized with cresyl violet, bacteria located inside a minimum of 100 infected cells were analysed in each experiment. Experiments were carried out in duplicate in three independent occasions.

#### (ii) Rab14 staining

At the indicated time points post infection, coverslips were washed with PBS and permeabilized with 0.1% (w/v) saponin (Sigma) in PBS for 30 minutes. Coverslips were then incubated for 120 minutes with anti-Rab14 (4 µg/ml in 0.1% (v/v) horse serum (Gibco), 0.1% (w/v) saponin in PBS; clone D-5, murine IgG1, sc-271401, Santa Cruz Biotechnologies), washed with PBS, followed by a 45 minutes incubation with anti-mouse IgG H&L labelled with AlexaFluor 647 (10 µg/ml in 0.1% (v/v) horse serum (Gibco), 0.1% (w/v) saponin in PBS, polyclonal, donkey IgG, ab150111, Abcam). Coverslips were washed with PBS, and then incubated with anti-Lamp1 (1 µg/ml in 0.1% (v/v) horse serum (Gibco), 0.1% (w/v) saponin in PBS, clone 1D4B, rat IgG2a, sc-19992, Santa Cruz Biotechnologies) for 20 min, washed with PBS, and incubated for 20 minutes with anti-rat IgG H&L labelled with AlexaFluor 568 (10 µg/ml in 0.1% (v/v) horse serum (Gibco), 0.1% (w/v) saponin in PBS, polyclonal, goat IgG, A11077, Life Technologies). Coverslips were mounted in microscope slides with ProLong Gold antifade mountant (Invitrogen), and visualised on a TCS-SP5 inverted microscope (Leica Biosystems). To determine the percentage of the Lamp1 positive KCV that co localized with Rab14, KCVs of at least 100 infected cells from three independent experiments were analysed.

### Intranasal murine infection model

Infections were performed as previously described (Ivin et al., 2017). Briefly, 8- to 12-week-old C57BL/6 mice (Charles River), *sarm1^-/-^*, B6.129X1-Sarm1tm1Aidi/J (The Jackson Laboratory, and bred at Queen’s University Belfast), *Sarm1^em1.1Tft^* (Doran et al., 2021) of both sexes were infected intranasally with ∼3 × 10^5^ Kp52145 in 30 μl PBS. Non-infected mice were mock infected with 30 μl sterile PBS. The number of mice per group are indicated in the figure legends. 24 h post infection, mice were euthanized using a Schedule 1 method according to UK Home Office approved protocols. For those mice used for mass cytometry analysis, 16 hours post infection, they were dosed intraperitoneally with 500 µg of monensin (Sigma) for intracellular cytokine staining.

Left lung samples from infected and uninfected control mice were immersed in 1 ml of RNA stabilisation solution (50% [w/v] ammonium sulphate, 2.9% [v/v] 0.5M ethylenediaminetetraacetic acid, 1.8% [v/v] 1 M sodium citrate) on ice and then stored at 4°C for at least 24 h prior to RNA extraction. Samples were homogenized in 1 ml ice-cold TRIzol (Ambion) using a VDI 12 tissue homogenizer (VWR). RNA was extracted according to the manufacturer’s instructions extraction, and cDNA was generated by retrotranscription of 1 μg of total RNA using M-MLV reverse transcriptase (Invitrogen) and random primers (Invitrogen). RT-qPCR analysis was undertaken using the KAPA SYBR FAST qPCR Kit, oligonucleotide primers as described in the in vitro protocol, and Rotor-Gene Q (Qiagen). Thermal cycling conditions were as follows: 95°C for 3 min for enzyme activation, 40 cycles of denaturation at 95°C for 10 s and annealing at 60°C for 20 s. Each cDNA sample was tested in duplicate, and relative mRNA quantity was determined by the comparative threshold cycle (ΔΔC_T_) method using hypoxanthine phosphoribosyltransferase 1 (m*hprt*) gene normalisation.

Right lung, spleen and liver samples from infected mice were immersed in 1 ml sterile PBS on ice and processed for quantitative bacterial culture immediately. Samples were homogenised with a Precellys Evolution tissue homogenizer (Bertin Instruments), using 1.4 mm ceramic (zirconium oxide) beads at 4500 rpm for 7 cycles of 10 seconds, with a 10-second pause between each cycle. Homogenates were serially diluted in sterile PBS and plated onto *Salmonella-Shigella* agar (Sigma), and the colonies were enumerated after overnight incubation at 37°C. Data were expressed as CFUs per gr of tissue.

### Mass cytometry

#### (i) Generation of metal-labelled antibodies

Carrier protein and glycerol-free antibodies were labelled with lanthanide isotopes using Maxpar X8 Antibody Labelling Kits (Fluidigm) according to the manufacturer’s instructions. Briefly, X8 polymer was loaded with the lanthanide isotype in L-buffer, and the metal-loaded polymer purified and washed in C-buffer using an Amicon Ultra-0.5 centrifugal filter unit with 3kDa cutoff (Millipore-Sigma). At the same time, the antibody was reduced with 4 mM tris(2- carboxyethyl)phosphine hydrochloride (TCEP) solution in R-buffer, and purified in C-buffer, using an Amicon Ultra-0.5 centrifugal filter unit with 50kDa cut-off (Millipore-Sigma).

Both the lanthanide-loaded polymer and the partially reduced antibody were mixed and incubated at 37 °C for 90 minutes. Once the incubation was completed, the conjugated antibody was washed several times with W-buffer using an Amicon Ultra-0.5 centrifugal filter unit with 50kDa cut-off (Millipore-Sigma), and quantified using a NanoDrop spectrophotometer (280 nm). The antibody was finally resuspended in antibody stabilizer PBS supplemented with 0.05% sodium azide at a final concentration of 0.5 mg/mL and stored at 4 °C.

#### (ii) Mass cytometry staining and acquisition

Mice lungs were aseptically collected in PBS and homogenized with a handheld homogenizer. Single-cell suspensions were obtained by flushing the samples through 70 µM strainer, incubated with nuclease (Pierce). Red blood cells were lysed with ACK buffer, and samples stained, according to manufacturer’s instructions. Briefly, cell suspensions were first incubated with 1 µM of 103Rh for live/dead discrimination, and later with cell surface metal-labelled antibodies, prepared in Maxpar Cell Staining Buffer (CSB; Fluidigm), for 30 minutes at room temperature. Cells were washed with CSB, fixed and permeabilized with Maxpar Fix I buffer (Fluidigm) for 10 minutes at room temperature, washed with 2 volumes ofMaxpar Perm-S buffer (Fluidigm), and incubated with metal-labelled antibodies for intracellular markers, prepared in Maxpar Perm-S buffer, for 30 minutes at room temperature. The list of antibodies used is shown in Table S1. Finally, samples were washed with CSB, incubated 10 minutes at room temperature with a 2% paraformaldehyde solution, washed once more with CSB, and left at 4 °C in Maxpar Fix and Perm buffer (Fluidigm) with 125 nM Cell-ID™ Intercalator Ir (Fluidigm) until acquisition. Samples were acquired between 12 and 48 hours after staining. Right before acquisition, cells were washed with CSB, followed by Maxpar Cell Acquisition Solution (CAS; Fluidigm). Cells were resuspended in CAS with 1 mM EDTA to a final concentration of 1×10^6^ cells/mL, flushed through a 35 µM strainer, and supplemented with 1/10 v/v EQ Four Element Calibration Beads (Fluidigm). Mass cytometry was performed using a Helios CyTOF instrument (Fluidigm) operated with software v7.0.8493. The CyTOF instrument was started, tuned, and cleaned according to the manufacturer’s protocol, and samples acquired with an injection speed of 30 µL/minute.

#### (iii) Mass cytometry data analysis

Data was exported as flow-cytometry FCS file format, and pre-processed with CyTOF software (v6.7.1014; Fluidigm) for normalization. Processed files were uploaded to the Cytobank platform (https://www.cytobank.org/) for initial gating (Gaussian parameters and cells/beads, live/dead and singlets/doublets discriminations). CD45^+^ populations were gated and exported in FCS file format an analysed with RStudio software (https://www.rstudio.com/) and cytofkit package (https://github.com/JinmiaoChenLab/cytofkit) for Phenograph clustering using the following parameters: 10.000 cells/sample, cytofAsinh as transformation Method, Phenograph as cluster method, k equal to 30 as Rphenograph, tsne as visualization method, a seed of 42.

### Statistical analysis

Statistical analyses were performed using one-way analysis of variance (ANOVA) with Bonferroni corrections, the one-tailed t test, or, when the requirements were not met, the Mann-Whitney U test. P values of <0.05 were considered statistically significant. Normality and equal variance assumptions were tested with the Kolmogorov-Smirnov test and the Brown-Forsythe test, respectively. All analyses were performed using GraphPad Prism for Windows (version 9.1.0) software.

## ACKNOWLEDGEMENTS

We thank the members of the J.A.B. and A.G.B. laboratories for their thoughtful discussions and support with this project. This work was supported by Biotechnology and Biological Sciences Research Council (BBSRC, BB/T001976/1) funds to J.A.B., a joint BBSRC-Science Foundation Ireland (SFI) grant to J.A.B. (BB/P020194/1) and A.G.B. (17/BBSRC/3414), and by SFI (16/IA/4376) to A.G.B. The mass cytometry equipment at Queen’s University Belfast was funded by an institutional grant, and the technical support was provided by the Wellcome-Wolfson Institute for Experimental Medicine.

## AUTHOR CONTRIBUTIONS

Conceptualization, J.A.B. and A.G.B; Investigation, C.F., J. sP., R. CG., L.G. B.G., J.L.I., M.C. and A.D. Resources, R.S., R.J.I, A.K.; Funding acquisition, J.A.B. and A.G.B.; Writing original draft, J.A.B. A.G.B. A.K. C.F.; Writing-Review and Editing M.C., R.J.I, A.K. J.A.B. and A.G.B. Supervision, J.A.B., and A.G.B.

## SUPPLEMENTARY FIGURE LEGENDS

**Figure S1. SARM1 negatively regulates *K. pneumoniae*-induced inflammation.**

A. ELISA of TNFα, IL1β, CXCL10 secreted by wild-type (WT) and *sarm1* BMDMs non-infected (ni) or infected with Kp52145 for 6 and 16 h. After 1 h contact, the medium was replaced with medium containing gentamicin (100 µg/ml) to kill extracellular bacteria.

B. Efficiency of transfection of SARM siRNA (siSARM) in wild-type iBMDMs. mRNA levels were assessed 16 h post transfection as fold change against control non-silencing agents AllStars (siAS).

C. ELISA of IL1β, TNFα, and CXCL10 secreted by wild-type (WT) macrophages transfected with All Stars siRNA control (siAS), or SARM1 siRNA (siSARM) non-infected (ni) or infected with Kp52145 for 16 h. After 1 h contact, the medium was replaced with medium containing gentamicin (100 μg/ml) to kill extracellular bacteria.

D. ELISA of IL1β, TNFα, and CXCL10 secreted by wild-type (WT) and *Sarm1em1.1Tft* macrophages non-infected (ni) or infected with Kp52145 for 6 and 16 h. After 1 h contact, the medium was replaced with medium containing gentamicin (100 μg/ml) to kill extracellular bacteria.

In panels A, C, and D, values are presented as the mean ± SD of three independent experiments measured in duplicate. ****P ≤ 0.0001; **P≤ 0.01; *P ≤ 0.05; for the indicated comparisons using one way-ANOVA with Bonferroni contrast for multiple comparisons test. In panel B, **P≤ 0.01 using unpaired t test.

**Figure S2. *K. pneumoniae* induction of IL10 is controlled by p38 and it is negatively regulated by type I IFN.**

A. ELISA of IL10 secreted by wild-type macrophages non-infected (ni) or infected with Kp52145 (Kp) 16 h. Cells were treated with the p38 inhibitor SB202190 or DMSO vehicle control. After 1 h contact, the medium was replaced with medium containing gentamicin (100 µg/ml) to kill extracellular bacteria.

B. Efficiency of transfection of SARM1 siRNA (siSARM) in *il10^-/-^* macrophages. mRNA levels were assessed 16 h post transfection as fold change against control non-silencing agents AllStars (siAS).

C. Immunoblot analysis of phosphorylated p38 (P-p38), and tubulin levels in lysates of wild-type (WT) and *ifnar1^-/-^* macrophages non-infected (NI) or infected with Kp52145 for the indicated time points. After 1 h contact, the medium was replaced with medium containing gentamicin (100 µg/ml) to kill extracellular bacteria.

D. Immunoblot analysis of phosphorylated p38 (P-p38), and tubulin levels in lysates of wild-type (WT) and *tlr4^-/-^* macrophages non-infected (NI) or infected with Kp52145 for the indicated time points. After 1 h contact, the medium was replaced with medium containing gentamicin (100 µg/ml) to kill extracellular bacteria.

E. Immunoblot analysis of phosphorylated p38 (P-p38), and tubulin levels in lysates of wild-type (WT) and *tram^-/-^trif^-/-^* macrophages non-infected (NI) or infected with Kp52145 for the indicated time points. After 1 h contact, the medium was replaced with medium containing gentamicin (100 µg/ml) to kill extracellular bacteria.

F. *il10* mRNA levels were assessed by qPCR, in wild-type (WT), *tlr4^-/-^*, *tram^-/-^trif^-/-^*, and *ifnar1^-/-^* macrophages non-infected (ni) or infected with Kp52145 for 6 and 16 h. After 1 h contact, the medium was replaced with medium containing gentamicin (100 µg/ml) to kill extracellular bacteria. In panel A, values are presented as the mean ± SD of three independent experiments measured in duplicate. ****P ≤ 0.0001 for the indicated comparisons using one way-ANOVA with Bonferroni contrast for multiple comparisons test. In panel B, values are presented as the mean ± SD of three independent experiments measured in duplicate. **P ≤ 0.01 using unpaired t test. In panel F, values are presented as the mean ± SD of three independent experiments measured in duplicate. ****P ≤ 0.0001 for the comparison between infected knock-out and wild-type cells for 6 h; # P ≤ 0.0001 for the comparison between infected knock-out and wild-type cells for 16 h using one way-ANOVA with Bonferroni contrast for multiple comparisons test.

In panels C, D and E the images are representative of three independent experiments.

**Figure S3. *K. pneumoniae* does not activate NLRP3 inflammasome.**

A. ELISA of IL1β secreted by wild-type (WT), *asc^-/-^*, and *gsmd^-/-^* macrophages non-infected (ni) or infected with Kp52145 (Kp) for 16 h. After 1 h contact, the medium was replaced with medium containing gentamicin (100 µg/ml) to kill extracellular bacteria.

B. Immunoblot analysis of processed pro-IL1β macrophages (WT) and *asc^-/-^* and *gsmd^-/-^* macrophages non-infected or infected with Kp52145 for 16h. After 1 h contact, the medium was replaced with medium containing gentamicin (100 µg/ml) to kill extracellular bacteria.

C. ELISA of IL1β secreted by wild-type (WT) macrophages non-infected (ni) or infected with Kp52145 (Kp) for 6 and 16 h. Cells were treated with the NLRP3 inhibitor MC950 or DMSO vehicle control. After 1 h contact, the medium was replaced with medium containing gentamicin (100 µg/ml) to kill extracellular bacteria.

D. ELISA of IL1β secreted by wild-type (WT) and *nlrp3* macrophages non-infected (ni) or infected with Kp52145 for 6 and 16 h. After 1 h contact, the medium was replaced with medium containing gentamicin (100 µg/ml) to kill extracellular bacteria.

E. Immunoblot analysis of processed pro-IL1β, and β-actin levels in lysates of wild-type macrophages (WT) and *nlrp3^-/-^* macrophages non-infected or infected with Kp52145 for 16h. After 1 h contact, the medium was replaced with medium containing gentamicin (100 µg/ml) to kill extracellular bacteria.

F. Immunoblot analysis of NLRP3 and tubulin levels in lysates of wild-type macrophages (WT) and *nlrp3^-/-^* macrophages non-infected (NI) or infected with Kp52145 for the indicated time points. After 1 h contact, the medium was replaced with medium containing gentamicin (100 µg/ml) to kill extracellular bacteria.

G. Efficiency of transfection of AIM2 siRNA (siAIM2) in sarm1*^-/-^* macrophages. mRNA levels were assessed 16 h post transfection as fold change against control non-silencing agents AllStars (siAS).

In panels A, C and D values are presented as the mean ± SD of three independent experiments measured in duplicate. ****P ≤ 0.0001; ns, P > 0.05 for the indicated comparisons using one way- ANOVA with Bonferroni contrast for multiple comparisons test. In panel G, values are presented as the mean ± SD of three independent experiments measured in duplicate. **P≤ 0.01 using unpaired t test. In panels B, E and F, images are representative of three independent experiments.

**Figure S4. *K. pneumoniae* induction of AIM2 is TLR4-TRAM-TRIF-IRF3 dependent.**

A. *aim2* mRNA levels were assessed by qPCR, in wild-type (WT), *myd88^-/-^*, *tlr4^-/-^*, *tram^-/-^trif^-/-^*, and *irf31^-/-^* macrophages non-infected (ni) or infected with Kp52145 for 6 and 16 h. After 1 h contact, the medium was replaced with medium containing gentamicin (100 µg/ml) to kill extracellular bacteria.

B. Immunoblot analysis of AIM2 and β-actin levels in lysates of wild-type (WT), *tlr4* and *tram trif^-/-^* macrophages non-infected (NI) or infected with Kp52145 for the indicated time points. After 1 h contact, the medium was replaced with medium containing gentamicin (100 µg/ml) to kill extracellular bacteria.

C. ELISA of IL1β secreted by wild-type (WT), *tlr4* , *tram trif* and *ifnar1* macrophages non- infected (ni) or infected with Kp52145 for 6 and 16 h. After 1 h contact, the medium was replaced with medium containing gentamicin (100 µg/ml) to kill extracellular bacteria.

D. Immunoblot analysis of pro-IL1β and β-actin levels in lysates of wild-type (WT), and *tlr4* macrophages non-infected (NI) or infected with Kp52145 for the indicated time points. After 1 h contact, the medium was replaced with medium containing gentamicin (100 µg/ml) to kill extracellular bacteria.

In panels A and C, values are presented as the mean ± SD of three independent experiments measured in duplicate. # P ≤ 0.0001; ns, P > 0.05 for the comparison between knockout and wild- type cells at 6 or 16 h post infection using one way-ANOVA with Bonferroni contrast for multiple comparisons test. In panels B and D, images are representative of three independent experiments.

**Figure S5. Adhesion and phagocytosis of *K. pneumoniae* by *sarm1^-/-^* macrophages.**

A. Adhesion in wild-type (WT) and *sarm1^-/-^* macrophages. Cells were infected with Kp52145 for 30 min, wells were washed and bacteria were quantified by lysis, serial dilution and viable counting on LB agar plates.

B. Phagocytosis of Kp52145 by wild-type (WT) and *sarm1^-/-^* macrophages. Cells were infected for 30 min, wells were washed, and it was added medium containing gentamicin (100 µg/ml) to kill extracellular bacteria. After 30 min, cells were washed and bacteria were quantified by lysis, serial dilution and viable counting on LB agar plates.

In panels A and B, values are presented as the mean ± SD of three independent experiments measured in triplicate. * P ≤ 0.05; ns, P > 0.05 for the indicated comparisons using unpaired t test.

**Figure S6. Description of mouse immune populations following *K. pneumoniae* infection.**

A. PhenoGraph cluster analysis of immune populations in the lungs wild-type (WT), and *sarm1^-/-^* mice non-infected (ni) or infected intranasally with Kp52145 for 24. Graphs shows the combine results of all groups.

B. Heat map showing relative signal intensities of the indicated markers on the clusters identified in panel A. The heat map is coloured based on signal intensity of the indicated markers. Results are based on data from three mice per group.

C. PhenoGraph cluster analysis of immune populations in the lungs wild-type (WT), and *sarm1^-/-^* mice non-infected (ni) or infected intranasally with Kp52145 for 24. Each graph represents an individual mouse.

## REFERENCES

Arnold, I.C., Mathisen, S., Schulthess, J., Danne, C., Hegazy, A.N., and Powrie, F. (2016). CD11c+ monocyte/macrophages promote chronic Helicobacter hepaticus-induced intestinal inflammation through the production of IL-23. Mucosal immunology 9, 352–363.

Askarian, F., Van Sorge, N.M., Sangvik, M., Beasley, F.C., Henriksen, J.R., Sollid, J.U., Van Strijp, J.A., Nizet, V., and Johannessen, M. (2014). A Staphylococcus aureus TIR domain protein virulence factor blocks TLR2-mediated NF-κB signaling. Journal of innate immunity 6, 485–498.

Bartholomew, T.L., Kidd, T.J., Sa Pessoa, J., Conde Alvarez, R., and Bengoechea, J.A. (2019). 2-Hydroxylation of Acinetobacter baumannii Lipid A Contributes to Virulence. Infect Immun 87.

Belinda, L.W., Wei, W.X., Hanh, B.T., Lei, L.X., Bow, H., and Ling, D.J. (2008). SARM: a novel Toll-like receptor adaptor, is functionally conserved from arthropod to human. Mol Immunol 45, 1732–1742.

Bengoechea, J.A., and Sa Pessoa, J. (2019). Klebsiella pneumoniae infection biology: living to counteract host defences. FEMS Microbiol Rev 43, 123–144.

Bratkowski, M., Xie, T., Thayer, D.A., Lad, S., Mathur, P., Yang, Y.-S., Danko, G., Burdett, T.C., Danao, J., and Cantor, A. (2020). Structural and mechanistic regulation of the pro-degenerative NAD hydrolase SARM1. Cell Reports 32, 107999.

Broug-Holub, E., Toews, G.B., van Iwaarden, J.F., Strieter, R.M., Kunkel, S.L., Paine, R., 3rd, and Standiford, T.J. (1997). Alveolar macrophages are required for protective pulmonary defenses in murine Klebsiella pneumonia: elimination of alveolar macrophages increases neutrophil recruitment but decreases bacterial clearance and survival. Infect Immun 65, 1139–1146.

Cai, S., Batra, S., Shen, L., Wakamatsu, N., and Jeyaseelan, S. (2009). Both TRIF- and MyD88-dependent signaling contribute to host defense against pulmonary Klebsiella infection. J Immunol 183, 6629–6638.

Cai, S., Batra, S., Wakamatsu, N., Pacher, P., and Jeyaseelan, S. (2012). NLRC4 inflammasome-mediated production of IL-1beta modulates mucosal immunity in the lung against gram-negative bacterial infection. J Immunol 188, 5623–5635.

Cai, X., Chen, J., Xu, H., Liu, S., Jiang, Q.X., Halfmann, R., and Chen, Z.J. (2014). Prion-like polymerization underlies signal transduction in antiviral immune defense and inflammasome activation. Cell 156, 1207–1222.

Cano, V., March, C., Insua, J.L., Aguilo, N., Llobet, E., Moranta, D., Regueiro, V., Brennan, G.P., Millan-Lou, M.I., Martin, C., et al. (2015). Klebsiella pneumoniae survives within macrophages by avoiding delivery to lysosomes. Cel Microbiol 17, 1537–1560.

Carlsson, E., Ding, J.L., and Byrne, B. (2016). SARM modulates MyD88-mediated TLR activation through BB- loop dependent TIR-TIR interactions. Biochimica et Biophysica Acta (BBA)-Molecular Cell Research 1863, 244–253.

Carty, M., Goodbody, R., Schroder, M., Stack, J., Moynagh, P.N., and Bowie, A.G. (2006). The human adaptor SARM negatively regulates adaptor protein TRIF-dependent Toll-like receptor signaling. Nat Immunol 7, 1074–1081.

Carty, M., Kearney, J., Shanahan, K.A., Hams, E., Sugisawa, R., Connolly, D., Doran, C.G., Munoz-Wolf, N., Gurtler, C., Fitzgerald, K.A., et al. (2019). Cell Survival and Cytokine Release after Inflammasome Activation Is Regulated by the Toll-IL-1R Protein SARM. Immunity 50, 1412–1424 e1416.

Chang, E.Y., Guo, B., Doyle, S.E., and Cheng, G. (2007). Cutting edge: involvement of the type I IFN production and signaling pathway in lipopolysaccharide-induced IL-10 production. J Immunol 178, 6705–6709.

Cirl, C., Wieser, A., Yadav, M., Duerr, S., Schubert, S., Fischer, H., Stappert, D., Wantia, N., Rodriguez, N., Wagner, H., et al. (2008). Subversion of Toll-like receptor signaling by a unique family of bacterial Toll/interleukin-1 receptor domain-containing proteins. Nat Med 14, 399–406.

Coll, R.C., Robertson, A.A., Chae, J.J., Higgins, S.C., Munoz-Planillo, R., Inserra, M.C., Vetter, I., Dungan, L.S., Monks, B.G., Stutz, A., et al. (2015). A small-molecule inhibitor of the NLRP3 inflammasome for the treatment of inflammatory diseases. Nat Med 21, 248–255.

Coronas-Serna, J.M., Louche, A., Rodríguez-Escudero, M., Roussin, M., Imbert, P.R., Rodríguez-Escudero, I., Terradot, L., Molina, M., Gorvel, J.-P., and Cid, V.J. (2020). The TIR-domain containing effectors BtpA and BtpB from Brucella abortus impact NAD metabolism. PLoS pathogens 16, e1007979.

Cortes, G., Borrell, N., de Astorza, B., Gomez, C., Sauleda, J., and Alberti, S. (2002). Molecular analysis of the contribution of the capsular polysaccharide and the lipopolysaccharide O side chain to the virulence of Klebsiella pneumoniae in a murine model of pneumonia. Infect Immun 70, 2583–2590.

DiAntonio, A. (2019). Axon degeneration: mechanistic insights lead to therapeutic opportunities for the prevention and treatment of peripheral neuropathy. Pain 160 Suppl 1, S17–S22.

Dong, C., Davis, R.J., and Flavell, R.A. (2002). MAP kinases in the immune response. Ann Rev Immunol 20, 55–72.

Doran, C.G., Sugisawa, R., Carty, M., Roche, F., Fergus, C., Hokamp, K., Kelly, V.P., and Bowie, A.G. (2021). Next generation SARM1 knockout and epitope tagged CRISPR-Cas9-generated isogenic mice reveal that SARM1 does not participate in regulating nuclear transcription, despite confirmation of protein expression in macrophages. bioRxiv, 2021.2008.2025.457655.

Fernandes-Alnemri, T., Yu, J.-W., Datta, P., Wu, J., and Alnemri, E.S. (2009). AIM2 activates the inflammasome and cell death in response to cytoplasmic DNA. Nature 458, 509–513.

Fernandes-Alnemri, T., Yu, J.-W., Juliana, C., Solorzano, L., Kang, S., Wu, J., Datta, P., McCormick, M., Huang, L., and McDermott, E. (2010). The AIM2 inflammasome is critical for innate immunity to Francisella tularensis. Nat Immunol 11, 385–393.

Fitzgerald, K.A., McWhirter, S.M., Faia, K.L., Rowe, D.C., Latz, E., Golenbock, D.T., Coyle, A.J., Liao, S.-M., and Maniatis, T. (2003). IKKε and TBK1 are essential components of the IRF3 signaling pathway. Nat Immunol 4, 491–496.

Fornarino, S., Laval, G., Barreiro, L.B., Manry, J., Vasseur, E., and Quintana-Murci, L. (2011). Evolution of the TIR domain-containing adaptors in humans: swinging between constraint and adaptation. Mol Biol Evol 28, 3087–3097.

Frank, C.G., Reguerio, V., Rother, M., Moranta, D., Maeurer, A.P., Garmendia, J., Meyer, T.F., and Bengoechea, J.A. (2013). Klebsiella pneumoniae targets an EGF receptor-dependent pathway to subvert inflammation. Cel Microbiol 15, 1212–1233.

Ge, J., Gong, Y.N., Xu, Y., and Shao, F. (2012). Preventing bacterial DNA release and absent in melanoma 2 inflammasome activation by a Legionella effector functioning in membrane trafficking. Proc Natl Acad Sci U S A 109, 6193–6198.

Giske, C.G., Monnet, D.L., Cars, O., Carmeli, Y., and ReAct-Action on Antibiotic, R. (2008). Clinical and economic impact of common multidrug-resistant gram-negative bacilli. Antimicrob Agents Chemother 52, 813–821.

Greenberger, M.J., Strieter, R.M., Kunkel, S.L., Danforth, J.M., Goodman, R.E., and Standiford, T.J. (1995). Neutralization of IL-10 increases survival in a murine model of Klebsiella pneumonia. J Immunol 155, 722–729.

Gu, D., Dong, N., Zheng, Z., Lin, D., Huang, M., Wang, L., Chan, E.W., Shu, L., Yu, J., Zhang, R., et al. (2018). A fatal outbreak of ST11 carbapenem-resistant hypervirulent Klebsiella pneumoniae in a Chinese hospital: a molecular epidemiological study. Lancet Infectious diseases 18, 37–46.

Hansen, K., Prabakaran, T., Laustsen, A., Jorgensen, S.E., Rahbaek, S.H., Jensen, S.B., Nielsen, R., Leber, J.H., Decker, T., Horan, K.A., et al. (2014). Listeria monocytogenes induces IFNbeta expression through an IFI16-, cGAS- and STING-dependent pathway. EMBO J 33, 1654–1666.

Henry, T., Brotcke, A., Weiss, D.S., Thompson, L.J., and Monack, D.M. (2007). Type I interferon signaling is required for activation of the inflammasome during Francisella infection. J Exp Med 204, 987–994.

Holt, K.E., Wertheim, H., Zadoks, R.N., Baker, S., Whitehouse, C.A., Dance, D., Jenney, A., Connor, T.R., Hsu, L.Y., Severin, J., et al. (2015). Genomic analysis of diversity, population structure, virulence, and antimicrobial resistance in Klebsiella pneumoniae, an urgent threat to public health. Proc Natl Acad Sci U S A 112, E3574–3581.

Horsefield, S., Burdett, H., Zhang, X., Manik, M.K., Shi, Y., Chen, J., Qi, T., Gilley, J., Lai, J.S., Rank, M.X., et al. (2019). NAD(+) cleavage activity by animal and plant TIR domains in cell death pathways. Science 365, 793–799.

Hua, K.-F., Yang, F.-L., Chiu, H.-W., Chou, J.-C., Dong, W.-C., Lin, C.-N., Lin, C.-Y., Wang, J.-T., Li, L.-H., and Chiu, H.-W. (2015). Capsular polysaccharide is involved in NLRP3 inflammasome activation by Klebsiella pneumoniae serotype K1. Infect Immun 83, 3396–3409.

Hughes, R.O., Bosanac, T., Mao, X., Engber, T.M., DiAntonio, A., Milbrandt, J., Devraj, R., and Krauss, R. (2021). Small Molecule SARM1 Inhibitors Recapitulate the SARM1(-/-) Phenotype and Allow Recovery of a Metastable Pool of Axons Fated to Degenerate. Cell Rep 34, 108588.

Imbert, P.R., Louche, A., Luizet, J.B., Grandjean, T., Bigot, S., Wood, T.E., Gagné, S., Blanco, A., Wunderley, L., and Terradot, L. (2017). A Pseudomonas aeruginosa TIR effector mediates immune evasion by targeting UBAP 1 and TLR adaptors. EMBO J 36, 1869–1887.

Ivashkiv, L.B., and Donlin, L.T. (2014). Regulation of type I interferon responses. Nat Rev Immunol 14, 36–49.

Ivin, M., Dumigan, A., de Vasconcelos, F.N., Ebner, F., Borroni, M., Kavirayani, A., Przybyszewska, K.N., Ingram, R.J., Lienenklaus, S., Kalinke, U., et al. (2017). Natural killer cell-intrinsic type I IFN signaling controls Klebsiella pneumoniae growth during lung infection. PLoS pathogens 13, e1006696.

Jenner, R.G., and Young, R.A. (2005). Insights into host responses against pathogens from transcriptional profiling. Nat Rev Microbiol 3, 281–294.

Jones, J.W., Kayagaki, N., Broz, P., Henry, T., Newton, K., O’Rourke, K., Chan, S., Dong, J., Qu, Y., and Roose-Girma, M. (2010). Absent in melanoma 2 is required for innate immune recognition of Francisella tularensis. Proc Natl Acad Sci U S A 107, 9771–9776.

Kidd, T.J., Mills, G., Sa-Pessoa, J., Dumigan, A., Frank, C.G., Insua, J.L., Ingram, R., Hobley, L., and Bengoechea, J.A. (2017). A Klebsiella pneumoniae antibiotic resistance mechanism that subdues host defences and promotes virulence. EMBO Mol Med 9, 430–447.

Kuijl, C., Savage, N.D., Marsman, M., Tuin, A.W., Janssen, L., Egan, D.A., Ketema, M., van den Nieuwendijk, R., van den Eeden, S.J., Geluk, A., et al. (2007). Intracellular bacterial growth is controlled by a kinase network around PKB/AKT1. Nature 450, 725–730.

Kyei, G.B., Vergne, I., Chua, J., Roberts, E., Harris, J., Junutula, J.R., and Deretic, V. (2006). Rab14 is critical for maintenance of Mycobacterium tuberculosis phagosome maturation arrest. EMBO J 25, 5250–5259.

Lam, M.M.C., Wick, R.R., Wyres, K.L., Gorrie, C.L., Judd, L.M., Jenney, A.W.J., Brisse, S., and Holt, K.E. (2018). Genetic diversity, mobilisation and spread of the yersiniabactin-encoding mobile element ICEKp in Klebsiella pneumoniae populations. Microbial genomics.

Lawlor, M.S., Hsu, J., Rick, P.D., and Miller, V.L. (2005). Identification of Klebsiella pneumoniae virulence determinants using an intranasal infection model. Mol Microbiol 58, 1054–1073.

Lery, L.M., Frangeul, L., Tomas, A., Passet, V., Almeida, A.S., Bialek-Davenet, S., Barbe, V., Bengoechea, J.A., Sansonetti, P., Brisse, S., et al. (2014). Comparative analysis of Klebsiella pneumoniae genomes identifies a phospholipase D family protein as a novel virulence factor. BMC biology 12, 41–7007-7012-7041.

Levine, J.H., Simonds, E.F., Bendall, S.C., Davis, K.L., Amir el, A.D., Tadmor, M.D., Litvin, O., Fienberg, H.G., Jager, A., Zunder, E.R., et al. (2015). Data-Driven Phenotypic Dissection of AML Reveals Progenitor-like Cells that Correlate with Prognosis. Cell 162, 184–197.

Lewis, S.M., Treacher, D.F., Edgeworth, J., Mahalingam, G., Brown, C.S., Mare, T.A., Stacey, M., Beale, R., and Brown, K.A. (2015). Expression of CD11c and EMR2 on neutrophils: potential diagnostic biomarkers for sepsis and systemic inflammation. Clin Exp Immunol 182, 184–194.

Lipworth, S., Vihta, K.-D., Chau, K., Barker, L., George, S., Kavanagh, J., Davies, T., Vaughan, A., Andersson, M., Jeffery, K., et al. (2021). Ten-year longitudinal molecular epidemiology study of Escherichia coli and Klebsiella species bloodstream infections in Oxfordshire, UK. Genome Medicine 13, 144.

Lu, A., Magupalli, V.G., Ruan, J., Yin, Q., Atianand, M.K., Vos, M.R., Schroder, G.F., Fitzgerald, K.A., Wu, H., and Egelman, E.H. (2014). Unified polymerization mechanism for the assembly of ASC-dependent inflammasomes. Cell 156, 1193–1206.

Magiorakos, A.-P., Suetens, C., Monnet, D.L., Gagliotti, C., and Heuer, O.E. (2013). The rise of carbapenem resistance in Europe: just the tip of the iceberg? Antimicrobial resistance and infection control 2, 1–3.

Man, S.M., Karki, R., Malireddi, R.S., Neale, G., Vogel, P., Yamamoto, M., Lamkanfi, M., and Kanneganti, T.-D. (2015). The transcription factor IRF1 and guanylate-binding proteins target activation of the AIM2 inflammasome by Francisella infection. Nat Immunol 16, 467–475.

March, C., Cano, V., Moranta, D., Llobet, E., Perez-Gutierrez, C., Tomas, J.M., Suarez, T., Garmendia, J., and Bengoechea, J.A. (2013). Role of bacterial surface structures on the interaction of Klebsiella pneumoniae with phagocytes. PloS one 8, e56847.

McNab, F.W., Ewbank, J., Howes, A., Moreira-Teixeira, L., Martirosyan, A., Ghilardi, N., Saraiva, M., and O’Garra, A. (2014). Type I IFN induces IL-10 production in an IL-27-independent manner and blocks responsiveness to IFN-gamma for production of IL-12 and bacterial killing in Mycobacterium tuberculosis-infected macrophages. J Immunol 193, 3600–3612.

Motani, K., Kushiyama, H., Imamura, R., Kinoshita, T., Nishiuchi, T., and Suda, T. (2011). Caspase-1 protein induces apoptosis-associated speck-like protein containing a caspase recruitment domain (ASC)-mediated necrosis independently of its catalytic activity. J Biol Chem 286, 33963–33972.

Nassif, X., Fournier, J.M., Arondel, J., and Sansonetti, P.J. (1989). Mucoid phenotype of Klebsiella pneumoniae is a plasmid-encoded virulence factor. Infect Immun 57, 546–552.

O’Neill, L.A., and Bowie, A.G. (2007). The family of five: TIR-domain-containing adaptors in Toll-like receptor signalling. Nat Rev Immunol 7, 353–364.

Ostrowski, P.P., Fairn, G.D., Grinstein, S., and Johnson, D.E. (2016). Cresyl violet: a superior fluorescent lysosomal marker. Traffic 17, 1313–1321.

Pattison, M.J., MacKenzie, K.F., and Arthur, J.S.C. (2012). Inhibition of JAKs in macrophages increases lipopolysaccharide-induced cytokine production by blocking IL-10–mediated feedback. J Immunol 189, 2784–2792.

Penalva, G., Hogberg, L.D., Weist, K., Vlahovic-Palcevski, V., Heuer, O., Monnet, D.L., Group, E.S.-N.S., and Group, E.A.-N.S. (2019). Decreasing and stabilising trends of antimicrobial consumption and resistance in Escherichia coli and Klebsiella pneumoniae in segmented regression analysis, European Union/European Economic Area, 2001 to 2018. Euro Surveill 24.

Peng, J., Yuan, Q., Lin, B., Panneerselvam, P., Wang, X., Luan, X.L., Lim, S.K., Leung, B.P., Ho, B., and Ding, J.L. (2010). SARM inhibits both TRIF-and MyD88-mediated AP-1 activation. Eur J Immunol 40, 1738–1747.

Rathinam, V.A., Jiang, Z., Waggoner, S.N., Sharma, S., Cole, L.E., Waggoner, L., Vanaja, S.K., Monks, B.G., Ganesan, S., Latz, E., et al. (2010). The AIM2 inflammasome is essential for host defense against cytosolic bacteria and DNA viruses. Nat Immunol 11, 395–402.

Regueiro, V., Moranta, D., Frank, C.G., Larrarte, E., Margareto, J., March, C., Garmendia, J., and Bengoechea, J.A. (2011). Klebsiella pneumoniae subverts the activation of inflammatory responses in a NOD1-dependent manner. Cel Microbiol 13, 135–153.

Rusinova, I., Forster, S., Yu, S., Kannan, A., Masse, M., Cumming, H., Chapman, R., and Hertzog, P.J. (2013). Interferome v2.0: an updated database of annotated interferon-regulated genes. Nucleic Acids Res 41, D1040–1046.

Sa-Pessoa, J., Przybyszewska, K., Vasconcelos, F.N., Dumigan, A., Frank, C.G., Hobley, L., and Bengoechea, J.A. (2020). Klebsiella pneumoniae Reduces SUMOylation To Limit Host Defense Responses. mBio 11.

Saraiva, M., and O’Garra, A. (2010). The regulation of IL-10 production by immune cells. Nat Rev Immunol 10, 170–181.

Schlam, D., Bagshaw, R.D., Freeman, S.A., Collins, R.F., Pawson, T., Fairn, G.D., and Grinstein, S. (2015). Phosphoinositide 3-kinase enables phagocytosis of large particles by terminating actin assembly through Rac/Cdc42 GTPase-activating proteins. Nat Comms 6, 8623.

Secombes, C.J., and Zou, J. (2017). Evolution of Interferons and Interferon Receptors. Front Immunol 8, 209.

Sester, D.P., Thygesen, S.J., Sagulenko, V., Vajjhala, P.R., Cridland, J.A., Vitak, N., Chen, K.W., Osborne, G.W., Schroder, K., and Stacey, K.J. (2015). A novel flow cytometric method to assess inflammasome formation. J Immunol 194, 455–462.

Shi, H., Murray, A., and Beutler, B. (2016). Reconstruction of the Mouse Inflammasome System in HEK293T Cells. Bio Protoc 6.

Szretter, K.J., Samuel, M.A., Gilfillan, S., Fuchs, A., Colonna, M., and Diamond, M.S. (2009). The immune adaptor molecule SARM modulates tumor necrosis factor alpha production and microglia activation in the brainstem and restricts West Nile Virus pathogenesis. J Virol 83, 9329–9338.

Taniguchi, K., and Karin, M. (2018). NF-kappaB, inflammation, immunity and cancer: coming of age. Nat Rev Immunol 18, 309–324.

Tomas, A., Lery, L., Regueiro, V., Perez-Gutierrez, C., Martinez, V., Moranta, D., Llobet, E., Gonzalez-Nicolau, M., Insua, J.L., Tomas, J.M., et al. (2015). Functional Genomic Screen Identifies Klebsiella pneumoniae Factors Implicated in Blocking Nuclear Factor kappaB (NF-kappaB) Signaling. J Biol Chem 290, 16678–16697.

Tsuchiya, K., Hara, H., Kawamura, I., Nomura, T., Yamamoto, T., Daim, S., Dewamitta, S.R., Shen, Y., Fang, R., and Mitsuyama, M. (2010). Involvement of absent in melanoma 2 in inflammasome activation in macrophages infected with Listeria monocytogenes. J Immunol 185, 1186–1195.

Uccellini, M.B., Bardina, S.V., Sánchez-Aparicio, M.T., White, K.M., Hou, Y.-J., Lim, J.K., and García-Sastre, A. (2020). Passenger mutations confound phenotypes of SARM1-deficient mice. Cell reports 31, 107498.

Uhlen, M., Oksvold, P., Fagerberg, L., Lundberg, E., Jonasson, K., Forsberg, M., Zwahlen, M., Kampf, C., Wester, K., and Hober, S. (2010). Towards a knowledge-based human protein atlas. Nat Biotech 28, 1248–1250.

Ulland, T.K., Buchan, B.W., Ketterer, M.R., Fernandes-Alnemri, T., Meyerholz, D.K., Apicella, M.A., Alnemri, E.S., Jones, B.D., Nauseef, W.M., and Sutterwala, F.S. (2010). Cutting edge: mutation of Francisella tularensis mviN leads to increased macrophage absent in melanoma 2 inflammasome activation and a loss of virulence. J Immunol 185, 2670–2674.

Wang, P.-H., Ye, Z.-W., Deng, J.-J., Siu, K.-L., Gao, W.-W., Chaudhary, V., Cheng, Y., Fung, S.-Y., Yuen, K.-S., Ho, T.-H., et al. (2018). Inhibition of AIM2 inflammasome activation by a novel transcript isoform of IFI16. EMBO Rep 19, e45737.

Wieland, C.W., van Lieshout, M.H., Hoogendijk, A.J., and van der Poll, T. (2011). Host defence during Klebsiella pneumonia relies on haematopoietic-expressed Toll-like receptors 4 and 2. Eur Respir J 37, 848–857.

Willingham, S.B., Allen, I.C., Bergstralh, D.T., Brickey, W.J., Huang, M.T.-H., Taxman, D.J., Duncan, J.A., and Ting, J.P.-Y. (2009a). NLRP3 (NALP3, Cryopyrin) facilitates in vivo caspase-1 activation, necrosis, and HMGB1 release via inflammasome-dependent and-independent pathways. J Immunol 183, 2008–2015.

Willingham, S.B., Allen, I.C., Bergstralh, D.T., Brickey, W.J., Huang, M.T., Taxman, D.J., Duncan, J.A., and Ting, J.P. (2009b). NLRP3 (NALP3, Cryopyrin) facilitates in vivo caspase-1 activation, necrosis, and HMGB1 release via inflammasome-dependent and -independent pathways. J Immunol 183, 2008–2015.

Xiong, D., Song, L., Geng, S., Jiao, Y., Zhou, X., Song, H., Kang, X., Zhou, Y., Xu, X., and Sun, J. (2019). Salmonella coiled-coil-and TIR-containing TcpS evades the innate immune system and subdues inflammation. Cell reports 28, 804–818. e807.

Xiong, H., Carter, R.A., Leiner, I.M., Tang, Y.W., Chen, L., Kreiswirth, B.N., and Pamer, E.G. (2015). Distinct Contributions of Neutrophils and CCR2+ Monocytes to Pulmonary Clearance of Different Klebsiella pneumoniae Strains. Infect Immun 83, 3418–3427.

Xiong, H., Keith, J.W., Samilo, D.W., Carter, R.A., Leiner, I.M., and Pamer, E.G. (2016). Innate Lymphocyte/Ly6C(hi) Monocyte Crosstalk Promotes Klebsiella Pneumoniae Clearance. Cell 165, 679–689.

Yao, H., Qin, S., Chen, S., Shen, J., and Du, X.D. (2018). Emergence of carbapenem-resistant hypervirulent Klebsiella pneumoniae. Lancet Infectious diseases 18, 25-3099(3017)30628-X.

Ye, P., Rodriguez, F.H., Kanaly, S., Stocking, K.L., Schurr, J., Schwarzenberger, P., Oliver, P., Huang, W., Zhang, P., Zhang, J., et al. (2001). Requirement of interleukin 17 receptor signaling for lung CXC chemokine and granulocyte colony-stimulating factor expression, neutrophil recruitment, and host defense. J Exp Med 194, 519–527.

Yin, Q., Sester, D.P., Tian, Y., Hsiao, Y.S., Lu, A., Cridland, J.A., Sagulenko, V., Thygesen, S.J., Choubey, D., Hornung, V., et al. (2013). Molecular mechanism for p202-mediated specific inhibition of AIM2 inflammasome activation. Cell Rep 4, 327–339.

Zhang, Q., Zmasek, C.M., Cai, X., and Godzik, A. (2011). TIR domain-containing adaptor SARM is a late addition to the ongoing microbe-host dialog. Dev Comp Immunol 35, 461–468.

Zhang, Y., Zeng, J., Liu, W., Zhao, F., Hu, Z., Zhao, C., Wang, Q., Wang, X., Chen, H., Li, H., et al. (2015). Emergence of a hypervirulent carbapenem-resistant Klebsiella pneumoniae isolate from clinical infections in China. J Infect 71, 553–560.

Zhang, Y., Zhao, C., Wang, Q., Wang, X., Chen, H., Li, H., Zhang, F., Li, S., Wang, R., and Wang, H. (2016). High Prevalence of Hypervirulent Klebsiella pneumoniae Infection in China: Geographic Distribution, Clinical Characteristics, and Antimicrobial Resistance. Antimicrob Agents Chemother 60, 6115–6120.

