## Supplementary figures and images for "*Klebsiella pneumoniae* hijacks the Toll-IL-1R protein SARM1 in a type I IFN-dependent manner to antagonize host immunity"

### Supplemental Figure 1

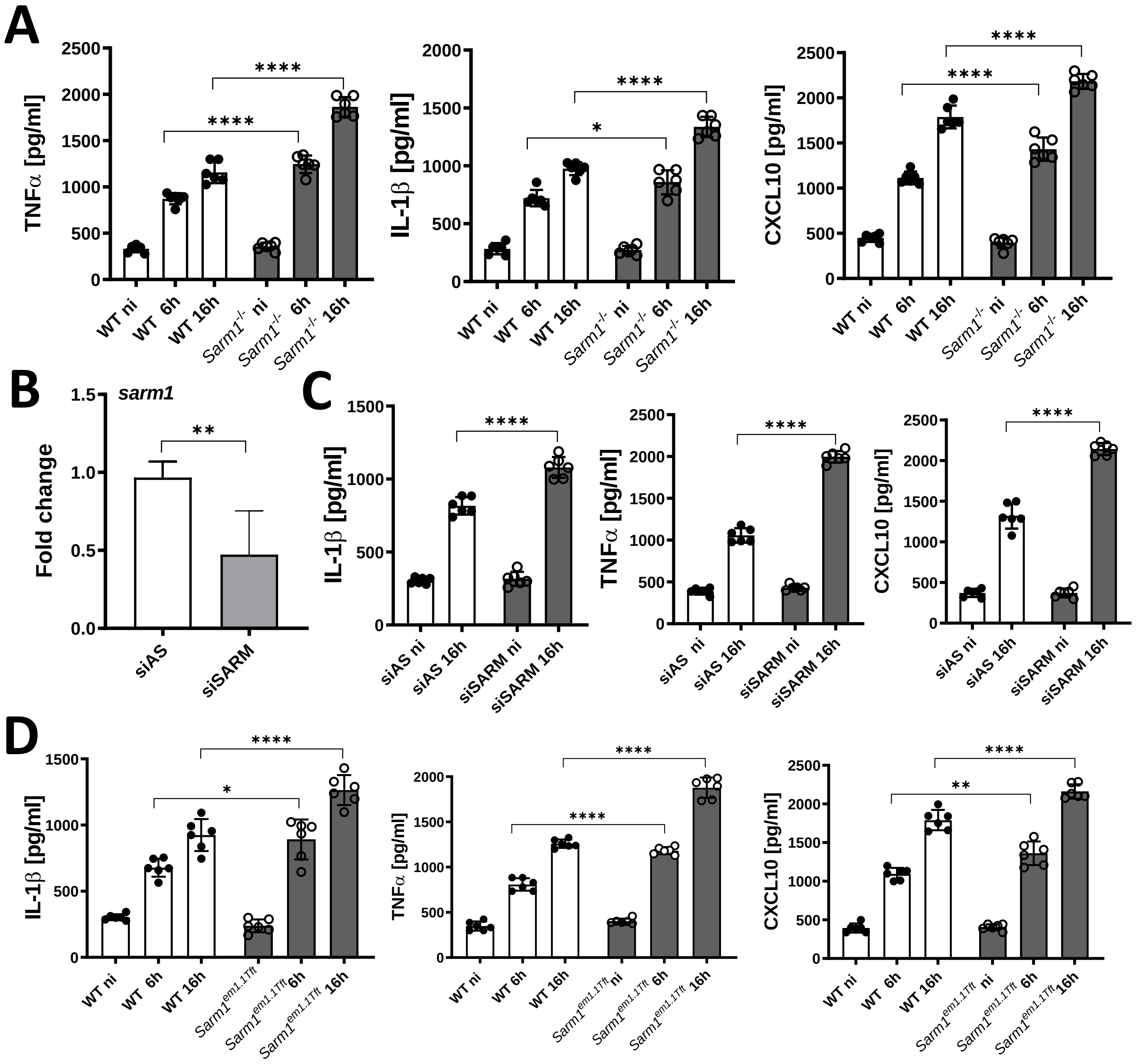

### Supplemental Figure 2

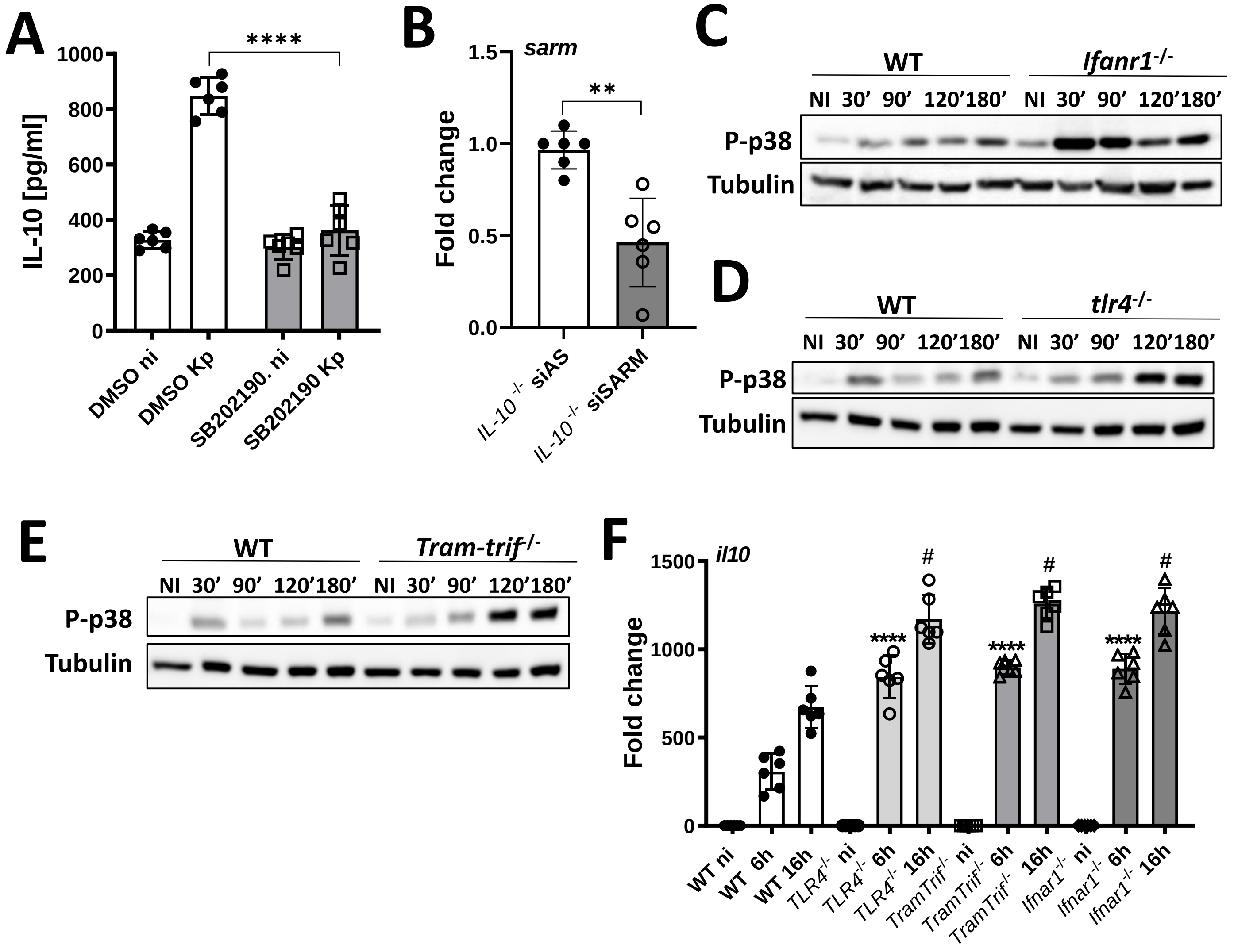

### Supplemental Figure 3

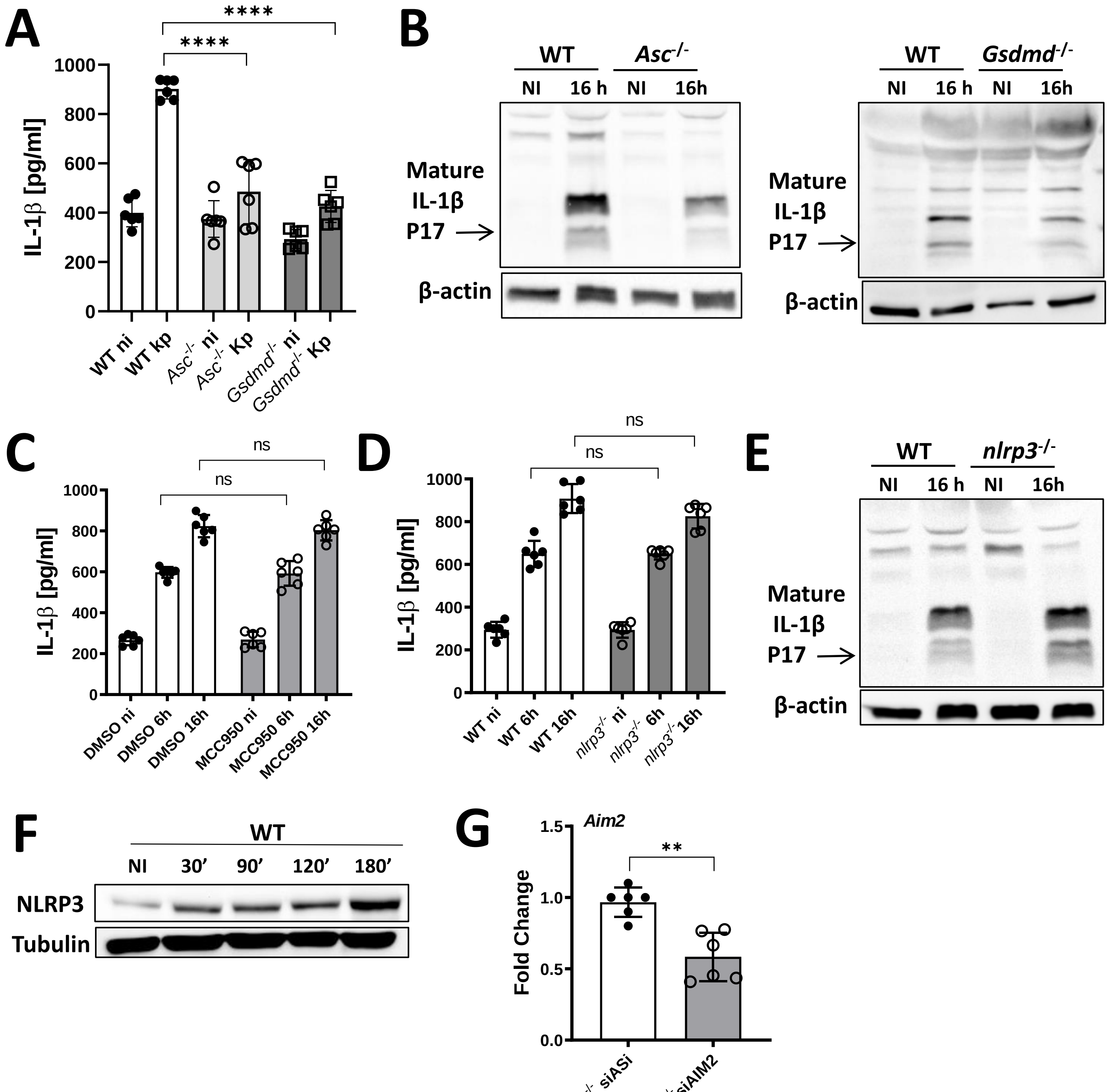

### Supplemental Figure 4

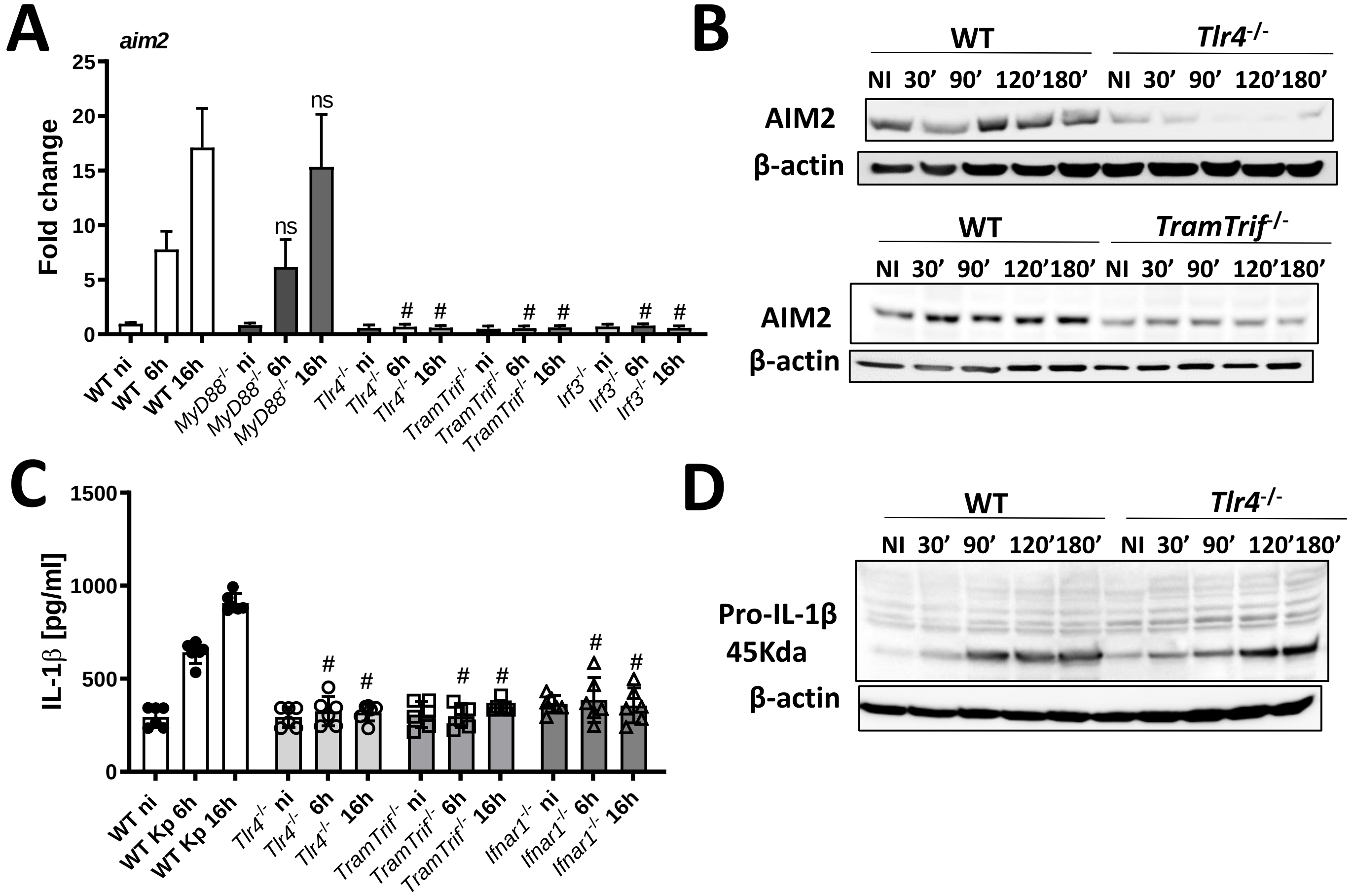

### Supplemental Figure 5

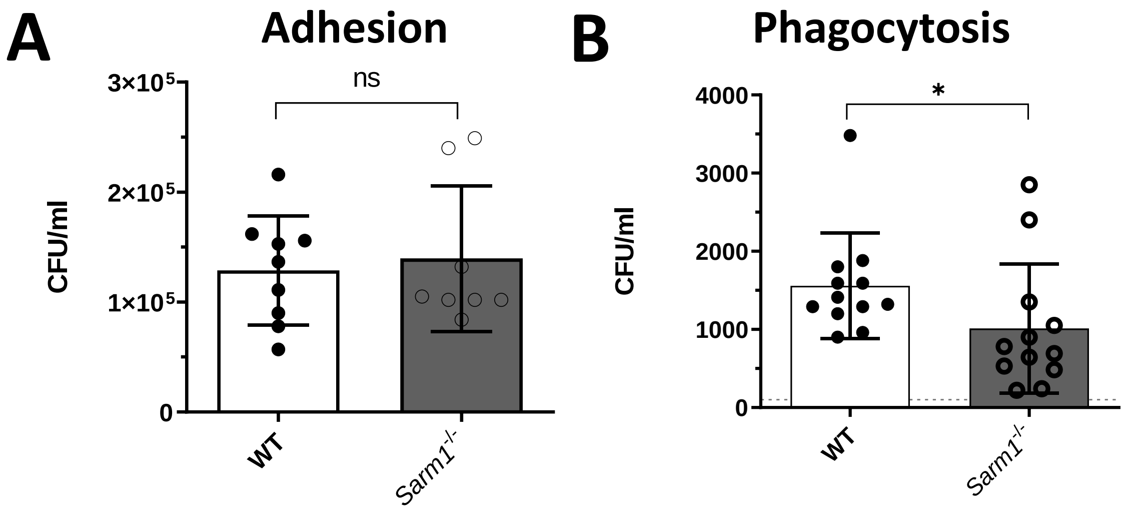

### Supplemental Figure 6

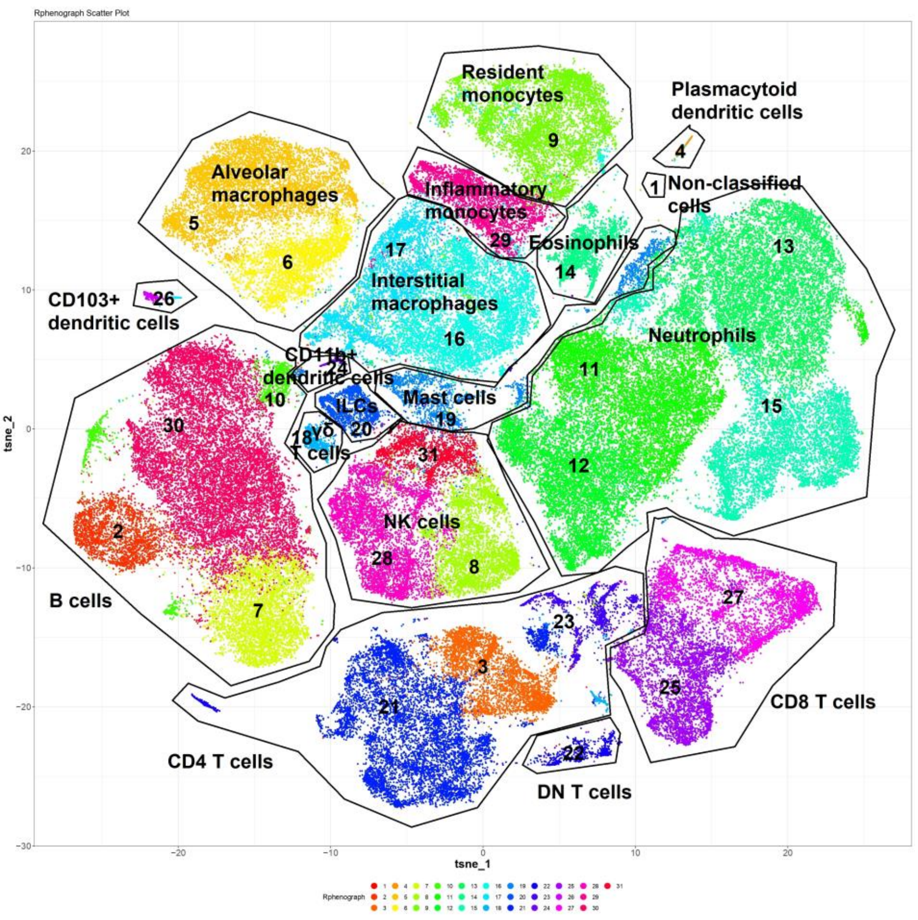

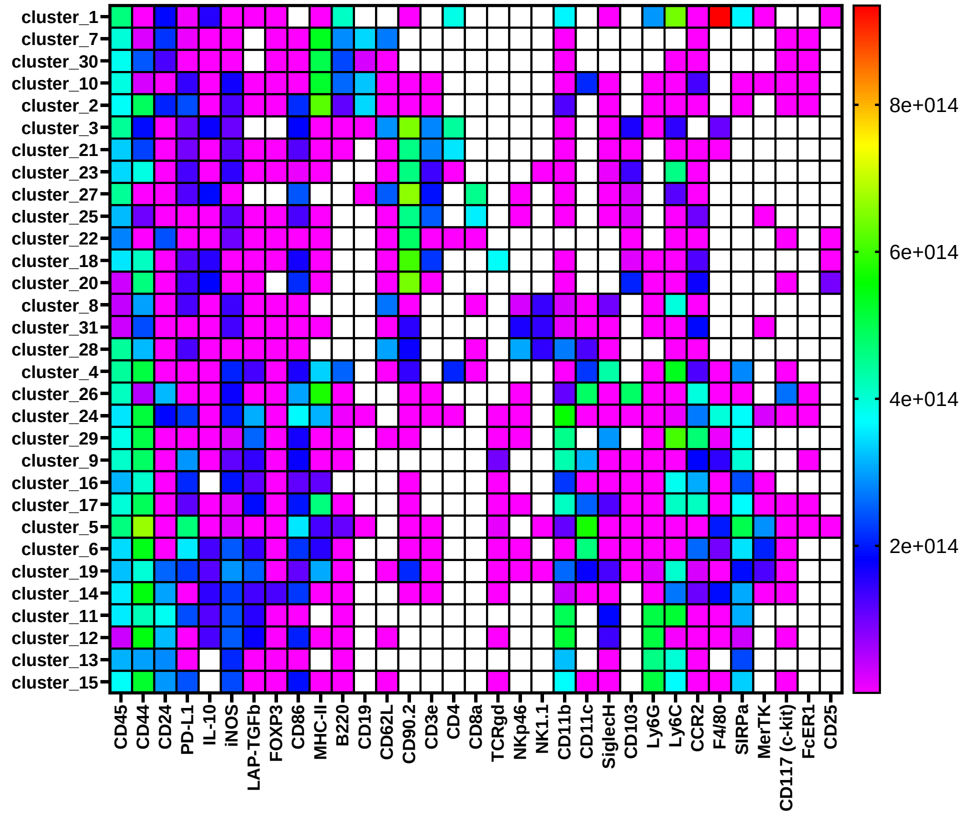

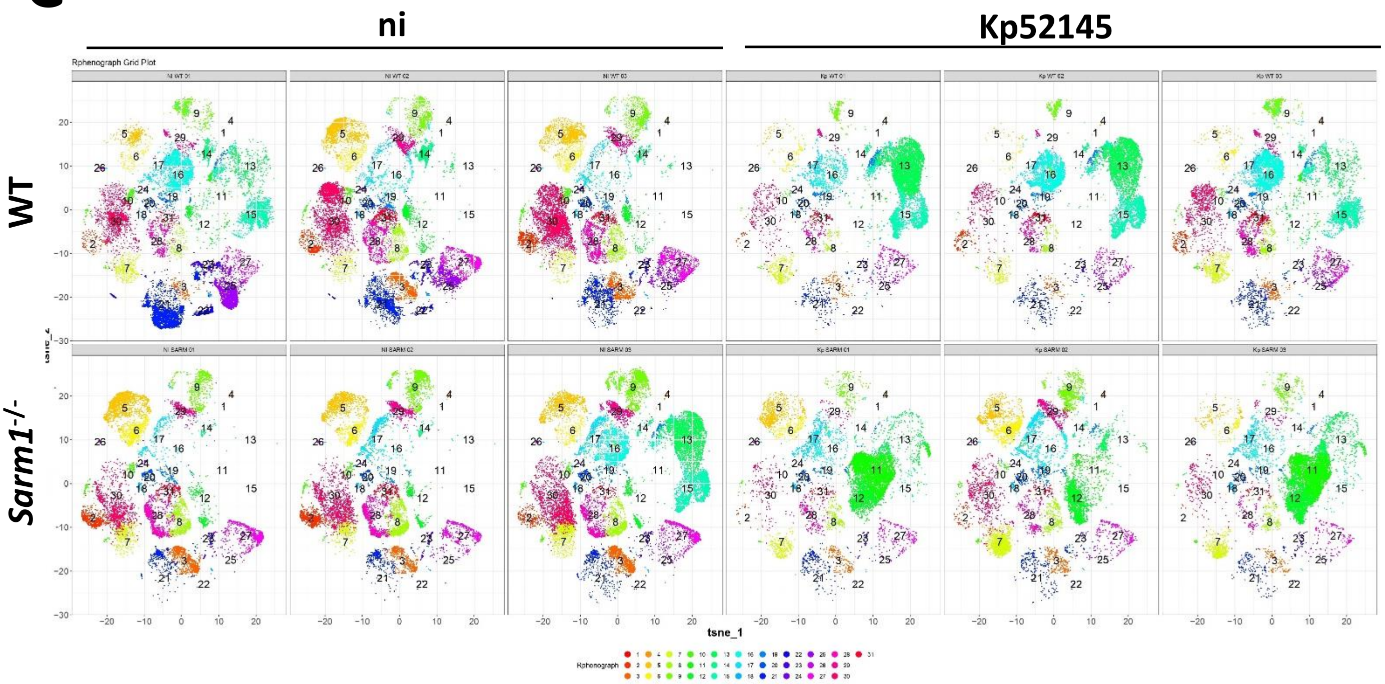
