## Supplemental Table 1 for "*Klebsiella pneumoniae* hijacks the Toll-IL-1R protein SARM1 in a type I IFN-dependent manner to antagonize host immunity"

**Table S1. Antibodies used to characterize immune populations by mass cytometry.**

| **Marker** | **Metal isotope** | **Clone** | **Reference** |
| --- | --- | --- | --- |
| Ly6G | 141Pr | 1A8 | Fluidigm 3141008B |
| SIRPa | 143Nd | P84 | BD 552371* |
| B220 | 144Nd | RA3-6B2 | Fluidigm 3144011B |
| CD4 | 145Nd | RM4-5 | Fluidigm 3145002B |
| F4/80 | 146Nd | BM8 | Fluidigm 3146008B |
| CD45 | 147Sm | 30-F11 | Fluidigm 3147003C |
| CD11b | 148Nd | M1/70 | Fluidigm 3148003C |
| CD19 | 149Sm | 6D5 | Fluidigm 3149002B |
| CD24 | 150Nd | M1/69 | Fluidigm 3150009B |
| CD25 | 151Eu | 3C7 | Fluidigm 3151007B |
| CD3e | 152Sm | 145-2C11 | Fluidigm 3152004B |
| PD-L1 | 153Eu | 10F.9G2 | Fluidigm 3153016B |
| CD103 | 155Gd | 2E7 | BioLegend 121402* |
| CD90.2 | 156Gd | 30-H12 | Fluidigm 3156006B |
| IL-10 | 158Gd | JES5-16E3 | Fluidigm 3158002C |
| TCRgd | 159Tb | GL3 | Fluidigm 3159012B |
| CD62L | 160Gd | MEL-14 | Fluidigm 3160008C |
| iNOS | 161Dy | CXNFT | Fluidigm 3161011B |
| Ly6C | 162Dy | HK1.4 | Fluidigm 3162014B |
| SiglecH | 163Dy | 551 | BioLegend 129602* |
| LAP/TGFb | 164Dy | TW7-16B4 | Fluidigm 3164014B |
| FOXP3 | 165Ho | FJK-16S | Fluidigm 3165024A |
| CCR2 | 166Er | 475301R | R&D MAB55381R* |
| CD335/NKp46 | 167Er | 29A1.4 | Fluidigm 3167008B |
| CD8a | 168Er | 53-6.7 | Fluidigm 3168003B |
| MerTK | 169Tm | *Polyclonal* | R&D AF591* |
| CD161/NK1.1 | 170Er | PK136 | Fluidigm 3170002C |
| CD44 | 171Yb | IM7 | Fluidigm 3171003C |
| CD86 | 172Yb | GL-1 | Fluidigm 3172016B |
| CD117/c-kit | 173Yb | 2B8 | Fluidigm 3173004B |
| MHC-II | 174Yb | M5/114.15.2 | Fluidigm 3174003B |
| FcER1 | 176Yb | MAR-1 | Fluidigm 3176006B |
| CD11c | 209Bi | N418 | Fluidigm 3209005B |

*****Antibodies conjugated to the indicated metal isotype using Maxpar X8 Antibody Labelling Kit.
