## Supplemental Table 2 for "*Klebsiella pneumoniae* hijacks the Toll-IL-1R protein SARM1 in a type I IFN-dependent manner to antagonize host immunity"

**Table S2. Immune populations identified by mass cytometry.**

| **Population** | **Subpopulation** | **Cluster** | **Markers** |
| --- | --- | --- | --- |
| B cells | Naïve B cells | 7 | B220^+^CD19^+^MHC-II^+^CD62L^high^ |
|  | Mature B cells | 30 | B220^+^CD19^+^MHC-II^+^CD62L^low^ |
|  | CD11c^+^ B cells | 10 | B220^+^CD19^+^MHC-II^+^CD62L^low^CD11c^+^ |
|  | Plasma cells | 2 | B220^low^CD19^+^MHC-II^+^ |
| CD4 T cells | Naïve CD4 T cells | 3 | CD90.2^+^CD3^+^CD4^+^CD62L^high^ |
|  | Mature CD4 T cells | 21 | CD90.2^+^CD3^+^CD4^+^CD62L^low^ |
|  | Ly6C^+^ mature CD4 T cells | 23 | CD90.2^+^CD3^+^CD4^+^CD62L^low^Ly6C^high^ |
| CD8 T cells | Naïve CD8 T cells | 27 | CD90.2^+^CD3^+^CD8^+^CD62L^high^ |
|  | Mature CD8 T cells | 25 | CD90.2^+^CD3^+^CD8^+^CD62L^low^ |
| Double negative T cells | | 22 | CD90.2^+^CD3^+^CD4^-/low^CD8^-/low^ |
| Gamma delta T cells | | 18 | CD90.2^+^CD3^+^TCRgd^+^ |
| Innate lymphoid cells | | 20 | CD90.2^+^CD3^-/low^ |
| NK cells | CD90.2^-^ NK cells | 8 | CD90.2^-/low^NK1.1^+^NKp46^+^ |
|  | CD90.2^+^CD62L^-^ NK cells | 31 | CD90.2^+^NK1.1^+^NKp46^+^CD62L^-/low^ |
|  | CD90.2^+^CD62L^+^ NK cells | 28 | CD90.2^+^NK1.1^+^NKp46^+^CD62L^+^ |
| Plasmacytoid dendritic cells | | 4 | MHC-II^+^CD11c^+^B220^+^SiglecH^+^F4/80^-/low^ |
| Myeloid dendritic cells | CD103+ dendritic cells | 26 | MHC-II^+^CD11c^+^CD11b^low^CD103^+^F4/80^-/low^ |
|  | CD11b+ dendritic cells | 24 | MHC-II^+^CD11c^low^CD11b^+^CD103^-^F4/80^+^ |
| Inflammatory monocytes | | 29 | MHC-II^-^Ly6G^-^Ly6C^+^CD11b^+^CD11c^-^CCR2^high^ |
| Resident monocytes | | 9 | MHC-II^-^Ly6G^-^Ly6C^-^CD11b^+^CD11c^+^CCR2^+^ |
| Interstitial macrophages | CD11c^-^ interstitial macrophages | 16 | MHC-II^+^Ly6G^-^Ly6C^+^CD11b^+^CD11c^-/low^ |
|  | CD11c^+^ interstitial macrophages | 17 | MHC-II^+^Ly6G^-^Ly6C^+^CD11b^+^CD11c^+^ |
| Alveolar macrophages | CCR2^-^ alveolar macrophages | 5 | MHC-II^+^Ly6G^-^Ly6C^-^CD11b^low^CD11c^+^CCR2^-^ |
|  | CCR2^+^ alveolar macrophages | 6 | MHC-II^+^Ly6G^-^Ly6C^-^CD11b^-^CD11c^+^CCR2^+^ |
| Mast cells/basophils | | 19 | MHC-II^+^Ly6G^-^Ly6C^+^F4/80^-/low^CD90.2^+^CD11b^+^ CD11c^+^ |
| Eosinophils | | 14 | MHC-II^-^Ly6G^-^Ly6C^+^F4/80^+^ |
| Neutrophils | SiglecH^high^PD-L1^+^ neutrophils | 11 | MHC-II^-^Ly6G^+^Ly6C^+^F4/80^-/low^SiglecH^high^PD-L1^high^ |
|  | SiglecH^high^PD-L1^-^ neutrophils | 12 | MHC-II^-^Ly6G^+^Ly6C^+^F4/80^-/low^SiglecH^high^PD-L1^low^ |
|  | SiglecH^low^PD-L1^-^ neutrophils | 13 | MHC-II^-^Ly6G^+^Ly6C^+^F4/80^-/low^SiglecH^low^PD-L1^low^ |
|  | SiglecH^low^PD-L1^+^ neutrophils | 15 | MHC-II^-^Ly6G^+^Ly6C^+^F4/80^-/low^SiglecH^low^PD-L1^high^ |
