## Supplemental Table 3 for "*Klebsiella pneumoniae* hijacks the Toll-IL-1R protein SARM1 in a type I IFN-dependent manner to antagonize host immunity"

**Table S3. Primers used in this study.**

| **Name** | **Sequence (5’-3’)** |
| --- | --- |
| mSARM Forward | GGT GCA CAA GGA GAT TGT GAC |
| mSARM Reverse | CAT GGG ACC ATT TGA TGC CGT T |
| mIL1B-F1 | AGA TGA AGG GCT GCT TCC AAA |
| mIL1B-R1 | AAT GGG AAC GTC ACA GAC CA |
| mTNFa-F1 | TTC TGT CTA CTG AAC TTC GGG GTG ATC GGT CC |
| mTNFa-R1 | GTA TGA GAT AGC AAA TCG GCT GAC GGT GTG GG |
| mIFNb-F | ATG GTG GTC CGA GCA GAG AT |
| mIFNb-R | CCA CCA CTC ATT CTG AGG |
| mCXCL10-F1 | AGT GCT GCC GTC ATT TTC TG |
| mCXCL10-R1 | ATT CTC ACT GGC CCG TCA T |
| mISG15-F | GGG GCC ACA GCA ACA TCT AT |
| mISG15-R | CGC TGG GAC ACC TTC TTC TT |
| m.Mx1_F1 | GAC TAC CAC TGA GAT GAC CCA GC |
| m.Mx1_R1 | ATT TCC TCC CCA AAT GTT TTC A |
| mIFIT1-F | CAG GTT TCT GAG GAG TTC TG |
| mIFIT1-R | TGA AGC AGA TTC TCC ATG AC |
| mIL10_F1 | GGA CTT TAA GGG TTA CTT GGG TTG CC |
| mIL10_R1 | CAT GTA TGC TTC TAT GCA GTT GAT GA |
| mIL12_p40_F1 | GGA AGC ACG GCA GCA GAA TA |
| mIL12_p40_R1 | AAC TTG AGG GAG AAG TAG GAA TGG |
| AIM2_Fwd | GTT GAA TCT AAC CAC GAA GTC C |
| AIM2_Rvr | CTA CAA GGT CCA GAT TTC AAC TG |
